# Topographic organization of the human subcortex unveiled with functional connectivity gradients

**DOI:** 10.1101/2020.01.13.903542

**Authors:** Ye Tian, Daniel S. Margulies, Michael Breakspear, Andrew Zalesky

## Abstract

Understanding the topographic organization of the human brain remains a major goal in neuroscience. Brain atlases are fundamental to this goal, yet many contemporary human atlases cover only the cerebral cortex, leaving the subcortex a *terra incognita*. We revealed the complex topographic organization of the human subcortex by disambiguating smooth connectivity gradients from discrete areal boundaries in resting-state fMRI data acquired from more than 1000 healthy adults. This unveiled four scales of subcortical organization, recapitulating well-known anatomical nuclei at the coarsest scale and delineating 27 new bilateral regions at the finest. Ultra-high field strength fMRI corroborated and extended this organizational structure, enabling delineation of finer subdivisions of hippocampus and amygdala, while task-evoked fMRI revealed a subtle reorganization of subcortical topography in response to changing cognitive demands. A new subcortical atlas was delineated, personalized to account for individual connectivity differences and utilized to uncover reproducible relationships between subcortical connectivity and individual variation in human behaviors. Linking cortical networks to subcortical regions recapitulated a task-positive to task-negative organizational axis. The new atlas enables holistic connectome mapping and characterization of cortico-subcortical connectivity.

## Introduction

The ubiquity of magnetic resonance imaging (MRI) has enabled high-throughput mapping of standardized brain parcellation atlases that represent consensus cortical boundaries across hundreds of scanned people^1^. Compared to the cerebral cortex, efforts to parcellate the human subcortex are scarcer. Indeed, many leading brain atlases delineate only cortical regions^2-5^.

The subcortex is widely implicated in the pathophysiology of many neuropsychiatric disorders and its role in cognition is unequivocal^6^. To better elucidate the role of the subcortex in health and disease, further work is needed to advance recent efforts to map the functional architecture of the subcortex^7, 8^. This study aims to unravel the incredibly complex functional organization of the human subcortex by developing and validating a comprehensive subcortical atlas using 3 and 7 Tesla functional MRI (fMRI).

Cortical boundaries^9^ within a brain atlas are often delineated at locations of abrupt spatial change in cortical microstructure, function and/or connectivity^2, 3, 5, 10^. Cortical gradients^11^ and the related connectopy concept^12^ offer a complementary representation of cortical topography. Rather than delineating discrete spatial boundaries, topographic change is represented continuously along overlapping organizational axes^13^.

Brain cartography studies tend to espouse one or the other of these two representations^1^. This is exemplified by current contention about whether the cerebellum’s functional topography is best represented by continuous gradients^14^ or discrete boundaries^15^. However, the two paradigms are not necessarily contradictory. Spatial variation across expanses of the cerebrum may be evident, but too gradual to warrant boundary delineation, and thus a continuum might be deemed to provide the most parsimonious representation. This leads to a question of model selection: How sharp and conspicuous must a change be to warrant boundary delineation^16^? Here, we mapped functional connectivity gradients of the human subcortex and delineated subcortical boundaries only when justified by a formal model selection process.

Subcortical geometry is fundamentally different to the convoluted sheet-like structure of the cerebral cortex^17^. New paradigms are thus required to facilitate subcortical cartography. To this end, we developed gradientography, an fMRI analogue of diffusion MRI tractography, which enables quantification of subcortical connectivity gradients and a statistically principled formalism to guide boundary delineation. This allowed us to comprehensively parcellate the subcortex based on resting-state functional connectivity profiles and produce a standardized atlas representing consensus boundaries among more than 1000 healthy adults. The new atlas comprises four scales, recapitulating well-known anatomical nuclei at the coarsest scale and revealing 27 bilateral regions at the finest. The new atlas was replicated using ultra-high field strength fMRI (7T) and personalized to account for individual connectivity differences. Analysis of task-evoked fMRI revealed a subtle reorganization of subcortical topography in response to changing cognitive demands. We used the new atlas to uncover correlational relationships between subcortical functional connectivity and individual variation in five behavioral domains, and to reveal a new task-positive to task-negative organizational axis in cortico-subcortical connectivity^18^. This study provides an important step toward understanding the functional organization of the human subcortex.

## Results

We investigated the functional connectivity architecture of the human subcortex using resting-state fMRI acquired at 3T (n=1080) and 7T (n=183), and task-evoked fMRI acquired at 3T (n=725) from healthy adults participating in the Human Connectome Project (HCP)^19^. Demographics are shown in Supplementary Table S1. Two 3T resting-state fMRI sessions were acquired for each individual, with the first session used as the primary dataset (REST1) and the second used for replication (REST2). Gradients in functional connectivity were mapped using Laplacian eigenmaps and a newly developed procedure called gradientography. Formal model selection was used to determine whether gradient magnitude peaks were sufficiently large to warrant boundary delineation. The competing model posited a gradual spatial change in connectivity, without discrete boundaries. This resulted in a new fMRI-based multiscale parcellation atlas of the subcortex, which we used to reveal new insights into subcortical function and architecture.

### Connectivity gradients and gradientography

Whole-brain functional connectivity fingerprints^20^ were mapped for each subcortical voxel and individual. Spatial gradients characterizing continuous modes of functional connectivity variation across the topography of the subcortex were computed using Laplacian eigenmaps and the connectopy method^12^ (Fig 1a). The eigenmap, or *connectopy*, explaining the most variance in functional connectivity was called Gradient I (Fig 1b), and the next two successive eigenmaps were called Gradient II and III, respectively (Supplementary Fig S1). Further gradients explained substantially less variance and were thus not considered (Supplementary Fig S2). A new technique called gradientography was developed to visualize the connectivity gradients. Gradientography involves estimating the principal gradient direction for each subcortical voxel, representing gradient directions and magnitudes with tensors, and then propagating streamlines through the tensor field using established tools for diffusion MRI tractography. In this way, approximately 15,000 streamlines were initialized and propagated throughout the entire subcortical volume to map the principal organizational axes of the subcortex for Gradient I (Fig 1b), and Gradient II and III (Supplementary Fig S1).

**Fig 1.**
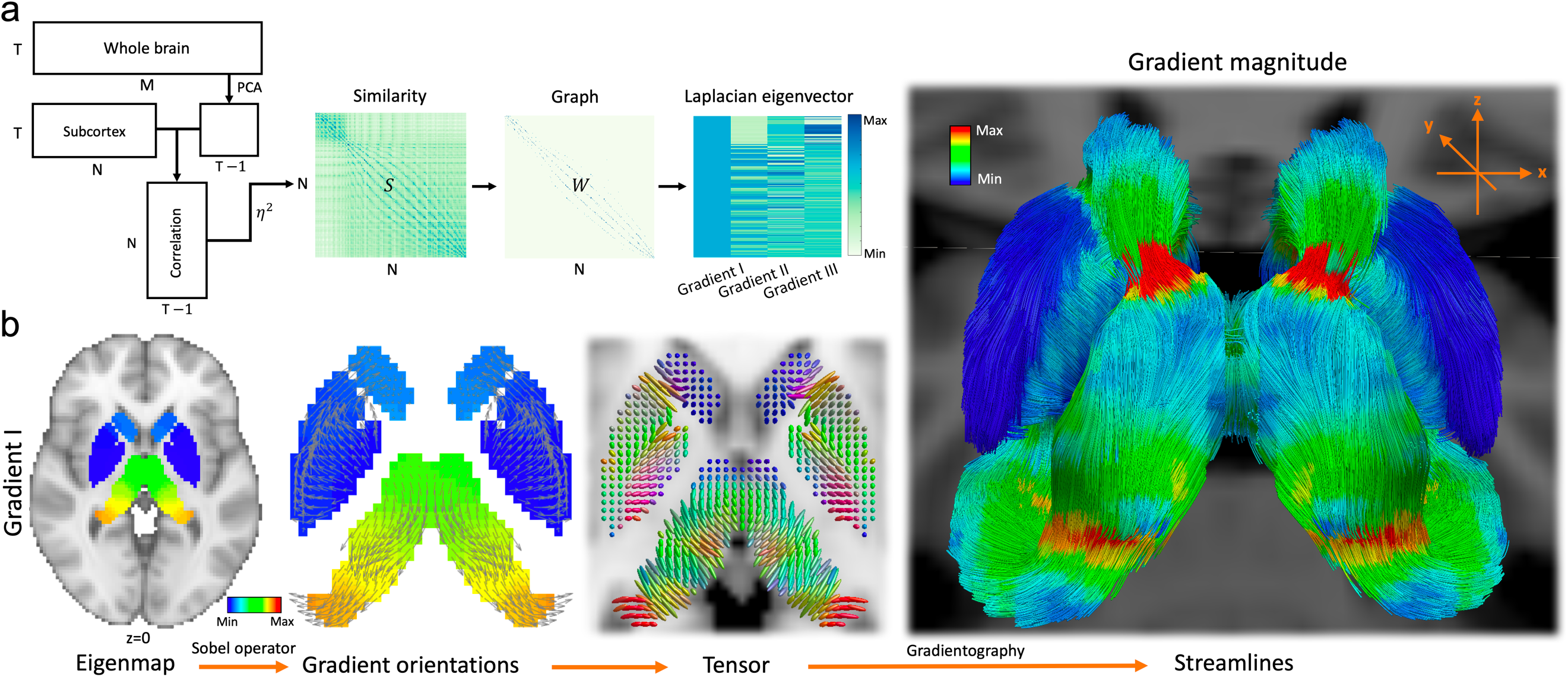
Gradientography in the human subcortex. **a**, Schematic of functional connectivity gradient mapping. Similarity in connectional fingerprints between pairs of subcortical voxels was measured using the *η*_2_ coefficient. Pairs of voxels with similar fingerprints connected to similar brain areas. The similarity matrix (*S*) shown represents a group consensus for 1080 healthy adults. The similarity matrix was transformed into a sparse graph with adjacency matrix *W*. Graph nodes correspond to subcortical voxels. Eigenvalues and eigenvectors were computed for the graph Laplacian. The eigenvectors with the second, third and fourth smallest eigenvalue are called Gradient I, II and III respectively. Each gradient provides a mapping of continuous variation in functional connectivity across the topography of the subcortex. ***N***, number of subcortical voxels; ***M***, number of whole-brain gray matter voxels; *T*, number of time frames. **b**, Gradientography pipeline. Gradients were projected to subcortical voxels to enable anatomical visualization. The axial slice shown is for Gradient I. The Sobel operator was used to estimate the local gradient direction and magnitude for each subcortical voxel. The arrows shown point in the direction of the estimated gradients. Arrow lengths are normalized to unit length. Tensors were fitted to the gradient field. Tensors are colored according to gradient direction (red: left-right, green: posterior-anterior, blue: superior-inferior). Streamlines were propagated through the tensor field using methods for diffusion MRI tractography. Streamlines are shown from a posterior viewpoint and colored according to gradient magnitude. The red bands indicate peaks in the gradient magnitude that demarcate hippocampus-thalamus and caudate-thalamus boundaries.

Gradient magnitude peaks indicated locations of putative functional boundaries between regions. To visualize peak locations, streamlines were colored according to gradient magnitude. Peak locations appear as circumscribed red bands (Fig 1b) and recapitulate boundaries between well-known anatomical nuclei. Local maxima in the gradient magnitude images showed moderate consistency among the three gradients (Supplementary Fig S1 & S3), and thus only Gradient I was used for boundary delineation. Gradient I streamlines were partitioned into dorsal and ventral groups at the location of the anatomical discontinuity between caudate tail and anterior extent of thalamus. The dorsal group covered globus pallidus (GP) and striatum (Fig 2a), whereas amygdala, hippocampus and thalamus comprised the ventral group (Fig 2b).

**Fig 2.**
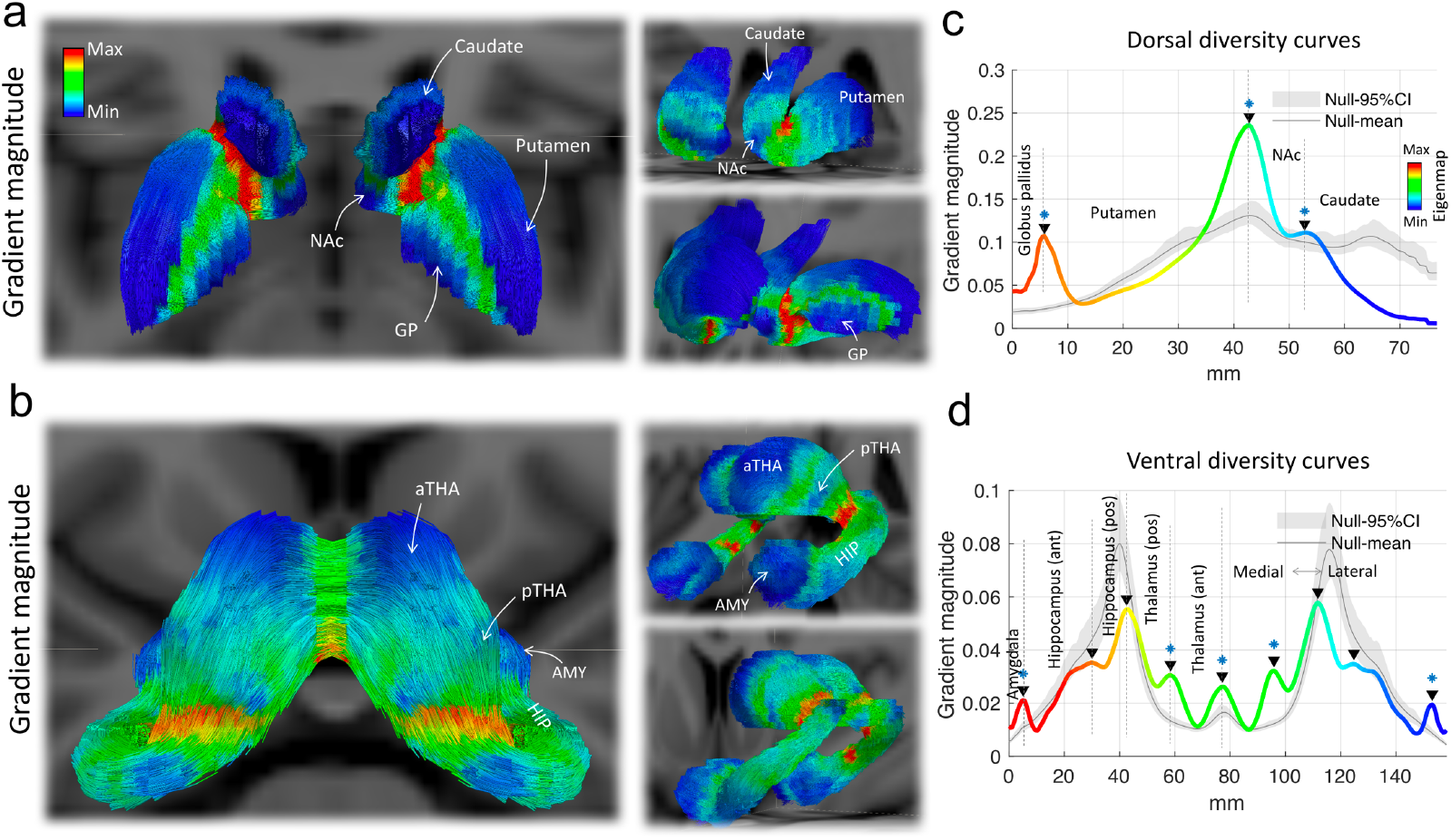
Model selection: Which gradient magnitude peaks are sufficiently large to warrant boundary delineation? Streamlines were partitioned into bilaterally symmetric dorsal (**a**) and ventral (**b**) groups. The streamlines shown are colored according to gradient magnitude. Dorsal streamlines traverse globus pallidus (GP), putamen, nucleus accumbens (NAc) and caudate. Ventral streamlines traverse left and right thalamus (THA), hippocampus (HIP) and amygdala (AMY). Dorsal (**c**) and ventral (**d**) diversity curves were mapped to parameterize gradient magnitude as a function of spatial position along the length of dorsal and ventral streamline trajectories. Black triangles indicate diversity curve peaks (local maxima), demarcating locations of putative functional boundaries. Diversity curves are colored according to eigenmap values for Gradient I. Null diversity curves for the null hypothesis of an absence of discrete boundaries are shown in gray, with shading used to show 95% confidence intervals. Peaks in the null diversity curves are due to the effects of geometry. Peaks for which the null hypothesis could be rejected are demarcated with blue asterisks (*p*<0.05). aTHA, anterior thalamus; pTHA, posterior thalamus; mm, millimeter.

We next sought to quantify the size of each gradient magnitude peak to determine whether gradients were sufficiently abrupt to warrant boundary delineation. To this end, gradient magnitude images were projected onto two-dimensional trajectories called diversity curves^16^. A separate diversity curve was mapped for the dorsal and ventral streamline group. The dorsal diversity curve comprised three distinct local maxima indicating the locations of putative boundaries between GP-putamen, putamen-nucleus accumbens (NAc) and NAc-caudate (Fig 2c). Nine local maxima were evident in the ventral diversity curve, suggesting boundary locations that distinguished amygdala, anterior and posterior hippocampus, anterior and posterior thalamus, as well as left and right thalamus (Fig 2d). Locations of gradient magnitude peaks could not be explained by structural MRI signal properties and potential confounds, including the fMRI signal-to-noise ratio, fMRI signal magnitude, T1-weighted and T1/T2-weighted contrast (Supplementary Fig S4).

### A new subcortical atlas

The magnitude of local maxima in the diversity curves varied by a factor of more than ten, suggesting that putative boundaries between some regions were markedly more abrupt than others. How large should a peak be to warrant boundary delineation? Small peaks can emerge due to the convoluted geometry of the subcortex and other confounds (Supplementary Fig S5). False positive boundaries should not be delineated at such peaks. Therefore, for each peak, a formal model selection procedure was used to test the null hypothesis that peak magnitude could be explained by geometry and/or other confounds. Rejection of the null hypothesis indicated that boundary delineation was warranted. Otherwise, if the peak magnitude was not sufficiently large to enable rejection of the null hypothesis, a continuous representation of spatial variation in connectivity was deemed to be the most parsimonious model^16^. The null hypothesis was tested using a geometry-preserving null model (see Methods). Diversity curves generated with the null model are colored gray in Fig 2c, d. The actual peaks significantly exceeded all null data (*p*<0.05, false discovery rate corrected for all 12 peaks), except for peaks separating bilateral hippocampus/thalamus and anterior-posterior hippocampus. However, given that hippocampus and thalamus are anatomically and functionally distinct, and the peak separating these two regions was unequivocal, we made an exception and delineated a boundary here, whereas the putative anterior-posterior hippocampus boundary was not delineated, consistent with the null data. While peaks in the null diversity curves often correspond in locations to peaks in the actual diversity curves, including the functional boundaries between putamen/NAc, hippocampus (pos)/thalamus (pos) and left/right thalamus, distinct peaks are also evident where the effect of geometry is minimal (Supplementary Fig S5a), such as amygdala/hippocampus (ant), thalamus (pos)/thalamus (ant) and GP/putamen. This indicates that subcortical geometry can predict and account for the location of some functional boundaries, but geometry *per se* cannot explain the magnitude of all boundaries, except for the hippocampus/thalamus (see discussion in Supplementary S1). Finally, the watershed transform algorithm was used to segment subcortical voxels into contiguous regions, such that boundaries demarcated watershed lines.

This resulted in 8 bilateral regions that constituted a relatively coarse atlas, referred to as the Scale I atlas (Fig 3a). The Scale I atlas recapitulates well-known anatomical nuclei; namely, amygdala, hippocampus, thalamus, GP, NAc, caudate and putamen as well as an anteroposterior thalamic partition, which corresponds to the spatial distribution of the two sub-populations of thalamic projection cells (matrix and core)^21^. Gradient mapping, boundary delineation and model selection were then iteratively performed separately for each region comprising the Scale I atlas. This resulted in a finer atlas comprising 16 bilateral regions, called the Scale II atlas. In turn, the null hypothesis was independently tested for each region in the Scale II atlas. Continuing this process recursively until the null hypothesis could not be rejected for any regions yielded four progressively more detailed atlases of the subcortex (Scale I: 16 regions, Fig 3a; II: 32 regions; III: 50 regions; IV: 54 regions, Fig 3b, Supplementary Fig S6).The new atlas is organized hierarchically, with the entire subcortex residing at the base of a hierarchical tree and the smallest regions comprising the Scale IV atlas defining the leaves (Fig 4a, b). Subcortical gradients were also hierarchically organized (Fig 4b, c & Supplementary Fig S7). A schematic of the recursive procedure is provided in Supplementary Fig S8.

**Fig 3.**
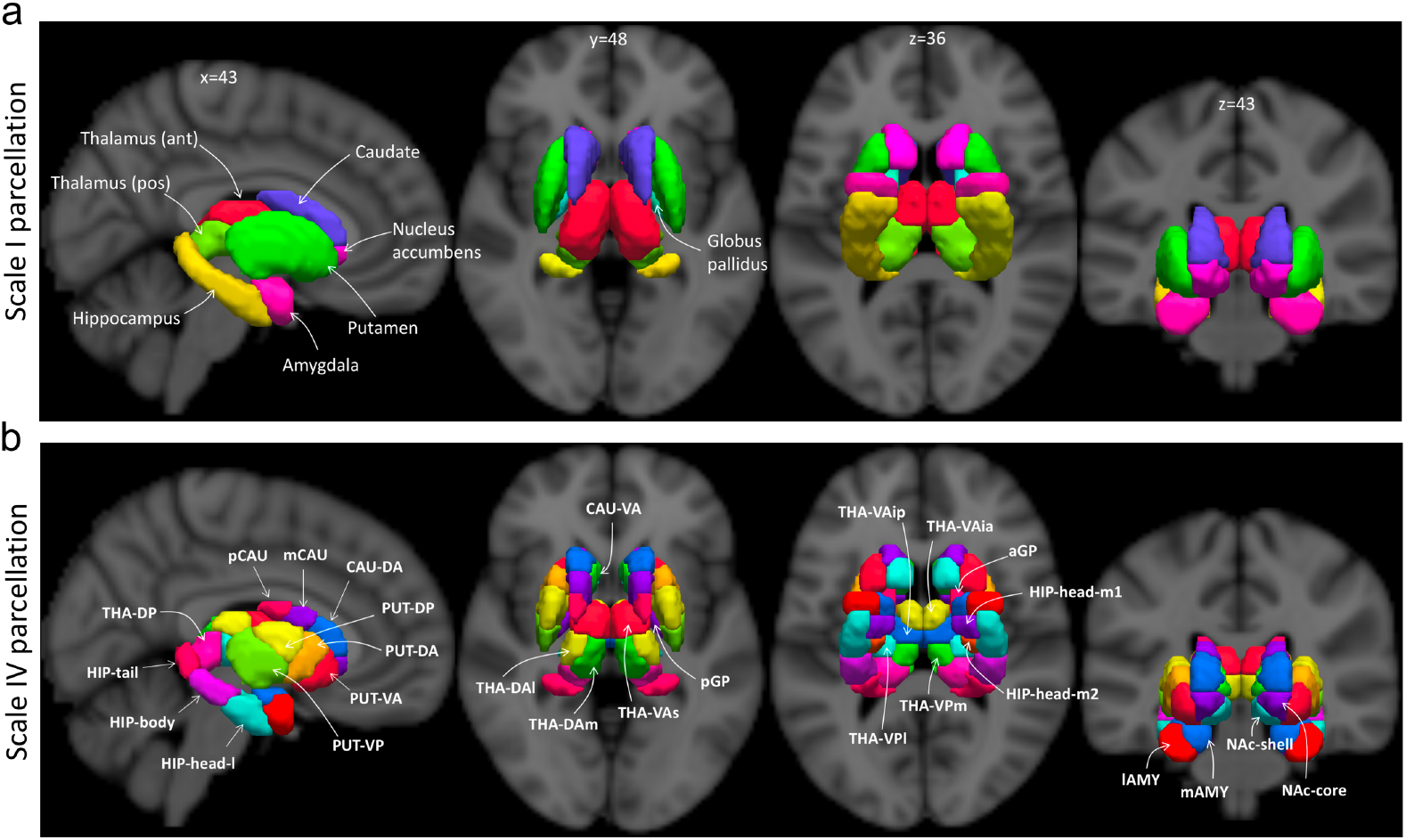
Multiscale group-consensus parcellation atlas derived from 3 Tesla resting-state functional MRI acquired in 1080 healthy adults. **a**, Scale I is the coarsest atlas and comprises 8 bilateral regions that recapitulate anatomical nuclei, including caudate nucleus, nucleus accumbens, putamen, amygdala, hippocampus and thalamus. An anteroposterior division within the thalamus is also present. **b**, Scale IV is the finest scale of the parcellation hierarchy and comprises 27 bilateral regions. Homologous regions are colored identically. A nomenclature for all regions is provided in Supplementary Table S2. PUT, putamen; NAc, nucleus accumbens; CAU, caudate nucleus; GP, globus pallidus; HIP, hippocampus; AMY, amygdala; THA, thalamus; a, anterior; p, posterior; v, ventral; d, dorsal; m, medial; l, lateral; s, superior; i, inferior.

**Fig 4.**
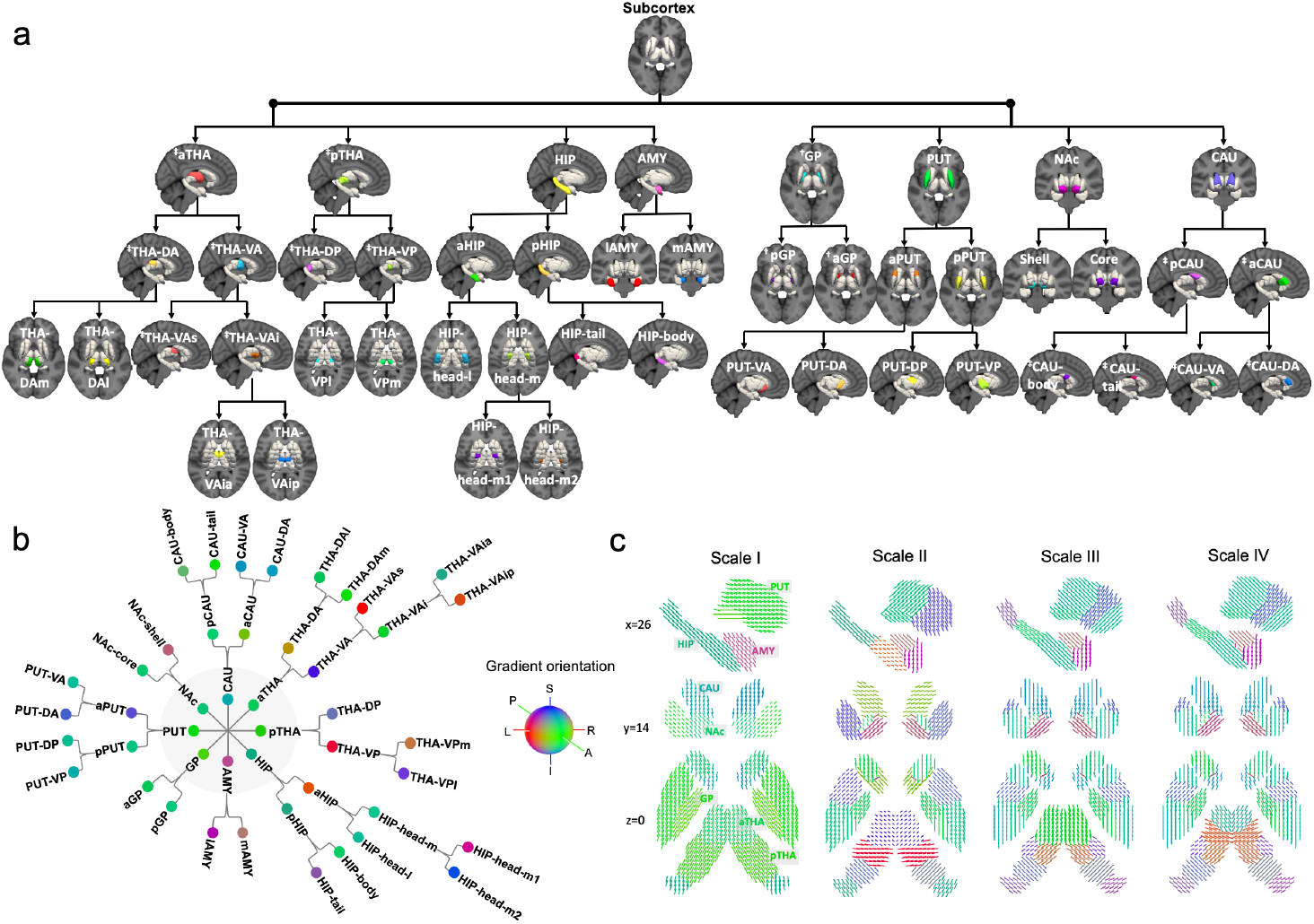
Organization of the human subcortex. Model selection and null hypothesis testing was used to determine whether gradient magnitude peaks were sufficiently large to warrant boundary delineation. This process was repeated recursively for each new region delineated, unveiling a multiscale parcellation architecture. The null hypothesis of an absence of boundaries could not be rejected for any of the regions comprising Scale IV, and thus further (finer) parcellation was not warranted. **a**, Top-down representation of the parcellation hierarchy. Regions are shown atop a reference structural MRI image. Homologous regions are colored identically. **b**, Parcellation hierarchy visualized in the form of a circular tree. The central node represents the entire subcortex and other nodes represent distinct regions. Regions are arranged within four concentric circles, where the innermost circle (gray) is the first level (Scale I). Nodes are colored according to principal gradient orientations within each region (red: left-right, green: posterior-anterior, blue: superior-inferior). The principal gradient orientation was estimated by averaging local gradient directions across all voxels within each region. **c**, Region-averaged gradient directions represented with quiver lines for Scale I-IV. The same images without region averaging are shown in Supplementary Fig S7. A nomenclature for all regions is provided in Supplementary Table S2. PUT, putamen; NAc, nucleus accumbens; CAU, caudate nucleus; GP, globus pallidus; HIP, hippocampus; AMY, amygdala; THA, thalamus; a, anterior; p, posterior; v, ventral; d, dorsal; m, medial; l, lateral. †, caudate removed for visualization purposes; ‡, putamen and GP removed for visualization purposes.

Parcellation of amygdala terminated at Scale II, striatal parcellation continued until Scale III, while only thalamus and hippocampus progressed to Scale IV. A nomenclature for all delineated regions was developed (Supplementary Table S2). In particular, regions that could be unequivocally matched to homologous anatomical structures were labelled as such, while the remaining regions were labelled with reference to standard anatomical planes (e.g. anterior-posterior, medial-lateral, dorsal-ventral and superior-inferior).

Scales I and II generally recapitulate anatomical boundaries, whereas some divergence emerges at finer scales between anatomy and the boundaries delineated here based on functional connectivity. The 8 thalamic subregions comprising Scale IV showed moderate spatial correspondence with histologically-delineated thalamic nuclei (Supplementary Fig S9a). The hippocampus was segmented along its long axis into anterior (head) and posterior (body and tail) components (Supplementary Fig S9b), consistent with previous function-based hippocampal parcellations^22, 23^. Although the head of hippocampus was further divided into medial and lateral components, the overall long-axis organization diverged from the characteristic mediolateral organization that is found histologically^24^.

Connectivity gradients (Gradient I-III) for hippocampus, striatum and thalamus were projected onto the cortical surface based on the cortical vertices with which they were most strongly connected^12, 25^ (Supplementary Fig S10).

### Parcellation homogeneity

Having delineated a new subcortical atlas, we next sought to investigate the atlas’s validity and reproducibility. A key marker of parcellation validity is within-parcel homogeneity of functional signals^1^. Homogeneity was measured based on within-parcel synchrony of fMRI time series acquired in the second fMRI session (REST2), whereas the first session (REST1) was used for parcellation. Homogeneity was computed separately for each region and then averaged across all regions to yield parcellation homogeneity estimates for Scale I-IV (Supplementary Fig S11a).

As expected^3^, homogeneity increased monotonically from Scale I to IV (Fig 5a). To test whether parcellation homogeneity was significantly greater than expected due to chance, homogeneity was also estimated for ensembles of random subcortical parcellations (Supplementary Fig S11b). Random parcellations were generated by delineating boundaries at arbitrary locations, subject to preservation of key geometric properties (see Methods). Scales II-IV were significantly more homogeneous than expected due to chance (*p*<0.01), whereas this was not the case for Scale I (*p*=0.23, Fig 5a). Scale I regions (Fig 3a) encompassed multiple functionally distinct substructures, potentially explaining the relatively low within-parcel homogeneity.

**Fig 5.**
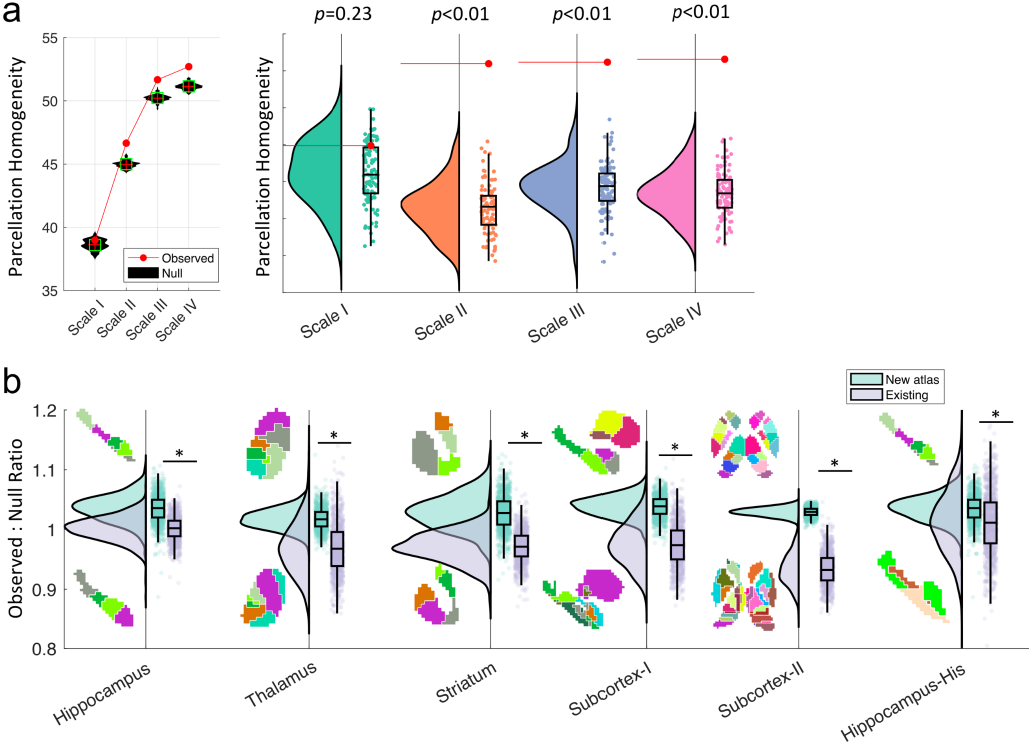
Parcellation homogeneity. **a**, Parcellation homogeneity of the new atlas was benchmarked to random parcellations of the subcortex comprising an identical number of regions and a comparable distribution of region sizes. Violin plots show the distribution of parcellation homogeneity measured within ensembles of 100 random parcellations. The p-value shown for each scale is the proportion of random parcellations comprising the ensemble that were more homogeneous than the observed parcellation. Horizontal red lines indicate the observed parcellation homogeneity values. Bottom and top edges of the boxes indicate 25th and 75th percentiles of the distribution, respectively. The vertical axis is omitted to aid visualization. The null hypothesis of parcellation homogeneity that is no better than chance could be rejected for all scales, except Scale I. **b**, Parcellation homogeneity comparison between the new atlas and existing parcellation atlases of the entire subcortex and specific subcortical nuclei (Supplementary Table S3). Parcellation homogeneity was computed separately for each individual (REST2, n=1021) and normalized by the random parcellation homogeneity, yielding and observed-to-null ratio. The box plots show the distribution of this ratio across individuals for the new (turquoise) and existing (violet) parcellation atlases. Homogenous parcellations have a ratio that exceeds one. Asterisks indicate that the observed-to-null ratio is significantly higher for the new atlas, relative to the existing atlas used for comparison (*p*<0.001, two-sample t-test). Anatomical visualizations of the new and comparison atlases are shown above and below the box plots, respectively.

Parcellation homogeneity was compared between the new atlas and existing imaging- and histology-based subcortical parcellations (Supplementary Table S3). Striatal, hippocampal and thalamic parcellations were isolated from the new atlas to enable comparison with existing parcellations of these regions. The new atlas was significiantly more homogeneous than existing imaging-based parcellations of the whole subcortex (Subcortex-I: *p*<0.001; Subcortex-II: *p*<0.001), parcellations of hippocampus (*p*<0.001), thalamus (*p*<0.001) and striatum (*p*<0.001) as well as a histological atlas of the hippocampus (Hippocampus-His: *p*<0.001; Fig 5b, Supplementary Fig S12). These results were replicated in an independent dataset comprising ten healthy adults (6 males, mean age 26±2.1yrs, Supplementary Fig S13).

In auxiliary analyses, we tested whether task-evoked activity mapped onto the new atlas and respected atlas boundaries. To this end, the standard deviation in task-evoked activity was computed across voxels comprising each region and then averaged over all regions. This was repeated for Scale I-IV. The lower the standard deviation, the more circumscribed task-evoked activity was to particular atlas regions. For all task conditions, we found that standard deviations were significantly lower for the new atlas than comparable random parcellations of the subcortex (*p*<0.01, Supplementary Fig S14). This cross-modality validation suggests that task-evoked activity is circumscribed to specific subcortical atlas regions more than expected by chance.

### Parcellation replication

Ultra-high field strength MRI can alleviate several technical challenges associated with subcortical imaging^26^. Hence, we sought to investigate whether the new atlas could be replicated using 7T fMRI. Qualitatively, gradient magnitude peaks were sharper and more circumscribed at 7T, compared to 3T (Fig 6). Gradientography, model selection and boundary delineation applied to 7T yielded a comparable, but more detailed, atlas of the subcortex, which also comprised four scales (Scale I: 16 regions, II: 34 regions, III: 54 regions, IV: 62 regions). The number of regions delineated at Scale I was identical between 3T and 7T (Fig 6). However, additional boundaries could be identified with 7T at Scale II-IV (Supplementary Fig S15 & S16). In particular, the amygdala was partitioned into medial and lateral subregions using 3T, whereas its medial extent was further divided into centromedial and superomedial components at 7T, and the body of hippocampus was further divided mediolaterally. Based on normalized mutual information (NMI), the spatial correspondence between the 3T and 7T atlases was excellent (Scale I: NMI=0.93, II: 0.88, III: 0.84, IV: 0.83; Supplementary Fig S16). Correspondence between 3T and 7T atlases was significantly higher than expected due to chance at all parcellation scales (*p*<0.01, Supplementary Fig S16).

**Fig 6.**
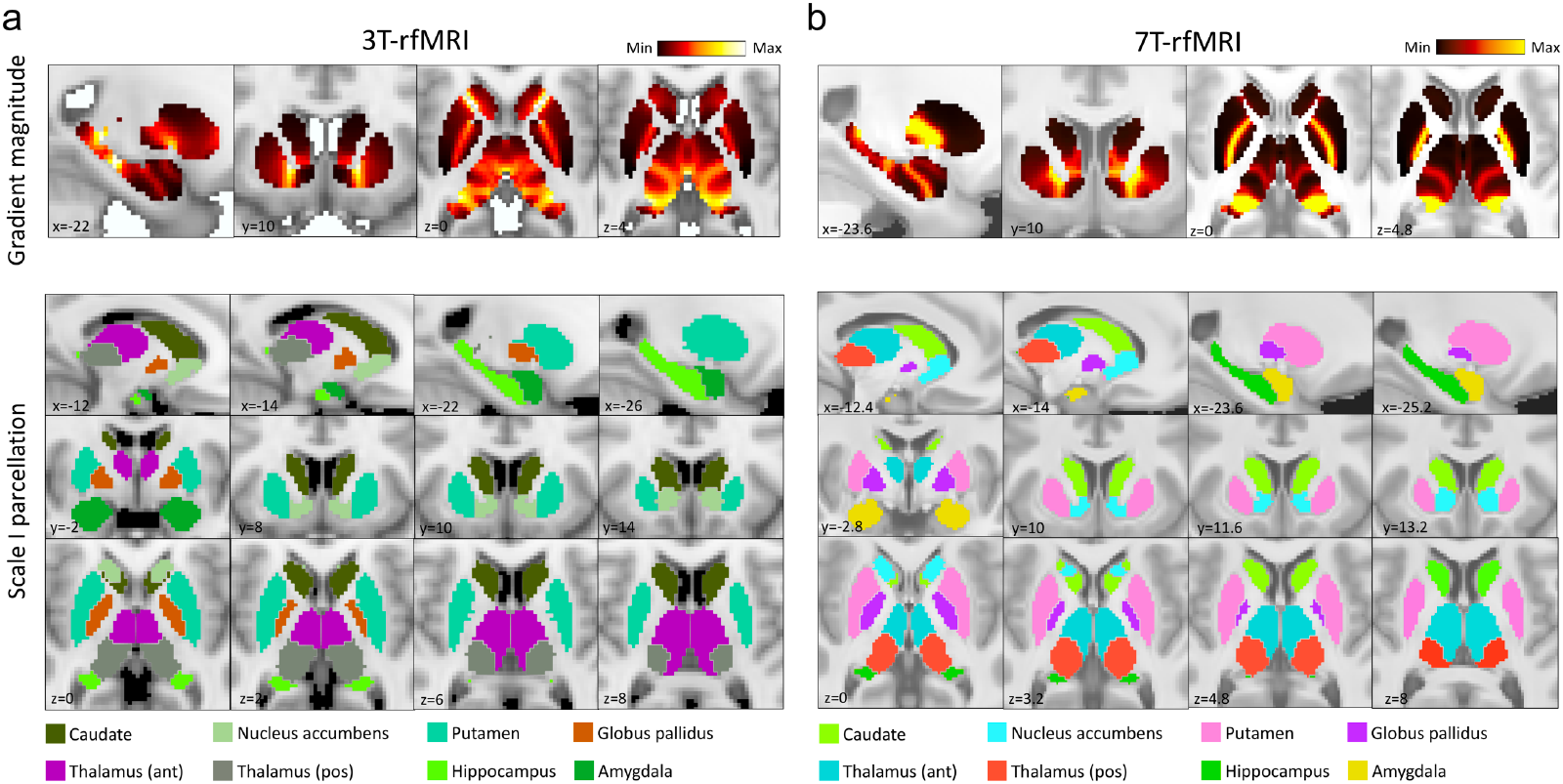
Replication of 3 Tesla parcellation using 7 Tesla resting-state functional MRI. **a**, Group-consensus gradient magnitude image (upper row) and Scale I atlas (lower three rows) delineated using 3T resting-state functional MRI (3T-rfMRI, n=1080). Bands colored in yellow-orange indicate local maxima in the gradient magnitude image. Homologous regions are colored identically. **b**, Same as **a**, but delineated using 7T resting-state functional MRI (7T-rfMRI, n=183). The 8 regions comprising Scale I were replicated with 7T imaging. All slice coordinates are indicated in Montreal Neurological Institute (MNI) space (mm). A reference structural MRI image is used as the background. ant, anterior; pos, posterior.

### Personalized parcellation

Having delineated a group-consensus atlas of the subcortex, we next sought to personalize the new atlas to account for individual variation in connectivity architecture^27, 28^. Following previous work on cortical atlas individualization^2^, a machine learning classifier was trained (n=100, REST2) to recognize putative connectivity fingerprints characteristic of each Scale IV region. The trained classifier was then used to delineate personalized boundaries for each region in an independent group (n=921, REST2). In particular, for each individual, voxels were assigned a classification probability of belonging to each region and a winner-takes-all rule was used to delineate personalized parcellations (Supplementary Fig S17).

Quantified with the Dice coefficient, the extent to which the personalized atlases recapitulated the group-consensus atlas varied markedly between individuals and regions. This was most evident within striatum (Fig 7a) and medial subdivisions of hippocampal head. Group-averaged Dice coefficients (Fig 7b) varied from 0.22 (subdivision of medial hippocampal head, HIP-head-m2-lh) to 0.85 (dorsoposterior putamen, PUT-DP-rh), for a classification probability threshold of 50%. Hotspots in the group-averaged classification probability maps were encircled by the group-consensus boundaries (Fig 7c), indicating self-consistency in classification performance. Individual variation in Dice coefficients was highly skewed, as evidenced for the dorsoposterior putamen and hippocampal body (Fig 7d). Specifically, a majority of individuals recapitulated the group-consensus (Dice>0.8), but 2-7% showed marked divergence (Dice<0.5). Boundary contraction was the most common source of divergence between individuals and the group-consensus atlas (Fig 7c). The same individuals consistently deviated from the group-consensus atlas for all regions, suggesting that this deviation was specific to individuals rather than regions (Supplementary Fig S18).

**Fig 7.**
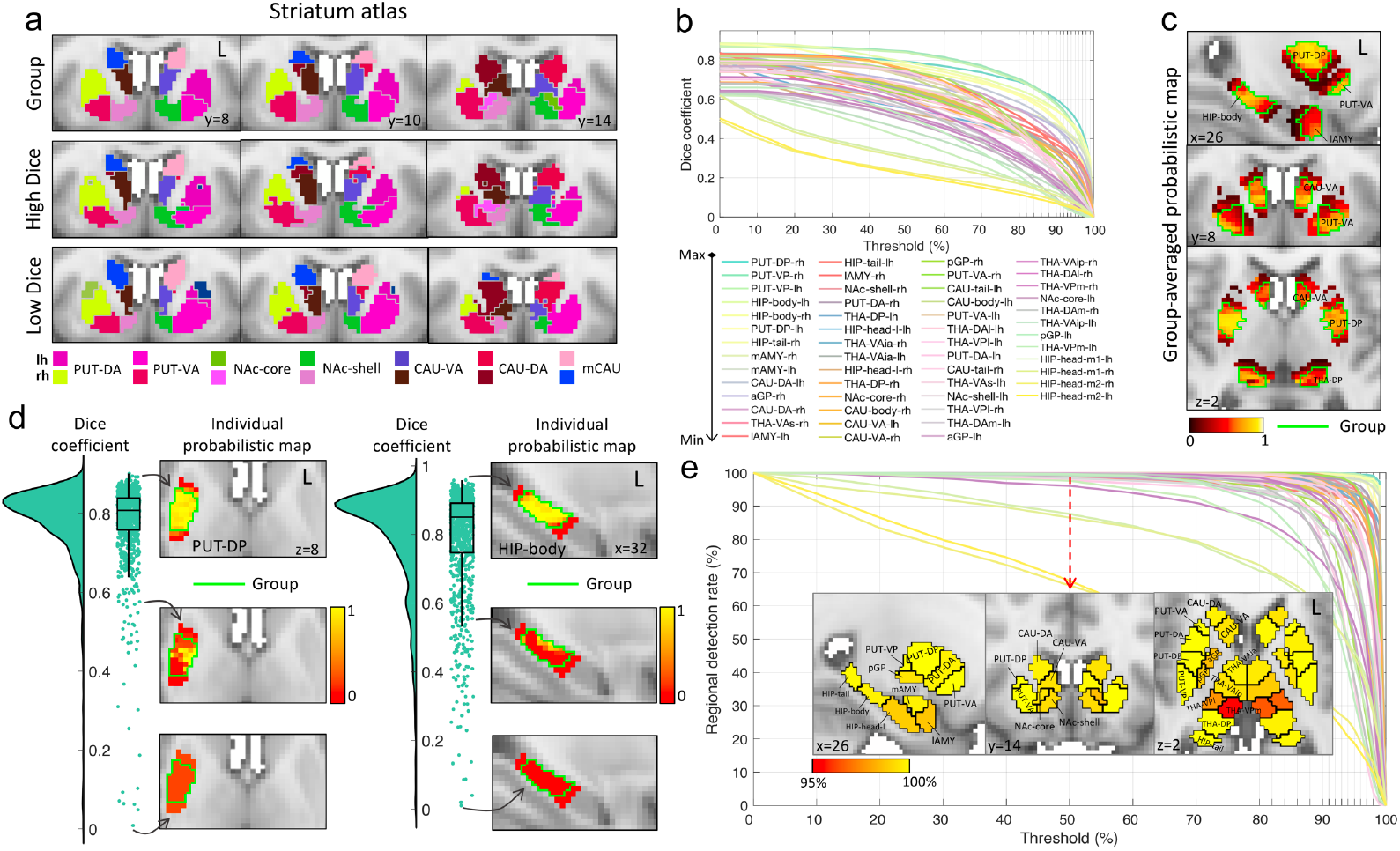
Personalization of the Scale IV atlas. **a**, Group-consensus atlas for striatal regions (top row) and personalized atlases for representative individuals with relatively high (middle row, Dice=0.81) and low (bottom row, Dice=0.15) similarity with the group-consensus atlas. **b**, Curves show group-averaged Dice coefficients as a function of classification probability threshold (0-100%). A separate curve is shown for each of the 54 regions comprising the Scale IV atlas. Classification probability is the probability that a subcortical voxel is classified as belonging to a specific region in a given individual. Parcellations were realized for each classification probability threshold and compared to the group-consensus atlas using the Dice coefficient. The area under the curve was computed for each region and the legend is shown in descending order from top-to-bottom and left-to-right. **c**, Group-averaged classification probability maps exemplified in 7 regions. Voxel color is scaled by the classification probability of the voxel belonging to the given region. The group-consensus boundary is colored in green. **d**, Box plots show the distribution of individual variation in Dice coefficients of dorsoposterior putamen (PUT-DP, left) and hippocampal body (HIP-body, right). Each data point represents the Dice coefficient of one of the 921 individuals used to evaluate classification performance. Classification probability maps are shown in anatomical space for representative individuals with relatively high (top), moderate (middle) and low (bottom) Dice coefficients. The group-consensus boundary is shown in green. **e**, Curves show regional detection rates as a function of the classification probability threshold. The inset shows anatomical renderings of regional detection rates for a nominal classification probability threshold of 50%. The two hippocampal regions (bilateral HIP-head-m1 and HIP-head-m2) with the lowest regional detection rates are obscured in the slices shown. All slice coordinates are indicated relative to Montreal Neurological Institute (MNI) space (mm). A reference structural MRI image is used as the background in anatomical visualizations. PUT, putamen; NAc, nucleus accumbens; CAU, caudate nucleus; GP, globus pallidus; HIP, hippocampus; AMY, amygdala; THA, thalamus; a, anterior; p, posterior; v, ventral; d, dorsal; m, medial; l, lateral; s, superior; i, inferior.

It was possible for the classifier to fail to detect particular regions in some individuals. For each region, the proportion of individuals for which at least one voxel was assigned to that region, called the regional detection rate, was analyzed as a function of classification probability threshold (Fig 7e). The regional detection rate for a classification probability threshold of 50% was rendered on the group-consensus atlas (Fig 7e, inset). The averaged regional detection rate was 97.8%, with detection rates falling below 95% for only two bilateral regions (i.e. HIP-head-m1 and HIP-head-m2). This suggests that the group-consensus atlas is representative of most individuals, although a minority of individuals may benefit from delineation of individual-specific atlases to account for inter-individual variation in regional boundaries.

### Rest compared to task-evoked conditions

We have thus far focussed on resting-state fMRI. Using task-evoked fMRI, we next investigated the extent to which the functional topography of the subcortex reconfigures in response to changing cognitive demands^29^. For each of seven tasks^30^ and the two rest sessions, gradient magnitudes (Gradient I) were projected onto the previously mapped ventral and dorsal streamlines, yielding task-evoked diversity curves.

Qualitatively, diversity curve peaks were consistent between the seven task conditions, but some peak locations and magnitudes differed relative to the rest condition (Supplementary Fig S19a). The most prominent task-rest variation was bifurcation of the boundary separating putamen-NAc and NAc-caudate during the task conditions. Functional separation between amygdala and the anterior extreme of hippocampus, posterior hippocampus and thalamus, as well as anterior-posterior partition within the thalamus were also weaker during task, compared to rest, whereas the separation between anterior and posterior hippocampus was stronger and extended anteriorly (Supplementary Fig S19a). To investigate this further, the Pearson correlation coefficient was used to quantify similarity in diversity curves between all pairs of conditions, yielding correlation matrices (Supplementary Fig S19b) that were visualized as correlation graphs (Supplementary Fig S19c). Diversity curves were highly consistent between the two rest sessions (dorsal: *r*=0.91, *p*<0.0001; ventral: *r*=0.98, *p*<0.0001), but were distinct from the seven task conditions, resulting in distinct task and rest modules (Supplementary Fig S19d). Task-evoked reconfiguration of functional organization was not due to differences in the number of fMRI volumes acquired between task and rest conditions (see Methods).

### Links between subcortical topography and cortical networks

Next, we investigated functional connectivity between canonical cortical networks and subcortical regions (Scale IV). Similarity in patterns of functional connectivity with the subcortex was quantified between pairs of 12 established cortical networks^3^ (Supplementary Fig S20a). Based on similarity in the subcortical areas to which they most strongly connected, the 12 cortical networks could be parsed into three distinct groups and positioned on a task-positive to task-negative^18^ organizational axis (panel b). One extreme of the axis comprised cortical networks that typically show increased activity during attention-demanding cognitive tasks (i.e. task-positive), including the salience, cingulo-parietal, cingulo-opercular, frontoparietal and dorsal attention network. In contrast, the default mode network, which is often characterized by task-related deactivation (i.e. task-negative) occupied the other extreme. Between them, sensorimotor related networks (visual, auditory, dorsal and ventral somatomotor) as well as ventral attention and retrosplenial-temporal networks formed a third group, in which the latter two also linked to the default mode network, suggesting a transition from sensorimotor toward task-negative functions.

### Using the new atlas to relate human behavior to subcortical connectivity

Finally, we used the new atlas to unveil new relationships between subcortical functional connectivity and individual variation in human behaviors^31^. Five behavioral dimensions characterizing: i) cognition, ii) illicit substance use, iii) tobacco use, iv) personality and emotional traits, and v) mental health were considered (Fig 8a, b & Supplementary Fig S21). These dimensions were derived from 109 behavioral items (Supplementary Table S4) using independent components analysis (see Methods). Using the network-based statistic^32^, higher tobacco use was found to be significantly associated with lower functional connectivity in thalamo-caudate, thalamo-NAc, and hippocampo-NAc circuits (Fig 8c). This is consistent with the involvement of the cortico-basal ganglia-thalamo-cortical circuit and mesolimbic dopamine pathways in reward and goal-directed behaviors^33^, such as smoking (see Discussion). Although connectivity explained 4% of individual variation in the behavioral dimension characterizing tobacco use, the relationship was highly consistent across all four parcellation scales (Scale I: *r*=-0.16, *p*<10^−6^; II: *r*=-0.17, *p*<10^−7^; III: *r*=-0.19, *p*<10^−8^; IV: *r*=-0.19, *p*<10^−8^) and reproducible across fMRI sessions (Supplementary Fig S22). The remaining four dimensions did not significantly relate to subcortical connectivity after correction for multiple testing.

**Fig 8.**
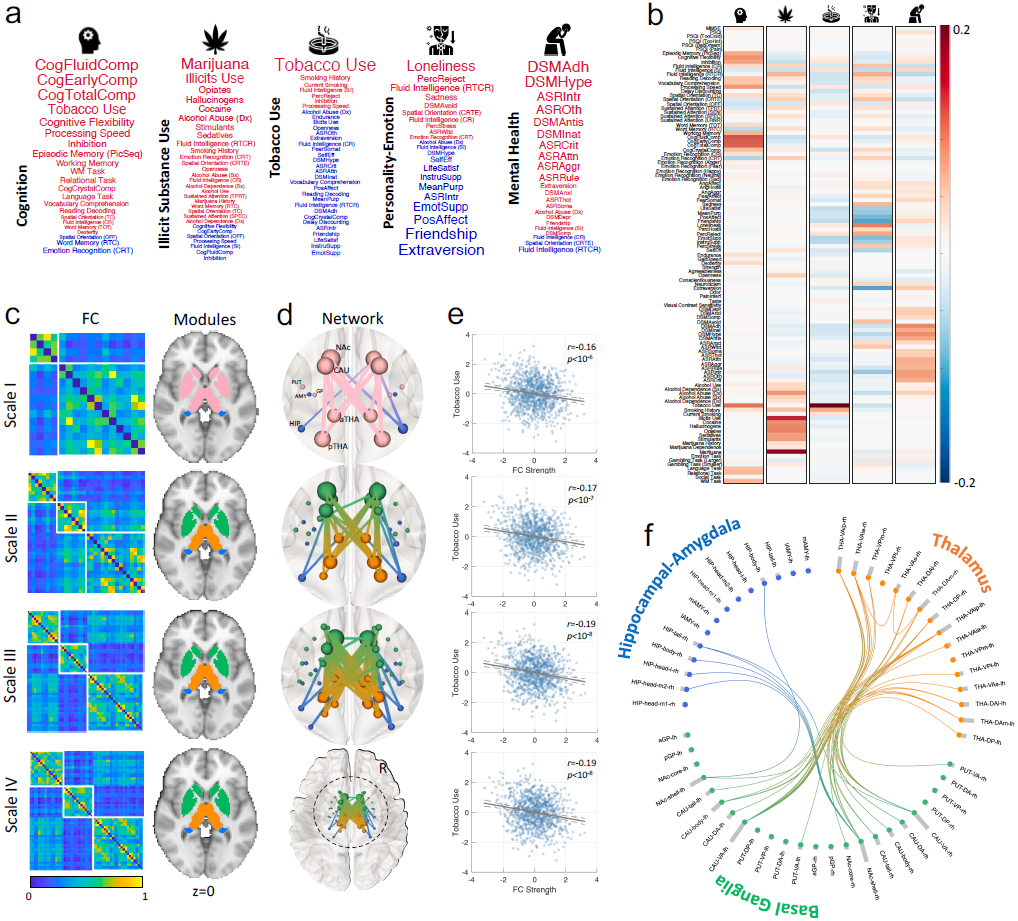
Associations between individual variation in behavioral dimensions and subcortical functional connectivity. **a**, Word clouds characterizing five behavioral dimensions derived from 109 behavioral items using independent component analysis (ICA). Only the most highly weighted behavioral items are shown. Font size is scaled according to the item’s absolute weight in the ICA de-mixing matrix. Font color indicates weight polarity (red-positive, blue-negative). **b**, Consensus de-mixing matrix estimated using ICA. Rows correspond to behavioral items. Columns correspond to the five latent dimensions. Cell color is modulated by de-mixing weight. **c**, Group-averaged functional connectivity matrices for Scale I-IV atlases. Rows and columns are reordered to accentuate modular structure (white blocks). Modules are also rendered in anatomical space, where regions comprising the same module are colored identically. Scale I comprises two modules (blue: hippocampus and amygdala, pink: basal ganglia and thalamus). The latter module was divided into two submodules that separated basal ganglia (green) and thalamus (orange) at Scales II-IV. **d**, Subcortical network for which functional connectivity significantly associates with individual variation in the behavioral dimension characterizing tobacco use. The linear association was significant for all atlas scales (Scale I-IV, *p*<0.001, network-based statistic). Nodes and edges are colored according to module allegiance. Thicker edges are associated with stronger effects. **e**, Scatter plots and lines of best fit for the linear association between subcortical functional connectivity and individual variation in the dimension characterizing tobacco use. Lines of best fit are indicated in black and dashed gray lines show 95% confidence intervals. **f**, Subcortical network associated with tobacco use shown as a circular network for the Scale IV atlas, highlighting the role of thalamo-striato-hippocampal circuits. Nodes and connections are colored according to module allegiance. The histogram (gray bar) shown for each node indicates nodal degree. Behavioral icons sourced from: https://www.flaticon.com/.

## Discussion

The field’s preponderance for the cerebral cortex, coupled with technical challenges inherent to subcortical imaging^34-36^, has led to the subcortex remaining a *terra incognita*^37^. Through the provision of high-quality, ultra-high field strength MRI data, collaborative initiatives such as the HCP^19^ herald new opportunities to overcome some of the challenges associated with reliably imaging this deep and anatomically complex brain structure.

To address the incomplete representation of subcortical regions in many leading MRI atlases, we used multi-session MRI of brain function acquired in more than 1000 healthy adults participating in the HCP to unveil fundamental new insight into the organizational architecture and connectivity of the human subcortex. Putative functional boundaries were recursively delineated at locations of sharp spatial change in subcortical-to-whole-brain functional connectivity fingerprints, which led to semi-automated delineation of a new multiscale parcellation atlas. The new atlas is openly available as a standalone parcellation and has also been integrated into established cortex-only^2, 3^ and cerebellar^15^ atlases. Crucially, the new subcortical atlas was replicated using ultra-high field strength MRI, and the regions delineated were functionally homogeneous and identifiable within individual participants, enabling delineation of personalized atlases that accounted for individual variation in connectivity. We used the new atlas to link cortical networks with the subcortex and uncover a robust and reproducible relationship between individual variation in behavioral dimensions and subcortical functional connectivity.

### Reconciling parcellations and gradients

Topographic representation of variation in brain structure, function and connectivity usually follows one of two paradigms: i) a mosaic of discrete parcels separated by hard boundaries^2-4, 15^; and more recently, ii) gradients of continuous topographic variation^12-14, 16, 25, 38^. We introduced a principled way forward to reconcile these two seemingly incompatible paradigms, resulting in a hybrid representation that parsimoniously accommodated both discrete boundaries and continuous gradients. Boundaries were only delineated at locations where gradient magnitudes were significantly larger than what would be expected due to chance. This was formalized with a model selection process and accompanying null model to test the null hypothesis of an absence of discrete spatial boundaries at candidate locations. Without model selection, boundaries would be delineated for all gradient magnitude peaks, potentially resulting in spurious boundaries that represent geometric constrictions rather than sharp changes in connectional topography. The bow-tie constriction shown in Supplementary Fig S5b provides an example of spurious boundary delineation, where a gradient peak appears at the bow knot, despite uniform spatial variation in connectivity. While the sheet-like geometry of the cerebral cortex is less likely to be affected by this phenomenon, complex subcortical geometry is more vulnerable to false positive boundaries and over-parcellation.

Gradientography was developed to enable gradient visualization and accurate localization of candidate boundaries. Gradientography is an fMRI analogue of diffusion MRI tractography and can be considered a form of fMRI tractography. Whereas tractography is conventionally used to map white matter fiber trajectories, gradientography, as introduced here, was used to map trajectories of maximum change in connectopic gradients. Future applications of gradientography are not limited to the subcortex and the method could provide a bridge between cortical gradients and parcellations.

### New multiscale subcortical atlas

The new subcortical atlas comprises four scales, labelled Scale I to IV, each providing a self-contained parcellation atlas. Additional scales were not warranted because the model selection procedure decided against boundary delineation for all regions comprising Scale IV. Researchers seeking to use the new atlas should select the atlas scale that provides adequate resolution to address the research question at hand and approximately matches the scale of any cortical atlas that is used in terms of the total number of regions. While the new atlas is unimodal, resting-state fMRI connectivity is argued to be one the most useful modalities in predicting areal boundaries^2^.

Hierarchy is a hallmark of brain organization^39^ and hierarchical organization is not necessarily a *fait accompli* of the recursive application of the model selection procedure used in this study (see extended discussion in Supplementary S2). Scale I of the atlas recapitulates subcortical regions delineated in existing MRI atlases, such as those available as part of Lead-DBS, Freesurfer and FSL. The position of gradient peaks enabled automated boundary demarcation between amygdala, hippocampus, thalamus, caudate, putamen, NAc and GP. Discrete partition of the basal ganglia into the latter four regions as well as the separation between amygdala and hippocampus were clearly supported by model selection. Contrastingly, the few existing connectivity-based subcortical atlases^8, 40^ generally do not preserve established anatomical boundaries^41^, potentially suggesting structure-function divergence or methodological variation. Given that these broad subcortical regions are distinguishable anatomically, rudimentary alignment with functional architecture might be expected^1^. Indeed, this assumption motivates parcellation of regions in isolation, such as the hippocampus^38, 42^, striatum^25, 43^ and thalamus^44^, without first establishing functional boundaries between them. Despite correspondence between the principal connectional topography (Gradient I) and anatomy at Scale I, fine variation within specific structures was captured by the secondary connectional gradients. As shown in Supplementary Fig S10c, the dominant striatal gradient separates putamen, NAc and caudate, whereas the second and third gradients are organized along a dorsal-to-ventral axis, reflecting connectional variation between dorsal (putamen and caudate) and ventral (NAc) striatum^45^.

Scale II delineates functional divisions within each of the Scale I regions. NAc was partitioned into its shell and core, anterior (head) and posterior (body and tail) caudate subdivisions were identified, and amygdala was divided into lateral and medial subregions. A striking divergence between anatomy and function was evident for the GP. GP is anatomically divided along a mediolateral axis into internal and external GP^41^, whereas functional connectivity gradients indicated an anteroposterior partition. Scale III and IV differ only in parcellation of thalamus and hippocampus.

### Parcellation validation

While histological validation is the gold standard, functional parcellations do not necessarily converge to anatomical or cytoarchitectural divisions to be deemed useful^1^. To establish confidence in the new atlas, we first demonstrated that all atlas scales, except Scale I, were significantly more homogenous^3^ than expected due to chance. While Scale I regions converged with subcortical anatomy, they encompass multiple functional systems, rendering Scale I too coarse to achieve homogeneity. Homogeneity of functional signals within delineated subregions is a fundamental principle of functional parcellation^1^. Interestingly, existing structural connectivity and histological parcellations of thalamus^44^ and hippocampus^24^, respectively, were not functionally homogenous (Supplementary Fig S12), indicating a need for modality-specific atlases.

Second, we reproduced the majority of regions using 7T fMRI. The improved contrast-to-noise ratio and spatial resolution of 7T compared to 3T MRI is particularly important for subcortical imaging^26^. Despite remarkable consistency between 3T and 7T atlases, 7T enabled delineation of finer structures. For example, a bipartite partition of amygdala (medial and lateral) was delineated using 3T, whereas 7T aided further division of the medial amygdala into central and superior components, which broadly recapitulate the three cytoarchitectural zones of the amygdala^24^. On the other hand, NAc was consistently parcellated into two subdivisions using 3T and 7T, presumably representing the immunohistochemically distinct shell and core subdivision that is common to primates^46^. Therefore, while subcortical cartography is viable with functional connectivity inferred from high-quality 3T fMRI, the level of detail that can be resolved with 7T is superior for several regions.

The subcortex atlas was personalized to account for individual connectivity differences. We showed that while the group-consensus atlas was sufficient to represent the majority of individuals, personalized subcortex atlases were warranted for a small proportion of others. Further work is needed to understand whether individual variation in subcortical connectional topography relates to anatomy, task-evoked activation and behavior. Individual-specific atlases of the subcortex can potentially facilitate personalization of targeted therapies in psychiatry^47^ and improve prediction of behavior^28^.

### Dynamic and intrinsic subcortical topography

Using task fMRI, we identified subtle reconfigurations in the functional topography of the subcortex in response to changing cognitive demands. While a previous study^29^ examined variation in parcel size in response to task stimuli, diversity curves enabled the characterization of changes in functional boundaries. During task conditions, the anterior putamen integrated more extensively with higher-order association networks, whereas the hippocampus segregated along its long-axis, yielding a more integrated anterior hippocampal-amygdala system that may have served cognitive-emotional processing^22, 23^, and a more integrated posterior hippocampal-thalamus that potentially benefited learning and spatial memory^22^. These task-induced dynamic reconfigurations may reflect a process of neuronal recruitment into specific functional processing systems, to cope with cognitive demands. The flexible reconfiguration of neural networks may largely depend on the intrinsic functional organization within local cortical circuits, given that a task-positive to task-negative organizational axis was evident in the subcortex (Supplementary Fig S20). However, context dependent recruitment across intrinsic parcels, leading to the adjustment of functional boundaries, may additionally assist adaptive task responses. Further work is needed to understand task-evoked reconfigurations of functional boundaries, not only in the subcortex, but across the brain.

### Tobacco and subcortical connectivity

We performed a hypothesis-free decomposition of HCP cognitive and behavioral measures into five latent dimensions to facilitate investigation of relationships between brain connectivity and behavior. Previous HCP investigations typically investigate hundreds of individual measures, leading to a formidable multiple testing problem, or preselect a limited number of specific behavioral domains or measures. The five latent dimensions identified here condense information from more than a hundred individual measures.

The new atlas unveiled a novel relationship between individual variation in functional connectivity and the behavioral dimension characterizing tobacco use. Higher tobacco use was associated with lower thalamo-caudate, thalamo-NAc, and hippocampo-NAc functional connectivity. The shell of NAc and adjacent ventroanterior caudate (CAU-VA) were most implicated (Fig 8c). This accords with the primary role of the shell of NAc and anterior caudate in reward anticipation, incentive motivation and action^48, 49^. NAc integrates glutamatergic (excitatory) inputs from hippocampus and thalamus^50, 51^, enabling the regulation of dopamine neuron activity in ventral tegmental area^51^. While nicotine, the main neuroactive compound of tobacco stimulates dopamine release and transmission in NAc, especially the medial shell^52^, chronic nicotine administration yields neuronal damage in hippocampus and striatum^53, 54^, potentially manifesting in lower functional connectivity in the thalamo-striato-hippocampus system. Therefore, nicotine’s putative down-regulation of the mesolimbic circuit may provide insight into the increased prevalence of tobacco smoking in individuals with schizophrenia^55^. In particular, this down-regulation effect of nicotine on the hyper-dopaminergic state in the mesolimbic system^56^ may correspond with the up-regulation effect of nicotine in the prefrontal cortex^57^, alleviating symptoms in the disorder.

## Conclusions

We unveiled the complex topographic organization of the human subcortex and parsed this complexity into four scales constituting a new atlas for the brain’s remaining *terra incognita*. The new atlas is scalable, personalizable, reproducible at ultra-high field strengths and openly available to anyone. Future work will focus on addition of structures such as subthalamic regions, red nucleus and hypothalamus. The new atlas can be incorporated with existing cortex-only atlases to aid the investigation of cortico-subcortical interactions and mapping of holistic connectomes using MRI^58^.

## Methods

### Data and pre-processing

Human neuroimaging data acquired as part of the Human Connectome Project S1200 release^19^ was analyzed here. Participants were healthy young adults ranging between 22 and 37 years old. Some participants were genetically related. These relationships were controlled in statistical analyses by constraining permutations^59, 60^. HCP datasets used in this study include: i) Two sessions (REST1 and REST2) of resting-state functional magnetic resonance imaging (rfMRI) acquired using multiband echo planar imaging (EPI) on a customized Siemens 3T MR scanner (Skyra system), where each session comprised two runs (left-to-right and right-to-left phase encoding) of 14min and 33s each (TR=720ms, TE=33.1ms, voxel dimension: 2mm isotropic). The two runs were temporally concatenated for each session, yielding 29min and 6s of data in each session. Concatenation of the two different phase encoded data ensured that any potential effect of phase encoding on gradient direction was counter balanced by the opposing phase encoding; ii) Task-evoked fMRI (tfMRI) acquired using the identical multiband EPI sequence as the resting-state fMRI, however, the run duration ranged between 2-5mins, depending on the specific task (7 tasks); iii) Resting-state fMRI acquired on a Siemens Magnetom 7T MR scanner using a multiband acquisition (TR=1000ms, TE=22.2ms, voxel dimension: 1.6mm isotropic), where participants completed four runs of approximately 16 minutes each. The first two runs (posterior-to-anterior and anterior-to-posterior phase encoding) were used in this study. We refer to this dataset as 7T-rfMRI. Details pertaining to the acquisition of rfMRI^26, 61^ and tfMRI^30^ are described elsewhere. The seven tfMRI tasks were chosen to tap a broad range of cognitive and affective processes and activated a wide range of neural systems^30^. Participants that had completed at least two runs in each dataset were included. Supplementary Table S1 shows the sample size and basic participant demographics for each dataset.

The second 3T-rfMRI session (REST2) was used for replication. Minimally preprocessed volumetric rfMRI and tfMRI data were sourced from the online HCP repository. Details of the minimal preprocessing pipeline can be found elsewhere^62^. The pipeline includes correction of head motion and spatially structured physiological noise with ICA+FIX^63, 64^. ICA-based denoising methods outperform, or equal, the performance of alternative approaches on most benchmarks^65^. After eliminating spatial artefacts and distortions, the functional images were spatially aligned to the MNI (Montreal Neurological Institute) standard space using the FNIRT nonlinear registration algorithm. MNI standard space herein always refers to the MNI ICBM 152 nonlinear 6th generation stereotaxic registration model. Additional nuisance regression and temporal filtering were not performed as part of the pipeline. However, head motion registration parameters estimated as part of spatial alignment were curated, reduced to a single summary measure known as framewise displacement^66^ and included as a confound in statistical analyses to control for individual variation in head motion^67, 68^.

The subcortical signal-to-noise ratio (SNR) and sensitivity of the blood-oxygenation level dependent (BOLD) signal was lower compared to the cortex because T2* signal decay times are shorter in subcortical regions compared to cortical regions, due to iron enrichment of subcortical tissue^34^. Therefore, to enhance subcortical SNR and BOLD sensitivity, the minimally preprocessed data were subjected to two additional operations (Supplementary Fig S8a). First, spatial smoothing was performed with a Gaussian smoothing kernel of 6mm FWHM (full width at half maximum) for 3T and 4mm FWHM for 7T images. This choice of kernel width was guided by evaluation of gradient magnitude images derived from a range of smoothing kernels (4, 6, 8 mm) and filtering options (i.e. Wishart filter, see below). Second, to further improve subcortical SNR and suppress the effects of unstructured and autocorrelated noise that share similar temporal and spatial properties, the fMRI time series were reconstructed from a principal component analysis (PCA), where the components were filtered according to an empirically-fitted Wishart distribution to dampen the effects of noise^2, 69, 70^. This PCA-based data reconstruction (i.e. Wishart filter) has been previously used in a multi-modal cortical parcellation study^2^ and comprehensively documented by Glasser and colleagues^69, 70^. The Wishart filter was performed over all gray matter voxels for each individual separately in all datasets. In addition, task block regressors were created for each task condition based on the time of onset, the duration and the relative magnitude of each stimulus, and then convolved with hemodynamic response functions (*spm_hrf*.*m* function from SPM12, https://www.fil.ion.ucl.au.uk/spm). The relevant regressors corresponding to each task were then regressed from the tfMRI data, to ensure that functional connectivity was disambiguated from the potential confound of task co-activation effects^71^. The residuals remaining from this regression were used for all subsequent analyses.

### Mask delineation

A binary subcortex mask comprising the left and right basal ganglia, thalamus, hippocampus and amygdala was delineated using three steps. First, the probabilistic Harvard-Oxford Subcortical Structural Atlas (https://fsl.fmrib.ox.ac.uk/fsl/fslwiki/Atlases) was used to delineate a binary mask for each of the following subcortical regions: thalamus, caudate, putamen, nucleus accumbens (NAc), globus pallidus (GP), hippocampus and amygdala. Voxels with a probability of 50% or more of belonging to one of these regions were included in the relevant mask, except for GP. A higher threshold (60%) was applied to GP to minimize potential white matter contamination of the fMRI signal, due to the numerous myelinated axons of the striato-pallido-nigral bundle abutting the periphery of this region^72^. Second, the masks for each individual subcortical region were merged to yield a binary mask for the entire subcortex. Finally, the subcortex mask was left-right symmetrized. Left-right symmetrization accords with previous subcortical atlas literature^41, 73^ and enabled symmetrization of subsequent computations to improve SNR. Symmetrization is not an unprecedented step, given that intra-hemispheric asymmetries in the volume of subcortical nuclei are less evident in healthy young adults, compared to cortical asymmetries^74^. Mask symmetrization was achieved by reflecting the mask’s left hemisphere around the mid-sagittal plane and computing the intersection between the mask’s right hemisphere and this reflection. Voxels comprising the intersection yielded a left subcortex mask, which was merged with its reflection to yield the final left-right symmetric subcortex mask. The final mask comprised a total of *N*=7,984 voxels (63.87cm^2^). The subthalamic nuclei, substantia nigra and comparably small subcortical nuclei were not included because the spatial resolution of the fMRI data was deemed insufficient to accurately resolve putative substructures within these nuclei (e.g. pars reticulata and compacta). Sub-millimetre fMRI and multi-echo planar imaging would benefit investigation of the functional architecture of these exceedingly small nuclei^75^. A binary gray matter mask was delineated using the MNI152 probabilistic gray matter atlas. Voxels belonging to cortical gray matter were included in the gray matter mask as well as any voxels comprising the final subcortex mask. fMRI data was consistently absent for a majority of individuals in a small proportion of voxels comprising the gray matter mask. After eliminating these voxels, the final gray matter mask comprised *M*=164,360 voxels (1314.9cm^3^), which included the midbrain, pons and cerebellum. The subcortex and gray matter masks were delineated in 2mm isotropic space and were used for the 3T fMRI data. To suit the 7T-rfMRI data, which was in 1.6mm isotropic space, the masks were resliced to this resolution and median filtered to smooth the mask edges.

### Functional connectivity and eigenmaps

Whole-brain functional connectivity was mapped for each subcortex voxel. Spatial gradients in the resulting maps were then computed to yield a continuous representation of functional connectivity variation across the subcortex. Connectopic mapping^12^ was used to map spatial gradients for each individual, which involved computing a sequence of eigendecompositions to yield a Laplacian eigenmap^76^. Specifically, following the procedure described by Haak and colleagues^12^, the temporally concatenated fMRI signals were represented for each individual in a matrix of dimension *T* × *M*, where *T* denotes the number of time frames and *M* denotes the number of gray matter voxels. Principal component analysis (PCA) was used to reduce the dimensionality of this matrix to *T* × (*T* − 1). The fMRI signal at each subcortical voxel was then correlated (Pearson correlation) with each column of the PCA-transformed matrix, resulting in a connectivity matrix of dimension *N* × (*T* − 1), where *N* denotes the number of subcortex voxels. Correlation coefficients were *r*-to-*z* transformed using the Fisher transformation. Each row of this matrix provided a connectional fingerprint^20^ for a particular subcortical voxel in the PCA-transformed space. Similarity in the connectional fingerprints between each pair of subcortical voxels was then quantified with the η^*2*^coefficient, resulting in a symmetric matrix of dimension *N* × *N* for every individual. The similarity matrix was transformed into a sparse graph using the weighted adjacency matrix, where the weights of all connections with a Euclidean distance less than ϵ were set to zero. The connection density was determined by the smallest value of ϵ that ensured a connected graph^12, 77, 78^. This threshold varied between individuals, yielding graphs with a connection density of 0.4% on average. The Laplacian matrix, *L*, was then computed according to *L* = *D* − *W*, where *D* denotes the diagonal matrix of node strengths and *W* denotes the sparse adjacency matrix. In particular, *D*_*ij*_ = ∑_*i*_ *W*_*ij*_ if *i* = *j*, otherwise *D*_*ij*_ = *0*. Finally, the eigenvectors and eigenvalues of the Laplacian matrix were computed. The smallest eigenvalue was necessarily zero, while all other eigenvalues were positive due to the connectedness of the graph. The eigenvector with zero eigenvalue was a constant and discarded. The eigenvectors with the second, third and fourth smallest eigenvalue were referred to as Gradient I, II and III respectively. These steps are shown in Fig 1a. Each of the three gradients, or *eigenmaps*, characterized a continuous mode of spatial variation in functional connectivity across the spatial extent of the subcortex. We use the terms eigenmap and gradient interchangeably. Eigenvectors corresponding to the next largest eigenvalues typically explained substantially less variance in the Laplacian matrix than the first four eigenvectors (Supplementary Fig S2) and were thus given no further consideration here. The *N* values defining each gradient were projected onto the three-dimensional anatomy of the subcortex in MNI standard space for visualization and further analyses. Unless otherwise specified, the same process described above was used to estimate gradients for each individual and for each imaging modality, including 3T-rfMRI, 7T-rfMRI and tfMRI.

### Gradientography

Tractography is an established three-dimensional modelling technique to visually represent axonal fiber bundles using diffusion MRI^79^. We developed an analogous technique for fMRI called *gradientography* to characterize spatial gradients in functional connectivity. Gradientography was performed at the group level using a group-consensus representation of each eigenmap. In particular, similarity matrices were averaged across all individuals and eigenmaps were computed for the group-averaged similarity matrix using the same process described above. The resulting group-consensus eigenmaps were projected onto the three-dimensional anatomy of the subcortex in MNI standard space (Fig 1b, column I & Supplementary Fig S1, column I). Using the Sobel gradient operator, directional gradients were computed for each subcortical voxel comprising the three-dimensional image representing each eigenmap. Each subcortical voxel was thus endowed with a gradient directed along each of the three canonical axes as well as a gradient magnitude (Fig 1b, column II & Supplementary Fig S1, column II). We denote the gradient direction and magnitude at a given voxel with *g* = [*g*_*x*_, *g*_*y*_, *g*_*z*_]^*T*^ and |*g*|, respectively. The gradient magnitude indicated how rapidly the functional connectivity of a local area of subcortex changed per unit length. Before application of the Sobel operator, the peripheral boundary of the subcortex in each eigenmap image was dilated by approximately one voxel to diminish spurious gradients from appearing along the direction that is perpendicular to the boundary. Erosion was performed after application of the Sobel operator to reverse this dilation and restore the original volume of the subcortex. The gradient directions and magnitudes were approximately mirror symmetric in the sagittal plane (Supplementary Fig S3a). Therefore, to enhance SNR, the gradient directions were left-right symmetrized, consistent with the symmetrization of the subcortex mask. In particular, gradient directions in the left hemisphere were reflected into the right hemisphere around the mid-sagittal plane and averaged with the corresponding directions in the right hemisphere. These averaged gradient directions were then reflected back into the left hemisphere and gradient magnitudes were recomputed (Supplementary Fig S3b). This was performed separately for each of the three eigenmaps, and the difference in the gradient direction before and after symmetrization was quantified (Gradient I: θ=5.42 ± 4.98°; Gradient II: θ=5.04 ± 5.0°; Gradient III: θ=4.95 ± 5.36°).

We developed the new gradientography procedure to provide a principled framework to visualize and analyze spatial gradients in functional connectivity. First, three-dimensional tensors were fitted to each subcortical voxel using the constrained tensor model^80, 81^, which is synonymous with diffusion tensor imaging. The constrained tensor model is given by,

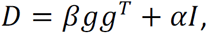

where *D* denotes a 3×3 symmetric tensor, *g* is a column vector denoting the normalized gradient direction computed with the Sobel operator,*I* is the identity matrix, and β and α are parameters that quantify the strength of anisotropic and isotropic diffusion, respectively. Here, we set β = *kc*|*g*| and α = *k*(1 − *c*), where *c* = 0.995 is a weighting factor that determines the extent of tensor anisotropy and *k* is a scale factor that governs tensor size for visualization purposes. In this way, cigar-shaped tensors indicated rapid change in subcortical functional connectivity along the orientation of the tensor’s longest axis. In contrast, spherical tensors indicated functional connectivity that was constant along all spatial directions. It is important to note that tractography was only dependent on the gradient direction, *g*. The choice of *β, α* and *c* only influenced visualization of the tensor field (Fig 1b, column III & Supplementary Fig S1, column III). While tensors were fitted separately for each of the three eigenmaps, an alternative methodology would have been to fit a bi- or multi-tensor model to each voxel^82^, enabling representation of multiple eigenmaps within a single tensor field. This is conceptually analogous to modelling crossing fibers in diffusion MRI tractography^83^. However, the potential utility of modeling interactions between crossing gradients in this way remains unclear, and thus each eigenmap was analyzed independently. We relegate multi-directional gradientography to future work. Streamline propagation was performed using the interpolated streamline method with a step length of 1 mm, angle threshold of 30° and seeding 20 streamlines from randomly chosen coordinates within each subcortical voxel, yielding approximately 15,000 streamlines in total. Streamline propagation was terminated when reaching the subcortex mask boundary. Gradientography was performed using Diffusion Toolkit software and visualized with TrackVis^84^. Streamlines were virtually unchanged when left-right symmetrization of the gradient directions was not performed. All streamlines were permitted to follow curvilinear trajectories of arbitrary complexity. The gradient magnitude image was projected onto the streamlines to parameterize the magnitude of change in functional connectivity and visualize transitions in the gradient field with respect to the curvature of streamlines. In particular, each spatial coordinate comprising a streamline was assigned a gradient magnitude based on trilinear interpolation of the gradient magnitude image (Fig 1b, right most). Gradientography was also performed for Gradients II and III using the same procedure (Supplementary Fig S1). While streamline trajectories varied between Gradients I, II and III, the locations of local maxima in the gradient magnitude images of the three gradients were quite consistent (Supplementary Fig S1 & S3). The gradient magnitude images for Gradient II and Gradient III were spatially correlated (Spearman correlation) to Gradient I (Gradient I/II: *r*=0.44, Gradient I/III: *r*=0.73), and thus only Gradient I was used to delineate functional boundaries.

We also observed that head motion (framewise displacement, FD) showed minimal effect on the group-consensus connectivity gradients. Specifically, individuals were ranked according to head motion and individuals in the top 20% (mean FD: 0.26±0.06 mm) and bottom 20% (mean FD: 0.10 ± 0.01 mm) defined two distinct subgroups. Group-consensus gradient magnitude images were computed for each of two subgroups and compared to the group-consensus gradient magnitude image generated within all individuals. We found that the group-consensus gradient magnitude images were strongly correlated between each pair of the three groups (Spearman correlation, *r*>0.9).

To preclude streamlines from traversing the slender anatomical gap between the tail of the caudate nucleus and the most anterior extent of the thalamus, the subcortex mask was separated at this precise location, giving rise to three spatially contiguous subcortical components: i) left basal ganglia, which comprised striatum and globus pallidus; ii) right basal ganglia; and, iii) left and right thalamus, hippocampus and amygdala. Because these three components were spatially discontiguous, tractography was performed separately for each component. A boundary between caudate and thalamus was thus imposed a priori, consistent with established anatomical knowledge. Due to left-right symmetrization of gradient directions, streamlines for the left and right basal ganglia were mirror symmetric, and thus it was sufficient to analyze only one of the hemispheres of the basal ganglia. We refer to the streamlines traversing the basal ganglia as the *dorsal streamlines* and the *dorsal connectional topography* (Fig 2a), while collectively referring to the streamlines traversing the left and right thalamus, hippocampus and amygdala as the *ventral streamlines* and *ventral connectional topography* (Fig 2b). Permitting streamlines to traverse the caudate-thalamus gap did not alter the location of local maxima in gradient magnitude, although they were less prominent in this case (Fig 1b).

### Diversity curves

The dorsal and ventral group of streamlines each comprised hundreds of distinct streamlines that characterized the connectional topographies associated with these regions. Streamlines shorter than 60mm and 160mm were discarded from the dorsal and ventral group respectively. The remaining streamlines within each group traversed trajectories that shared similar profiles and thus it was feasible to determine a single representative streamline for each group. To determine a representative streamline, the mean closest point distance^85^ was computed between all pairs of streamlines comprising a group. In particular, the distance from one streamline to another was determined by first computing the distance between each point on the first streamline to all points on the second streamline. The minimum distance was then determined for each point on the first streamline. The distance from the first to second streamline was defined as the mean of these minimum distances. This was repeated for all pairs of streamline to generate a *J* × *J* distance matrix, where *J* denotes the number of streamlines. The distance matrix was symmetrized in a way that the mean of the two distances between a pair of streamlines was used^85^. The representative streamline was chosen as the streamline with the shortest distance, on average, to all other streamlines. Having defined a representative streamline for the dorsal and ventral group of streamlines, it was then possible to compute a diversity curve with respect to the each of the representative streamlines. This yielded a curvilinear parameterization of the group-consensus eigenmaps and their gradient magnitudes, thereby enabling statistical inference. The diversity curve concept is described in detail elsewhere^16^.

Diversity curves were mapped in three steps: i) projection of the gradient magnitude image onto each streamline; ii) spatial alignment of streamlines to the representative streamline using dynamic time warping^86^; and iii) averaging across the aligned streamlines to yield a group-representative diversity curve to provide a basis for statistical inference. Specifically, each spatial coordinate comprising a streamline was assigned a gradient magnitude based on trilinear interpolation of the gradient magnitude image. While each streamline comprised the same number of equidistant coordinates, the exact lengths of streamlines varied, resulting in some degree of misalignment between streamlines in terms of coordinate-to-anatomy correspondence. For example, the tenth coordinate on a given streamline was not necessarily closest to the tenth coordinate on a neighbouring streamline. Thus, averaging across a set of streamlines could not be performed based on correspondence in the index of coordinates. Dynamic time warping was used to spatially align individual streamlines to the representative streamline and facilitate averaging of any measure parameterized along the length of each individual streamline.

Dynamic time warping is typically used to align signals in time, whereas the method was used here to achieve spatial alignment. A warping path was computed for each individual streamline, such that the sum of the Euclidean distance was minimal between the warped and representative streamline. This involved either stretching or compressing coordinates along the trajectory of each individual streamline to achieve alignment with the representative streamline. Alignment quality was assessed visually and streamlines with a warp distance exceeding the averaged warping distance over all streamlines (dorsal: 1.72mm, ventral: 0.54mm) were omitted. This yielded a total of 4,693 and 3,862 aligned streamlines for the dorsal and ventral group, respectively. The gradient (eigenmap) and gradient magnitude images projected onto each of these streamlines were then averaged for each point along the streamline trajectory, yielding a single gradient and gradient magnitude diversity curve for the dorsal (Fig 2c) and ventral group (Fig 2d). Gradient magnitude peaks in the diversity curves indicated putative functional boundaries between regions. Consistent with the gradient magnitude image across the entire subcortex, the gradient magnitude images for Gradient II and III were also spatially correlated (Spearman correlation) to Gradient I for both dorsal (Gradient I/II: *r*=0.79, Gradient I/III: *r*=0.79) and ventral (Gradient I/II: *r*=0.80, Gradient I/III: *r*=0.54) components. Statistical inference was therefore confined to the dominant gradient (Gradient I).

Focusing on the dominant gradient is not without precedent and this gradient shows good correspondence with underlying neuroanatomy^25^, although non-dominant gradients can reveal complementary connectivity topographies^12, 25^. Three local maxima were evident in dorsal streamline group, corresponding to established neuroanatomical boundaries between GP/putamen, putamen/NAc and NAc/caudate (Fig 2c). For the ventral streamline group, 9 local maxima were evident in the diversity curve representing gradient magnitude, suggesting putative functional boundaries between amygdala/hippocampus (ant), hippocampus (ant)/hippocampus (pos), hippocampus (pos)/thalamus(pos), thalamus (pos)/thalamus (ant), as well as left and right thalamus (Fig 2d).

### Null model

Although local maxima in the gradient magnitude images or the diversity curves representing these images indicated putative functional boundaries, these maxima could have also potentially arisen as a matter of random fluctuations reflecting inter-individual variation and finite sample effects, or due to the geometry of the subcortex (Supplementary Fig S5). Therefore, null data were generated to statistically determine whether local maxima in the gradient images were larger than expected due to these random (sample) effects or confounds. The null hypothesis was of a spatially homogeneous gradient magnitude, such that the rate at which whole-brain connectivity changed per unit length was constant at all points in the subcortex. Under this null hypothesis, any spatial heterogeneity was assumed to be due to sample effects or geometry. Functional boundaries were deemed to present at local maxima in the empirical gradient magnitude images when this null hypothesis could be locally rejected. Null data were generated by rewiring edges in the graph defined by the sparse adjacency matrix *W*. As detailed above (see Functional connectivity and eigenmaps), *W* defines a graph in which each node corresponds to a subcortical voxel and the edge weights quantify similarity in functional connectivity profiles. A rewiring algorithm was developed to randomly reposition each edge, subject to the constraint that edges were more likely to be placed between pairs of voxels in close spatial proximity. This constraint was imposed to preserve the effect of smoothing in the null data. The following methodology was used to determine where edges should be repositioned in order to respect fMRI smoothing and subcortex geometry.

First, the fMRI data for each subcortical voxel was replaced with independent and randomly sampled Gaussian data that was matched in length and amplitude distribution. Second, the Gaussian data was smoothed using the same FWHM as the fMRI data. Third, the Pearson correlation coefficient was computed using the smoothed Gaussian data between all pairs of subcortical voxels. Finally, all voxel pairs were ranked from highest to lowest based on correlation coefficient. At each iteration of the rewiring algorithm, the edge being rewired was repositioned to the highest-ranking pair of voxels that had not yet received a rewired edge. To ensure that the rewired graphs were not fragmented, the first *N* − 1 edges were repositioned according to a minimum spanning tree (MST), where the MST was computed for a lattice graph in which each node corresponded to a subcortical voxel. Edges in the rewired graphs inherited their edge weights from *W*. An ensemble of 100 rewired graphs and corresponding sparse adjacency matrices 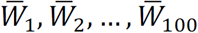 were generated by repeatedly initializing this rewiring procedure with randomly sampled Gaussian data. We found that repositioning edges to randomly chosen pairs of spatially neighbouring voxels yielded virtually identical null data.

Supplementary Fig S4a shows the graph corresponding to *W* and 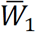 for an axial slice that exemplifies the functional boundary between GP and putamen. Very few edges in the graph for *W* intersect this boundary, whereas the graph for 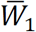 comprises numerous rewired edges that interconnect GP and putamen. Consistent with anatomy, this suggests that the strong gradient in functional connectivity that yields the boundary between these two regions is most likely not due to chance or the effect of smoothing and/or subcortex geometry. Supplementary Fig S5b shows an example that demonstrates the importance of preserving geometry in the null data. In the bow-tie shaped region comprising this example, binary edges are randomly positioned between neighbouring pixels comprising the bow tie. This positioning of edges is consistent with the null hypothesis of a spatially homogenous gradient magnitude. However, the bow knot is the location of a geometric constriction, and if this geometry is not preserved in the null data, a specious boundary between the left and right sides of the bow is suggested.

Laplacian eigenmaps and gradient magnitude images were computed for each of 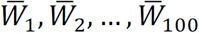 by following the same procedure described above for *W* (see Functional connectivity and eigenmaps). An ensemble of 100 diversity curves were then computed by projecting each of these gradient magnitude images onto the original streamlines that were computed with respect to *W*. These diversity curves were consistent with the null hypothesis and we refer to them as *null diversity curves* (Fig 2c, d).

### Boundary delineation

Statistical inference was performed to determine whether each local maximum in the gradient magnitude images was sufficiently large to warrant delineation of a discrete functional boundary. If the null hypothesis could not be rejected and thus delineation was not warranted, connectional topography was deemed to be most parsimoniously represented as a continuum. The decision to delineate a boundary or not was informed by statistical testing of the null hypothesis performed on both the gradient magnitude images and the diversity curves onto which these images were projected. All local maxima in the gradient magnitude images were considered as candidate boundaries, but candidate boundaries were delineated only if the null hypothesis was rejected; that is, the gradient magnitude was larger than expected due to chance and the effects of smoothing and subcortex geometry. In particular, a p-value for each local maximum on a diversity curve was estimated as the proportion of null diversity curves with local maxima that: i) exceeded or equalled the observed maximum; and, were located within the same vicinity as the observed maximum. The false discovery rate (FDR) was controlled at a threshold of 5% across the set of all maxima. The decision to delineate a boundary was supported for maxima with p-values that survived FDR correction. Figure 2c, d shows examples of diversity curve maxima for which the null hypothesis was rejected (denoted by blue asterisks).

The null hypothesis could also be tested with respect to the gradient magnitude images. In this case, the Kolmogorov-Smirnov (KS) test was used to assess the null hypothesis of equality in the distribution across voxels between the observed gradient magnitudes and the null gradient magnitudes. The null hypothesis was rejected if the tail of the distribution of gradient magnitudes was longer in the observed data, compared to the null data. The main reasons for reverting to the KS test at finer scales were that: i) generally only one candidate boundary was evident within each region comprising Scales II-IV (Supplementary Fig S23); and, ii) we found that mapping diversity curves for relatively small regions did not provide substantial additional insight. Diversity curves and gradientography yield most insight when applied across a relatively large expanse that comprises multiple candidate boundaries (i.e. Scale I). Supplementary Fig S23 shows examples of using the KS to determine whether or not to delineate a candidate boundary in two specific subcortical regions. A strong gradient is evident in the anterior putamen (panel a, yellow colored strip), and the KS test indicates that the gradient magnitude distribution across the voxels of this region (green curve) is significantly longer tailed than the null data (gray curves, *p*<0.01). Therefore, a boundary was delineated that resulted in parcellation of the anterior putamen into one dorsal and one ventral component. In contrast, the null hypothesis could not be rejected with the KS test for the NAc-shell (panel b, *p*=0.22) and thus subdivision of this region was not warranted, despite the presence of a candidate boundary (yellow strip).

The final decision to delineate a boundary was not exclusively determined by whether the null hypothesis could be rejected or not. Other criteria that informed this decision were:

i. Size criterion: A candidate boundary was not delineated if at least one of the regions resulting from boundary delineation would have comprised fewer than 100 voxels, irrespective of whether the null hypothesis was rejected.
ii. Prior knowledge: A candidate boundary that corresponded with established subcortex anatomy or cytoarchitecture was delineated, irrespective of whether the null hypothesis was rejected.
iii. Inter-hemispheric homologue: A candidate boundary was not delineated if the null hypothesis could not be rejected for the corresponding contralateral boundary. Edges in the rewired graphs (null data) inherited their edge weights from *W*, which was not necessarily symmetric. Therefore, the null data was not symmetrized, and thus discrepancies in statistical inference were possible between contralateral boundaries.

These criteria were only ever used for three boundaries of the more than 50 decisions pertaining to boundary delineation (Supplementary Fig S6). In particular, the size criterion prevented further segmentation of a region of the caudate (CAU-DA), the prior knowledge criterion was used to delineate the thalamus-hippocampus boundary at Scale I, despite failure to reject the null hypothesis, and the homologue criterion was used for the medial amygdala (mAMY) at Scale II. These criteria were not used for any other cases and all other boundary delineations were supported by the model selection procedure.

### Hierarchical parcellation

The subcortex was parcellated into distinct functional regions in a hierarchical manner. Each successive scale of the hierarchy involved delineation of additional boundaries that conveyed finer details in connectional topography. Foremost, boundaries were delineated based on maxima in the dorsal and ventral diversity curves for which the null hypothesis was rejected. This resulted in a subcortex parcellation comprising 8 bilateral regions that recapitulated well-known subcortical regions and established anatomical boundaries (Fig 3a). These 8 bilateral regions defined Scale I of the hierarchical parcellation (16 regions in total). Scale I is not intended to provide a novel parcellation, but rather establishes the validity of the model selection procedure and represents a necessary first step to delineate the novel Scale II-IV atlases at finer scales. The next scale of the hierarchy (Scale II) was delineated by separately computing Laplacian eigenmaps and gradient magnitudes for each of the 16 regions using the same procedure that was initially applied to the whole subcortex. This process was repeated in a recursive manner until the null hypothesis could not be rejected for any candidate boundaries, a condition that was satisfied at Scale IV of the hierarchy (Fig 3b & Fig 4). A given region in Scale *k* appeared as either two or more subregions in Scale *k +* 1, if the null hypothesis was rejected, otherwise, it appeared unaltered. For each scale, the false discovery rate (FDR) was controlled at 5% across the family of null hypothesis tests performed for all candidate boundaries. Unlike alternative multiple testing corrections, the FDR is scalable and adaptive^87^, meaning that the greater number of candidate boundaries tested at Scale IV, compared to Scale I, did not make FDR correction at Scale IV inherently more stringent. Note that FDR was performed independently for each scale. Successive scales of the parcellation hierarchy conveyed finer boundaries and more regions (Scale I: 16 regions, II: 32 regions, III: 50 regions, IV: 54 regions). The diameter of all regions comprising Scale IV substantially exceeded the FWHM of the smoothing kernel width (min: 10.2mm, max: 24.4mm, median: 17.2mm). While exceedance of the FWHM is a heuristic criterion, this suggests that the extent of smoothing was sufficiently constrained to enable delineation of the smallest region.

For Scale I, the null hypothesis was tested by performing inference on maxima in the dorsal and ventral diversity curves. In particular, inference was performed to determine whether each maximum in the gradient magnitude was significantly larger than expected due to chance, geometry and smoothing (see Boundary delineation). For Scale II and beyond, as described above, the KS statistic was used to test the null hypothesis, due to challenges in mapping diversity curves for the relatively small regions comprising these scales. Delineation of small regions was also more susceptible to the effects of noise, especially signal contamination from surrounding white matter and cerebrospinal fluid. To reduce these effects, for Scale II and beyond, the adjacency matrix (*W*) used to compute Laplacian eigenmaps was thresholded using the disparity filter^88^. The significance level for the disparity filter was set to the minimum value required to yield a graph that was not fragmented.

#### Watershed algorithm

The watershed transform algorithm^89^ was used to cluster voxels and thereby identify spatially contiguous regions that resulted from delineation of a boundary, as applied elsewhere to parcellate the cortex^3^. The watershed transform was applied to the three-dimensional gradient magnitude images. To initiate the watershed, seed voxels were first chosen, such that exactly one seed voxel resided on each side of a boundary. Seed voxels were approximately positioned to reside within a regional minimum of the gradient magnitude image. A regional minimum that was distant from the boundary was preferred, although the parcellation was robust to the precise seed voxel location. Clusters were then iteratively grown for each seed region until locations were reached where voxels could no longer be unequivocally assigned to one of the clusters. Clusters were therefore analogous to water catchment basins (i.e. minima in the gradient magnitude images), while boundaries represented watershed ridge lines (i.e. maxima in these images). Before applying the watershed transform, the gradient magnitude images were: i) normalized to ensure that all values resided on the unit interval; and, ii) morphologically reconstructed to ensure that regional minima were only present at the location of the chosen seed voxels. The latter step aimed to enhance the contrast between voxels within the expected cluster and boundary voxels. The output of the watershed transform was a cluster of voxels for each seed voxel. Voxels that resided at boundary locations where not necessarily assigned to any cluster, yielding empty space between parcellated subregions. To fill the empty space, for each unallocated voxel, the shortest path was computed to each seed voxel. Shortest paths were computed with respect to the sparse adjacency matrix (*W*), following a strength-to-distance remapping of the edge weights. Unallocated voxels were then assigned to subregions with the minimum shortest path distance.

### Parcellation validation and replication

#### Homogeneity estimation

A valid functional parcellation should comprise regions that are internally homogenous with respect to the functional connectivity profile of each constituent voxel. The regional homogeneity of Scale I-IV parcellations was compared to the extent of homogeneity achieved with alternative subcortex parcellations as well as the homogeneity that could be expected due to chance.

The fMRI data for a region was represented as a matrix of dimension *N* × *T*, where *N* and *T* denote the number of voxels comprising the region and the number of time frames, respectively. *Regional homogeneity* was defined as the variance explained by the first principal component of this matrix, which was estimated using PCA. Homogeneity thus quantified the extent of synchrony in fMRI activity across voxels comprising a region. While homogeneity can be computed with respect to functional connectivity^3^, we performed estimation directly on the fMRI time series acquired in the second session (REST2). This ensured that validation was performed using independent data and an independent quantity. Homogeneity was computed separately for each region and each individual. For each individual, homogeneity was then averaged across all regions to yield an overall estimate of homogeneity for the parcellation as a whole, referred to as *parcellation homogeneity* (Supplementary Fig S11a).

Parcellation homogeneity increased from Scale I to IV (Fig 5a, red dotted line). This was because small regions were inherently more homogeneous than large regions^3^. To determine whether the observed homogeneity values were larger than expected due to chance, the subcortex was parcellated randomly, such that the total number of regions and the distribution of region size were matched between the random and observed parcellations (Supplementary Fig S11b). Parcellation homogeneity was then estimated for 100 such random parcellations to yield a null distribution for homogeneity under the null hypothesis of randomly delineated regions. A p-value for this null hypothesis was estimated as the proportion of random parcellations with homogeneity exceeding or equalling the observed homogeneity. Parcellations were deemed homogeneous if the null hypothesis was rejected (*p*<0.05). Random parcellations were generated by: i) randomly placing seed voxels throughout the subcortex mask in such a way that the Euclidean distance between all pairs of seed voxels was approximately maximized; and, ii) assigning all non-seed voxels to their closest seed voxel in terms of Euclidean distance. This assignment process defined a set of randomly delineated regions. The number of seed voxels was set to the number of desired regions. Parcellations were discarded if they comprised any regions that were not spatially contiguous or not matched with respect to the distribution of region sizes. Discarded parcellations were replaced until 100 valid random parcellations were generated.

Parcellation homogeneity was also computed for existing subcortex parcellation atlases (Supplementary Table S3). The scale (I-IV) that comprised the same, or fewer parcels than the existing parcellation was chosen for comparison. Choosing a scale with fewer parcels advantaged the existing parcellations. We specifically evaluated the homogeneity of two existing parcellations atlases for the whole subcortex^8, 40^ as well as parcellation atlases for hippocampus^24, 42, 90^, thalamus^44^ and striatum^43^. Cerebellar and brainstem regions were omitted from the parcellation delineated by Ji et al^8^ and Fan et al^40^ to ensure homogeneity estimates were specific to the subcortex and therefore comparable to the other parcellations. To facilitate a fair comparison across parcellations comprising a disparate number of regions, observed parcellation homogeneity values were normalized with respect to the mean homogeneity across the set of random parcellations. Ratios exceeding one indicated that parcellations were more homogeneous than expected due to chance.

#### 7T Replication

High resolution fMRI acquired at ultra-high magnetic field strength (7T) can enhance functional contrast-to-noise ratios, improve spatial localization and alleviate partial volume effects^26^. We sought to test the extent to which parcellations delineated using 3T-rfMRI could be replicated using 7T-rfMRI. To this end, the subcortex was hierarchically parcellated by applying the same procedures described above to the 7T-rfMRI data. This involved mapping Laplacian eigenmaps and gradient magnitudes, null hypothesis testing for each maximum comprising the gradient magnitude images and application of the watershed transform to delineate subregions. This yielded a 7T parcellation atlas of the subcortex that also comprised four scales (Scale I: 16 regions, II: 34 regions, III: 54 regions, IV: 62 regions).

To enable comparison between 3T and 7T parcellations, the 3T parcellations were up-sampled to the same spatial resolution as 7T (1.6mm isotropic) and the spatial similarity between each scale of 3T and 7T parcellations was estimated using the normalized mutual information (NMI)^91^. NMI is an information theoretic measure that quantifies the extent to which knowledge of a parcellation reduces the uncertainty in the location of regions in a second parcellation. NMI ranges between zero and one, with higher NMI indicating greater spatial correspondence between parcellations.

### Personalized parcellation

Following Glasser and colleagues^2^, a machine learning approach was used to personalize the group-consensus parcellations and distinguish individual differences in functional organization, yielding a unique parcellation for each individual. This enabled quantification of the extent of variation among individuals for each region. The subcortex mask remained fixed for all individuals, and thus only variation in functional boundaries within the subcortex was permitted, without variation in the overall subcortical volume. The periphery of each region comprising the Scale IV parcellation was first dilated by approximately 4-6 mm to define a zone of uncertainty around each region. The uncertainty zone enabled flexibility in the location of boundaries for individuals who exceeded the group-consensus boundary. The extent of dilation was such that each region in the group-consensus atlas comprised approximately the same number of voxels as its uncertainty zone, meaning that a voxel was equally likely to be randomly classified as belonging to the region, compared to its uncertainty zone. Enlarging the uncertainty zone would have permitted individual boundary locations to deviate to a greater extent from the group-consensus boundaries, although major deviations were not expected in young and healthy adults, consistent with previous work on the cortex^2^. A binary support vector machine (SVM) classifier was trained to classify whether a voxel resided in the region or its uncertainty zone. We use *M* to denote the combined number of voxels comprising the region and its uncertainty zone.

Combining REST1 and REST2 would potentially yield more accurate estimates of functional connectivity compared to any of the individual sessions. However, given that parcellation was performed using data from the first 3T-rfMRI session (REST1), to assess inter-session reproducibility and minimize circularity, the SVM was trained and evaluated using data from the second session (REST2). One hundred individuals were randomly selected to train the SVM and the trained SVM was then applied to delineate a personalized parcellation for the remaining 921 individuals. It is important to remark that each individual contributed *M* training samples because classification was undertaken on a per-voxel basis. The SVM feature space was defined over the symmetric *N* × *N* similarity matrix, where each cell in this matrix stored the similarity in functional connectivity fingerprints between a pair of subcortex voxels (see Functional connectivity and eigenmaps). The feature vector for a particular voxel was selected as the relevant row in this matrix. The feature space was mapped separately for each individual and required computation of individual similarity matrices. A radial basis function kernel was used, and the feature space was standardized.

For each of the 921 individuals in the test set, the SVM was used to predict the likelihood (posterior probability) of each voxel belonging to the region under consideration, as opposed to belonging to the region’s uncertainty zone. See Supplementary Fig S17 for a schematic of this binary classification problem. The threshold for the posterior probability of a voxel belonging to the region was varied between 0 and 100%, where a threshold of 0% led to all voxels, including those in the uncertainty zone, being classified as belonging to the region. Increasing the threshold eventually resulted in loss of spatial contiguity. The extent to which the threshold could be increased while preserving these properties provided insight into parcellation robustness. This process was repeated separately for each region comprising the Scale IV parcellation, requiring an SVM to be independently trained for each region. It was possible for voxels to be included in the uncertainty zones of multiple regions. A voxel could therefore be assigned a probability of belonging to each of many regions. To delineate a personalized parcellation for each individual, a winner-takes-all rule was used to break ties, such that voxels were allocated to regions with the highest probability. Crucially, if the highest probability region did not exceed the posterior probability threshold, the voxel remained unassigned. This introduced the possibility of certain regions in the group-consensus parcellation being absent from an individual’s personalized parcellation. The *regional detection rate* was the proportion of individuals for which a given region was allocated at least one voxel. The detection rate of each region was evaluated as a function of the posterior probability threshold. The group-consensus parcellation was assumed to provide a valid representation of individuals for regions with high detection rates.

The spatial correspondence between the group-consensus and personalized parcellations was also evaluated with the Dice coefficient. In particular, the Dice coefficient was computed separately for each individual and each region, as a function of the posterior probability threshold. If *X* and *Y* denote the indices of the voxels comprising an individual’s personalized region and the group-consensus region, respectively, the Dice coefficient was computed as 2|*X* ∩ *Y*|/(|*X*| *+* |*Y*|η. An average Dice coefficient was then computed for each individual by averaging the regional Dice coefficients.

### Task-evoked connectional gradients

We aimed to investigate whether the connectional topography of the subcortex would exhibit significant variation between task conditions and rest^29, 92-94^. The tasks considered were emotion processing, gambling, language processing, motor execution, relational matching, social inference and working memory. Details pertaining to each task condition are described elsewhere^30^.

Following the same procedure described above (see Functional connectivity and eigenmaps), group-consensus eigenmaps and gradient magnitude images were computed for each of the seven tasks. For each task, the gradient magnitude image for Gradient I was projected onto the dorsal and ventral streamlines. This yielded diversity curves for each task (Supplementary Fig S19a). Tractography of the task-derived gradients was not performed and streamlines determined previously in rest (See Diversity curves) were used as references. To facilitate comparison of diversity curves between different tasks, only the 725 individuals (360 males) that completed all nine conditions (two rest and seven tasks) were included. The mean diversity curve across the nine conditions was first computed and regressed from the diversity curves for each of the nine conditions. This eliminated the common variance across the conditions. The Pearson correlation coefficient was computed between the resulting residuals to generate a correlation matrix of dimension 9 × 9, where each element corresponded to a pair of task conditions. Permutation testing was used to determine a p-value for each correlation coefficient. Specifically, for each pair of comparisons, the points on one of the diversity curves were randomly permutated and the correlation was recomputed on the permuted diversity curves. This was repeated for 10,000 permutations, yielding a null distribution for the correlation coefficient. The p-value was given by the proportion of permutations with the correlation coefficient that exceeded or equalled the correlation coefficient in the unpermuted data. Bonferroni correction was controlled at 0.05/36=0.0014 across the set of all unique pairs of conditions. The pairs of conditions that survived correction are shown in Supplementary Fig S19b and visualized as a graph to show similarities in task conditions (Supplementary Fig S19c). Further, a matrix of dimension *H* × 9 was then constructed, where *H* denotes the number of points along the diversity curve and each column corresponds to one of the nine conditions. Each cell in the matrix stored the residual value of each diversity curve point. Rows and columns were reordered according to average-linkage clustering based on the Pearson correlation coefficient (Supplementary Fig S19d).

Given that substantially fewer fMRI volumes were acquired during task compared to rest (see Data and pre-processing), supplementary analyses were undertaken to test whether task-related changes in connectional gradients could be attributed to the acquisition of fewer volumes. Specifically, the group-consensus eigenmaps and the gradient magnitude images were recomputed using only the first quarter of time frames (T=278) from each run of the resting-state fMRI data (278 × 2 in total), thereby matching the average number of time frames across the seven tasks. We found that the gradient magnitude images computed using the shortened and full-length resting-state fMRI acquisitions were highly correlated (Spearman correlation: *r*>0.98 for both REST1 and REST2). This suggests that differences in the number of volumes between the task and resting-state fMRI acquisitions do not explain the task-evoked reconfiguration of connectional gradients and diversity curves.

### Relating subcortical functional network to behaviors

Individual variation in functional connectivity associates with human behaviors that relate to cognition, emotion and psychopathology in both health and mental illness^20, 31, 95, 96^. While cortico-cortical connections have been well studied and predict individual variation in human behavior, less is known about the predictive utility of functional connectivity within the subcortex. Our multiscale subcortex parcellation facilitated investigation of associations between intra-subcortical functional connectivity and behavioral phenotypes for each atlas scale.

#### Functional connectivity and modular structure

Subcortical functional connectivity matrices were mapped for each individual and each scale of the parcellation. To this end, representative fMRI time series were determined for each region by averaging across all relevant voxels. Pearson correlation coefficients in regional time series were computed for all pairs of regions and represented in the form of a symmetric functional connectivity matrix. The connectivity matrices were then *r*-to-*z* transformed (Fisher transformation). The rows and columns of the connectivity matrices were reordered to accentuate modular structure, where modular structure was determined with the Louvain community detection algorithm^97^. Separate connectivity matrices were mapped for the first (REST1) and second (REST2) 3T-rfMRI sessions, to provide a discovery and within-sample replication set.

#### Behavioral phenotypes

Behavioral measurements and procedures for the HCP are described in detail elsewhere^30^. Items that tapped alertness, cognition, emotion, sensory-motor function, personality, psychiatric symptoms, substance use and life function were selected for further analyses, subject to the following conditions:

i. Only total or subtotal scores were selected for psychological assessments.
ii. Only raw scores were selected for items with both raw and age-adjusted scores.
iii. Items based on strictly binary responses were excluded.
iv. Items with missing responses in more than 10% of individuals were excluded.

Items relating to task performance in emotional processing, language processing, social recognition, motor execution, working memory, relational matching and reward/decision-making processing were also included. This resulted in a total of 109 behavioral measures for each individual (Supplementary Table S4). Participants with missing responses for any of these 109 items were then excluded, resulting in a final sample comprising 958 individuals (453 males, mean age 28.7±3.7yrs). Two 3T-rfMRI sessions were acquired in all these individuals.

Independent component analysis (ICA) was used to reduce the set of behavioral measures to a set of latent dimensions, with the aim of performing inference on the dimensions rather than the full set of behaviors. Before performing ICA, behavioral items with continuously distributed scores (87 of 109) were normalized with a rank-based inverse Gaussian transformation^98^. Age and sex were then regressed from all items and ICA was performed on the residuals that resulted from this regression. Other confounding factors such as blood pressure, hemoglobin, height and weight were not included due to missing responses for several individuals. ICA was performed using the Icasso package^99^ for MATLAB, which provides an implementation of the FastICA algorithm^100^. Under ICA, the matrix of residuals (behaviors × individuals), denoted with *Y*, was factorized into the product *Y* = *A*^−1^*S*, where *A*^−1^ denotes the mixing matrix (behaviors × components) and *S* denotes the estimated independent sources (components × individuals). The inverse of *A*^−1^ is denoted *A* (components × behaviors) and referred to as the de-mixing matrix.

To estimate the de-mixing matrix, FastICA was performed with default parameter settings, including a cubic nonlinearity and deflationary orthogonalization. Bootstrapping was used to sample individuals with replacement and ICA was performed independently on each bootstrapped sample using randomly chosen initials conditions. A total of 500 bootstrapped samples were generated. Independent components that represented a consensus across the 500 samples were then derived using a clustering procedure. In particular, agglomerative clustering with average-linkage was used to partition the de-mixing matrix into putative clusters. The number of dimensions in the feature space was the number of behaviors and each cluster comprised groups of components (i.e. rows from *A*). Clearly separated clusters indicated that independent components were consistently and reliably estimated, despite randomization of initial conditions and bootstrapping. A cluster quality index was used to quantify cluster separation and provided a metric to select the number of independent components (see below). Cluster separation was quantified by the difference in average intra-cluster similarity and averaged inter-similarity, denoted by =*Iq*. A higher = *Iq* suggests compact clusters, whereas a lower = *Iq* suggests that clusters are not well separated. These operations were performed internally in the Icasso package and further details are available in the package’s documentation.

In addition to these internal procedures, we performed a sampling and matching process to ensure stability and reliability of the estimated independent components. Specifically, the ICA as described above was repeated for 10 trials, yielding an ensemble of de-mixing matrices, *A*_1_, *A*_*2*_, …, *A*_10;_ and corresponding independent components, *S*_1_, *S*_*2*_, …, *S*_10_. Variability in *A*_*u*_ and *S*_*u*_ across trials was due to randomization of initial conditions and random sampling inherent to bootstrapping. Each row of *A* corresponded to an independent component and each column quantified the weighting of a behavioral item in the given component. We sought to evaluate the similarity of the resulting estimates across trials, to investigate whether the Icasso procedure yielded reproducible independent components across the 10 trials. For each pair of trials *u* and *v*, a correlation matrix of dimension *I* × *I* was computed, where *I* denotes the number of independent components. Element (*i, j*); of this correlation matrix stored the Pearson correlation coefficient between row 1 of *A*_*u*_ and row *j* of *A*_*v*_. We denote the correlation matrix between trial *u* and *v* with *C*_*uv*_ *u, v* = 1,2, …, 10. Given that independent components are resolved up to an arbitrary sign flip, the absolute value of all correlation coefficients was considered. The *n*th component in trial *u* did not necessarily correspond to the *n*th component in trial *v*, and thus a one-to-one matching of components was required for each pair of trials. The Hungarian algorithm^101^ was applied to *C*_*uv*_ to determine a one-to-one matching that maximized the correlation coefficients between the matched pairs of components. If ICA estimates were reproducible between trials *u* and *v*, each column of *C*_*uv*_ should comprise a correlation coefficient that is substantially larger than all others in the column, indicating an obvious matching. The trial that was best matched to all other trials was chosen as the *reference trial*, where matching quality was quantified as the average correlation coefficient computed over all matched pairs of components in *C*_*uv*_. Once the reference trial was determined, the components in each of the other trials were reordered to match the reference, as dictated by the optimal one-to-one matching computed with the Hungarian algorithm. The polarity of the component was potentially flipped to ensure consistency with the reference trial. The reordered de-mixing matrices *A*_1_, *A*_*2*_, …, *A*_10_ were then averaged across trials to yield a single de-mixing matrix that represented a consensus across the 10 trials. The independent components *S*_1_, *S*_*2*_, …, *S*_10_ were reordered and then averaged in the same way.

Due to the effects of bootstrapping and randomization, the de-mixing matrix was not necessarily perfectly orthogonal and some components were minimally correlated. The Pearson correlation coefficient was computed between all pairs of components in the de-mixing matrix. The reciprocal of the largest correlation coefficient was used to quantify the extent of interdependence between components, such that higher values of the reciprocal were desired. This was performed for each re-ordered de-mixing matrices *A*_1_, *A*_*2*_, …, *A*_10_ and the average across trials resulted in a consensus index of the independence.

Three criteria were therefore used to evaluate ICA performance and assist in determining the best number of components to estimate:

i. *Cluster separation:* Extent of separation between clusters among independent components found across bootstrapped samples. Estimated internally as part of the Icasso package and provided an estimate of reliability and stability of independent components.
ii. *Trial matching quality:* Quality of the one-to-one matching of independent components across ICA trials. Quantified the reproducibility of independent components and was measured as the average distance between matching independent components of each trial and the reference trial.
iii. *Independence index:* Independence of components in the de-mixing matrix. Quantified as the reciprocal of the largest correlation coefficient between all pairs of components.

The ICA procedure described above, including bootstrap resampling and trial matching, was repeated separately for candidate models ranging from 3 to 30 independent components. The optimal model (i.e. number of components) was informed by the plotting the above measures of cluster separation, trial matching quality and independence index as a function of the number of components and identifying abrupt changes based on the elbow criterion (Supplementary Fig S24). While both cluster separation (panel a) and trial matching quality (panel b) showed abrupt reductions when moving from six to seven components, suggesting six components was optimal, the independence index (panel c) suggested that five components was most suitable. We could therefore narrow down model selection from 28 candidates to only two candidate models: five or six independent components. To break the tie between these two models, the independent components comprising both models were visualized as word clouds, with the font size of each behavior scaled according to its de-mixing weight. Although, the five and six component models were highly comparable, we selected the five-component model because it was the simpler model and enabled clear labelling of each independent component. The five independent components characterized cognitive performance, illicit substance use, tobacco use, personality and emotional traits, as well as mental health. Hence, the 109 behavioral items selected initially were decomposed into five independent components, which we referred to as behavioral dimensions (Fig 8a, b & Supplementary Fig S21).

#### Network-based statistic

The network-based statistic^32^ was used to test whether functional connectivity mapped within each hierarchical parcellation scale (Scale I-IV) significantly associated with any of the five behavioral dimensions. The NBS was applied independently to each dimension and parcellation scale. For each pair of regions, a general linear model (GLM) was formulated in which functional connectivity was the dependent variable and the independent variables included an intercept term, the behavioral dimension (i.e. relevant column from the estimated independent sources) and confounds including head motion parameters summarised by the framewise displacement^66^ and total subcortical volume estimated by FreeSurfer (https://surfer.nmr.mgh.harvard.edu/). Given that age and sex were initially regressed from all behavioral scores before performing ICA, these potential confounds were not included in the statistical model. Each connection was endowed with a t-statistic and corresponding uncorrected p-value for the null hypothesis that the association between functional connectivity and the behavioral dimension was not significantly different from zero. The premise of the NBS is to reject the null hypothesis for connected graph components^32^, rather than for individual connections, thereby relinquishing some precision in effect localization for a gain in statistical power. To this end, connected components were identified among the set of connections with a t-statistic exceeding 4. This t-statistic threshold was chosen to ensure that the smallest effect size of interest that could be detected was approximately a Cohen’s d of 0.1. The number of supra-threshold connections comprising each component defined the component’s size. Permutation testing was used to generate an empirical null distribution for component size. For each of 10,000 permutations, an established method^102^ was used to refit the GLM to data in which the correspondence between individuals and behaviors was permuted. Permutation sequences were obtained with multi-level block permutation^59, 60^ to ensure that the data were shuffled in a way that respected known familial relationships among individuals. In particular, each family could only be shuffled as a whole and family members could only be permuted within their family. Dizygotic twins and non-twin siblings were assumed to share the same kinship and they could therefore be permuted. The size of the largest connected component was stored for each permutation to ensure control of the family-wise error rate across all components identified. The family-wise error corrected p-value for an observed component was given by the proportion of permutations for which the largest component identified was larger than or equal in size to the observed component. The NBS was repeated independently for each of the five behavioral dimensions and for the two alternative hypotheses of positive and negative correlation. This resulted in a total of ten multiple comparisons. The false discovery rate was controlled at 5% across these multiple comparisons.

The NBS was performed on functional connectivity mapped using 3T-rfMRI data from the second acquisition (REST2), given that the new atlas was delineated using the first acquisition (REST1). The reproducibility of any significant associations identified with the NBS in REST2 were evaluated using functional connectivity mapped using 3T-rfMRI data from REST1. To enable comparison, functional connectivity mapped with an alternative subcortex atlas (Subcortex-II)^40^ was also tested for associations with behavioral dimensions following the same procedure described above.

## Data and code availability

Analyses were undertaken on publicly available human neuroimaging datasets acquired and maintained by the Human Connectome Project^19^. These datasets are available at: https://db.humanconnectome.org/. The new atlas is openly available in the form of NIFTI (Neuroimaging Informatics Technology Initiative) and CIFTI (Connectivity Informatics Technology Initiative) files. See Supplementary Table S5 for further details. To facilitate mapping of whole-brain connectomes, we have also integrated the new atlas into several well-known cortex-only parcellation atlases and the combined cortex-subcortex atlases are made openly available. In addition to these key resources, custom MATLAB code to compute Laplacian eigenmaps, gradient magnitudes, diversity curves and other computational analyses undertaken as part of this study are openly available at: https://github.com/yetianmed/subcortex

All additional software packages used for this study are freely and openly available:

Diffusion Toolkit and TrackVis: http://trackvis.org/

NBS: https://www.nitrc.org/projects/nbs/

Icasso: https://research.ics.aalto.fi/ica/icasso/

NeuroMArVL: https://immersive.erc.monash.edu/neuromarvl/

PALM: https://fsl.fmrib.ox.ac.uk/fsl/fslwiki/PALM/ExchangeabilityBlocks

## Acknowledgements

We thank Dr Matthew Glasser for provision of the Wishart filter. We thank Dr Luca Cocchi for provision of the additional validation dataset (Supplementary S3). All other data were provided by the Human Connectome Project, WU–Minn Consortium (1U54MH091657; Principal Investigators: David Van Essen and Kamil Ugurbil) funded by the 16 National Institutes of Health (NIH) institutes and centers that support the NIH Blueprint for Neuroscience Research; and by the McDonnell Center for Systems Neuroscience at Washington University.

