## Supplementary Information for "Topographic organization of the human subcortex unveiled with functional connectivity gradients"

### Supplementary Figures

All anatomical images and renderings are shown in Montreal Neurological Institute (MNI, 6th generation) anatomical reference space. Slice location of images is indicated in MNI coordinates (millimeters). Background images are derived from the MNI152 standard-space T1-weighted average reference image. For visualization of the 7T atlas, the reference image was resliced to the resolution of the 7T functional MRI data.

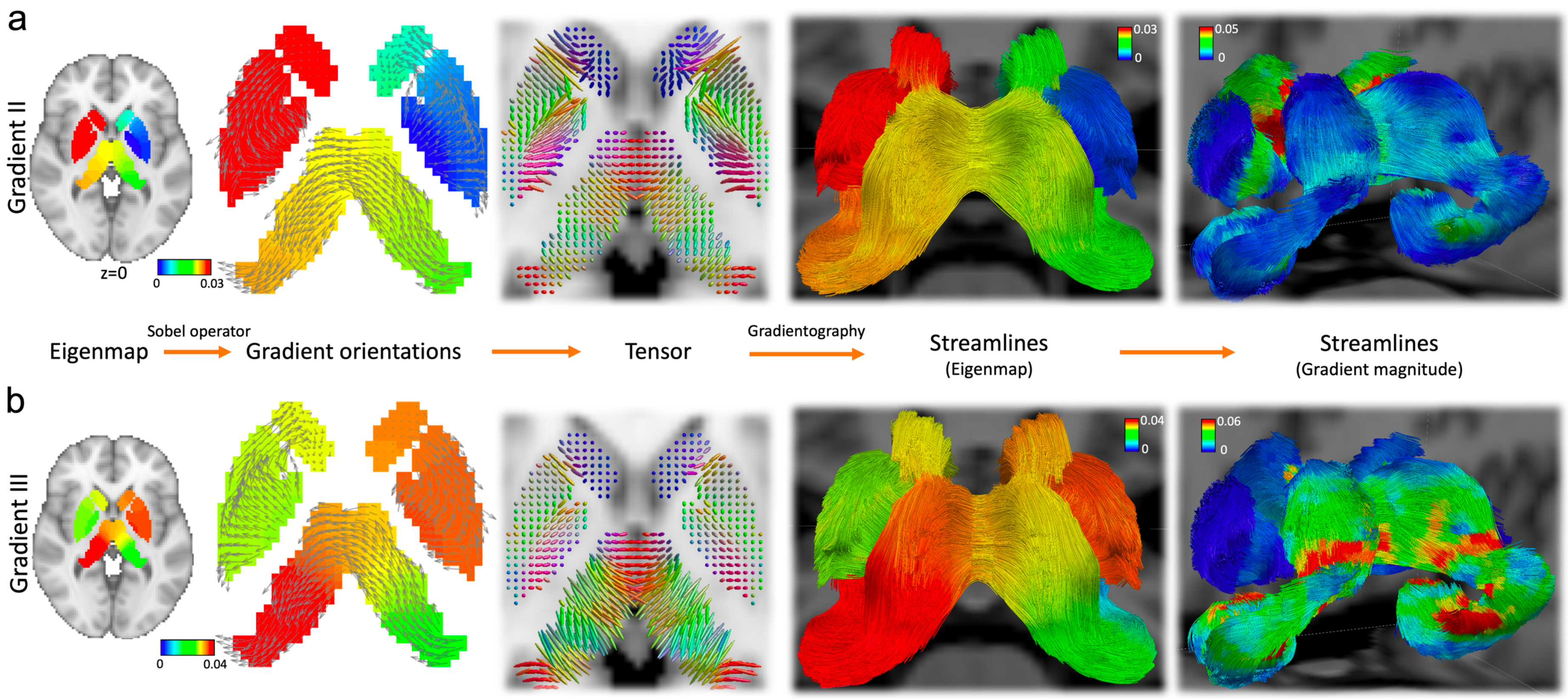

**Fig S1. Mapping of Gradient II and III.** Group-consensus eigenmaps, tensors and gradientography for Gradient II (**a**) and Gradient III (**b**). Gradients were mapped to subcortical voxels to enable anatomical visualization. Eigenmaps for each gradient are shown as axial slices, with a reference structural MRI image used as the background. The Sobel operator was used to estimate the local gradient direction and magnitude for each subcortical voxel. The arrows shown point in the direction of the estimated gradients. Arrow lengths are commensurate with gradient magnitude. Tensors were fitted to the gradient field. Tensors are colored according to gradient direction (blue: superior-inferior, red: left-right, green: posterior-anterior). Long cigar-shaped tensors indicate large gradient magnitudes. Streamlines were propagated through the tensor field using tools for diffusion MRI tractography. Streamlines are colored according to eigenmaps (second from right) and gradient magnitude (rightmost). Local maxima in the gradient magnitude are evident within circumscribed bands along streamlines, indicating putative functional boundaries. While Gradient I characterized an ipsilateral organizational axis with extremes located at amygdala and globus pallidus (see Fig 1b), Gradient II and III maximally differentiated contralateral regions in dorsal (bilateral globus pallidus) and ventral (bilateral amygdala), respectively. Of note, the anterior-posterior or rostrocaudal axis explains much of the spatial variation in cortical microstructure, which is evident in a broad range of the mammalian species, including rodents, marsupials and primates[1-4]. This rostrocaudal organization also aligns with the spatial gradients in neurodevelopment[5, 6].

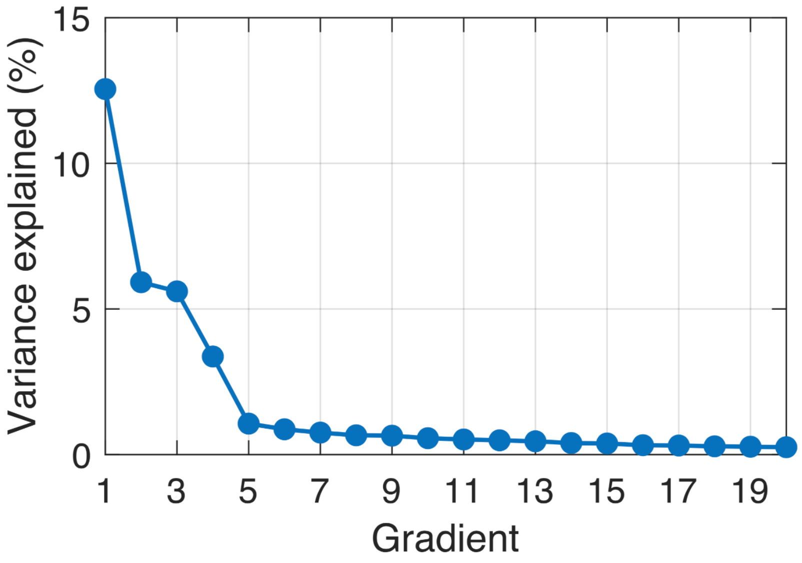

**Fig S2. Variance explained by successive gradients.** Variance explained in subcortical functional connectivity by successive gradients (Laplacian eigenvectors) shown as a function of gradient index, ordered from smallest to largest eigenvalue. The eigenvector associated with the second smallest eigenvalue is labelled Gradient I, while the eigenvector with the third smallest eigenvalue is labelled Gradient II, and so on. The variance explained by the first 20 eigenvectors is shown. The variance explained by Gradient IV and beyond falls below 5%, and thus only Gradients I-III are analyzed in this study.

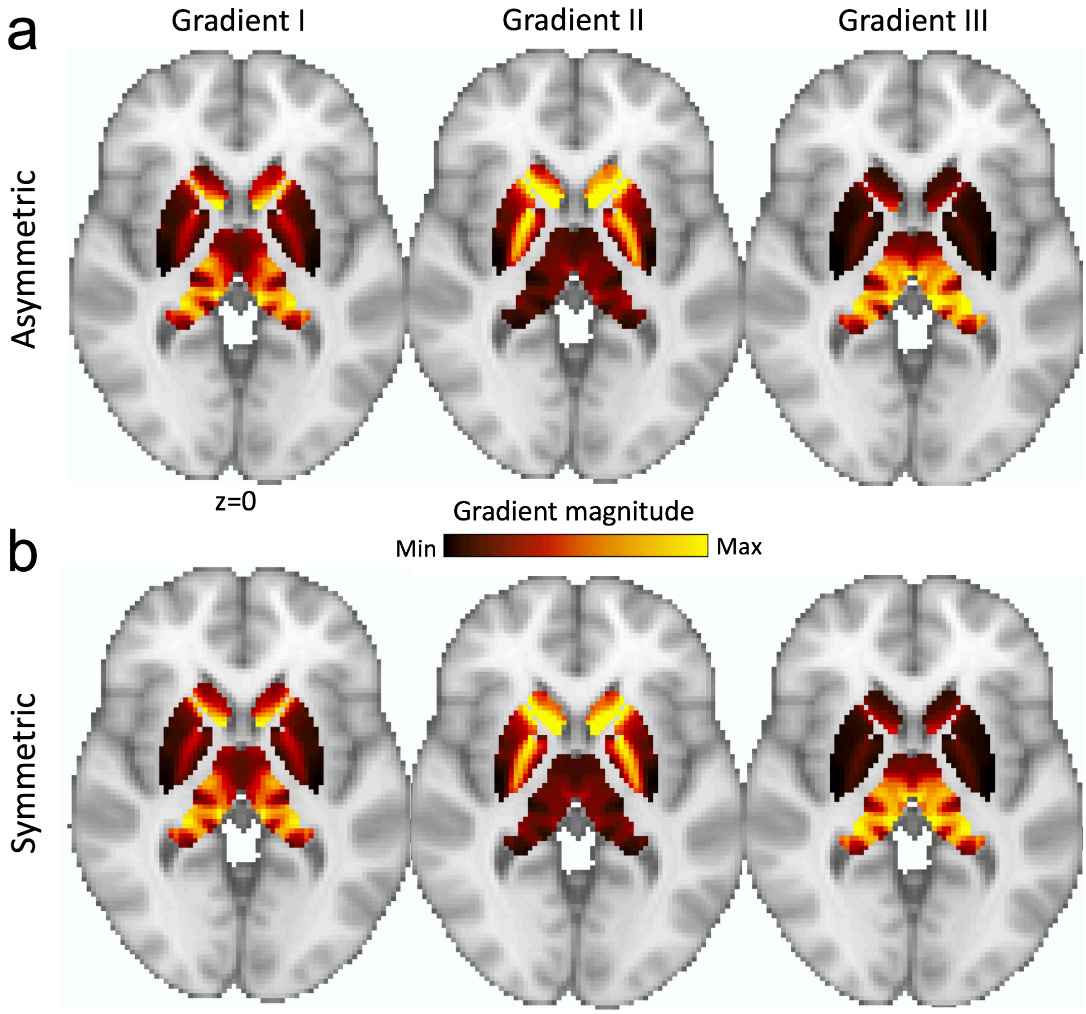

**Fig S3. Impact of left-right symmetrization on gradient magnitude images.** Left-right symmetrization was performed to enhance the signal-to-noise ratio. **a,** Gradient magnitude images for Gradient I-III without left-right symmetrization. Images show a large degree of left-right symmetry, even without explicit symmetrization. **b,** Left-to-right symmetrized gradient magnitude images. Images are shown as axial slices, with a reference structural MRI image used as the background. The same axial slice is used for all images.

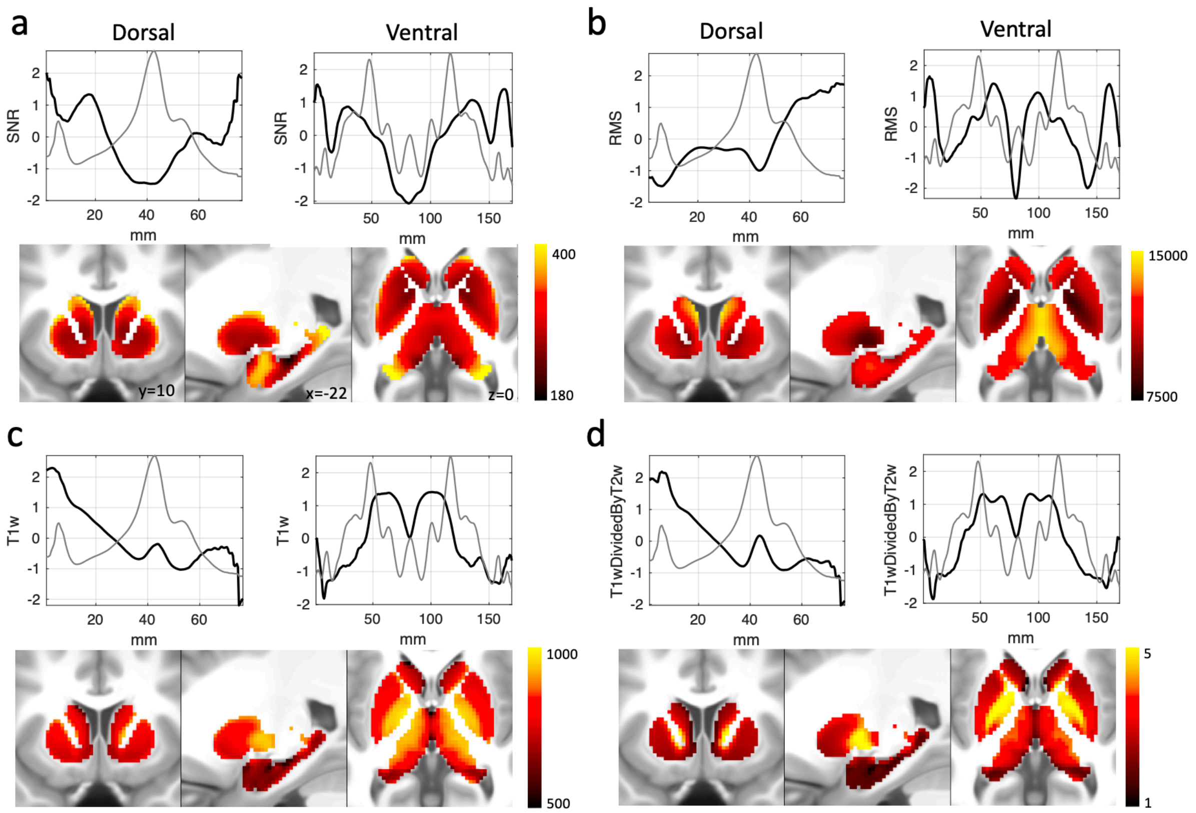

**Fig S4. Spatial variation in primary MRI signal properties in the subcortex. a,** Signal-to-noise ratio (SNR) was computed for each subcortical voxel by averaging the signal intensity across all time frames in each run of the preprocessed fMRI data and normalizing by the standard deviation over time[7]. SNR was averaged across runs for each individual and then averaged across individuals (REST1, n=1080) to yield a group-consensus subcortical SNR map. **b,** Blood-oxygenation level dependent (BOLD) signal magnitude was computed for each subcortical voxel by calculating the root mean squared (RMS) of the signal intensity over time. RMS was then averaged across individuals, yielding a group-consensus subcortical BOLD signal magnitude map. **c,** Group-averaged (S1200) T1-weighted contrast in the subcortex resliced to 2mm isotropic voxel resolution. **d,** Group-averaged (S1200) T1/T2-weighted contrast resliced to 2mm isotropic voxel resolution. The spatial correlation between each of the four group-consensus signal maps and the connectivity gradient magnitude map (Fig 6a) was quantified by the Spearman correlation coefficient (SNR: $r$=-0.21; RMS: $r$=0.42; T1-weighted: $r$=0.02; T1/T2-weighted: $r$=0.02).The four group-consensus signal maps were then projected onto the previously mapped streamlines (Fig 2), yielding dorsal and ventral diversity curves (black) for each of the four signal maps. The diversity curves for each of these four measures do not recapitulate peaks in the gradient magnitude images (light gray), suggesting that spatial variation in structural factors and SNR does not appear to confound the computation of functional connectivity gradients.

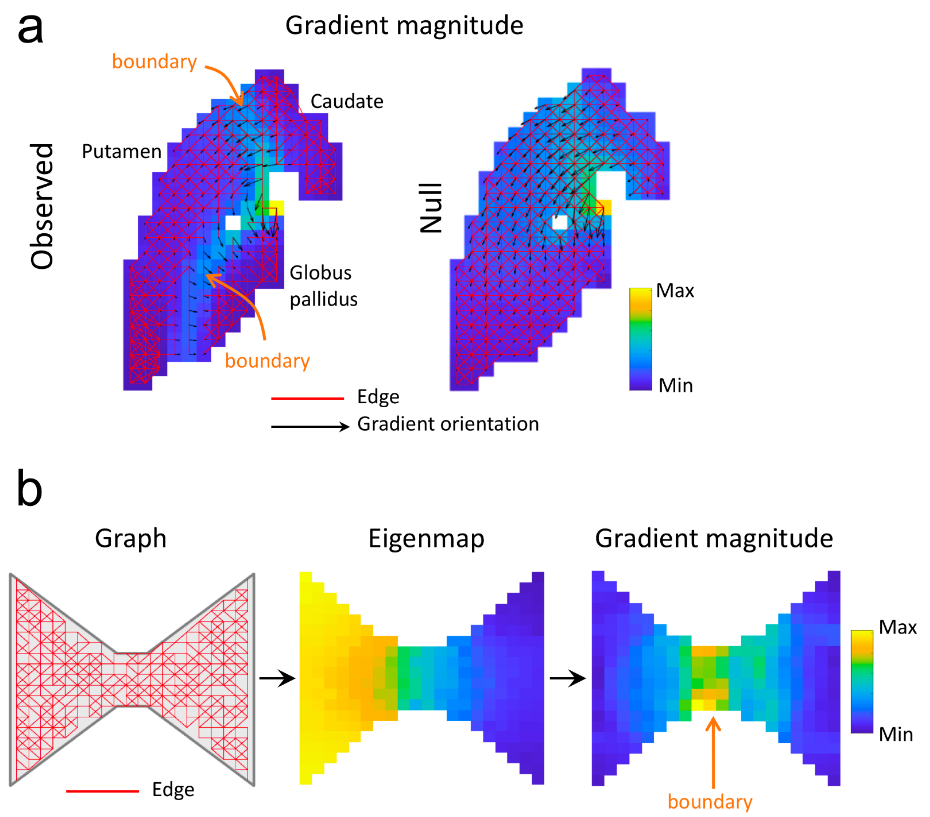

**Fig S5. Testing whether gradient magnitude peaks are large enough to warrant boundary delineation.** Null data was generated to test the null hypothesis that a gradient magnitude peak was due to chance, the geometry of the subcortex and/or other confounds. The alternative hypothesis was that the peak demarcated a discrete boundary in functional connectivity topography. **a,** Graph defined by the sparse adjacency matrix $W$ computed from empirical fMRI data (observed data) is compared to one rewired graph generated from the null data ($\overline{W}_1$). Nodes in the graph correspond to subcortical voxels. Edges (solid red lines) are drawn between voxels that share similar functional connectivity profiles. Axial slices from the gradient magnitude images are shown in the background of each graph. It should be noted that very few edges in the graph for $W$ (observed data) traverse the functional boundary between globus pallidus and putamen (indicated by orange arrows), whereas the graph for $\overline{W}_1$ (null data) comprises a number of rewired edges that cross this boundary. Therefore, unlike the observed data, a peak in the gradient magnitude image between the globus pallidus and putamen is absent in the null data, suggesting that the null hypothesis can be rejected for this particular boundary. Indeed, this was confirmed by generating 100 rewired graphs ($\overline{W}_1$, $\overline{W}_2$,…, $\overline{W}_{100}$) and showing that the proportion of rewired graphs with peaks at this location that exceeded the peak in $W$ did not exceed 5%. **b,** Hypothetical example demonstrating how the effect of geometry alone can intrinsically lead to delineation of spurious boundaries. Edges (solid red lines) were randomly placed between pairs of neighboring pixels residing within a hypothetical bow-tie shaped object (left). Random placement of edges between neighboring pixels is consistent with the null hypothesis of a uniform spatial gradient (i.e. rate of change in gradient is homogenous across the extent of the bow tie). The Laplacian eigenmap (center) and the eigenmap's gradient magnitude (right) is shown for the graph (left). Note the peak in the gradient magnitude image that is located at the knot of the bow tie (indicated by the orange arrow), providing evidence for a putative boundary, despite the null hypothesis being true by design. The gradient magnitude peak is exclusively due to the geometric constriction owing to the knot. Edges are more likely to be placed within the ends of the bow tie, rather than within the knot, because the ends occupy a comparably larger area. This hypothetical example demonstrates the importance of accounting for geometry when generating null data to test whether gradient magnitude peaks are sufficiently large to warrant boundary delineation. Without accounting for the effect of geometry, specious boundaries could potentially be delineated due to the convoluted geometry of the subcortex.

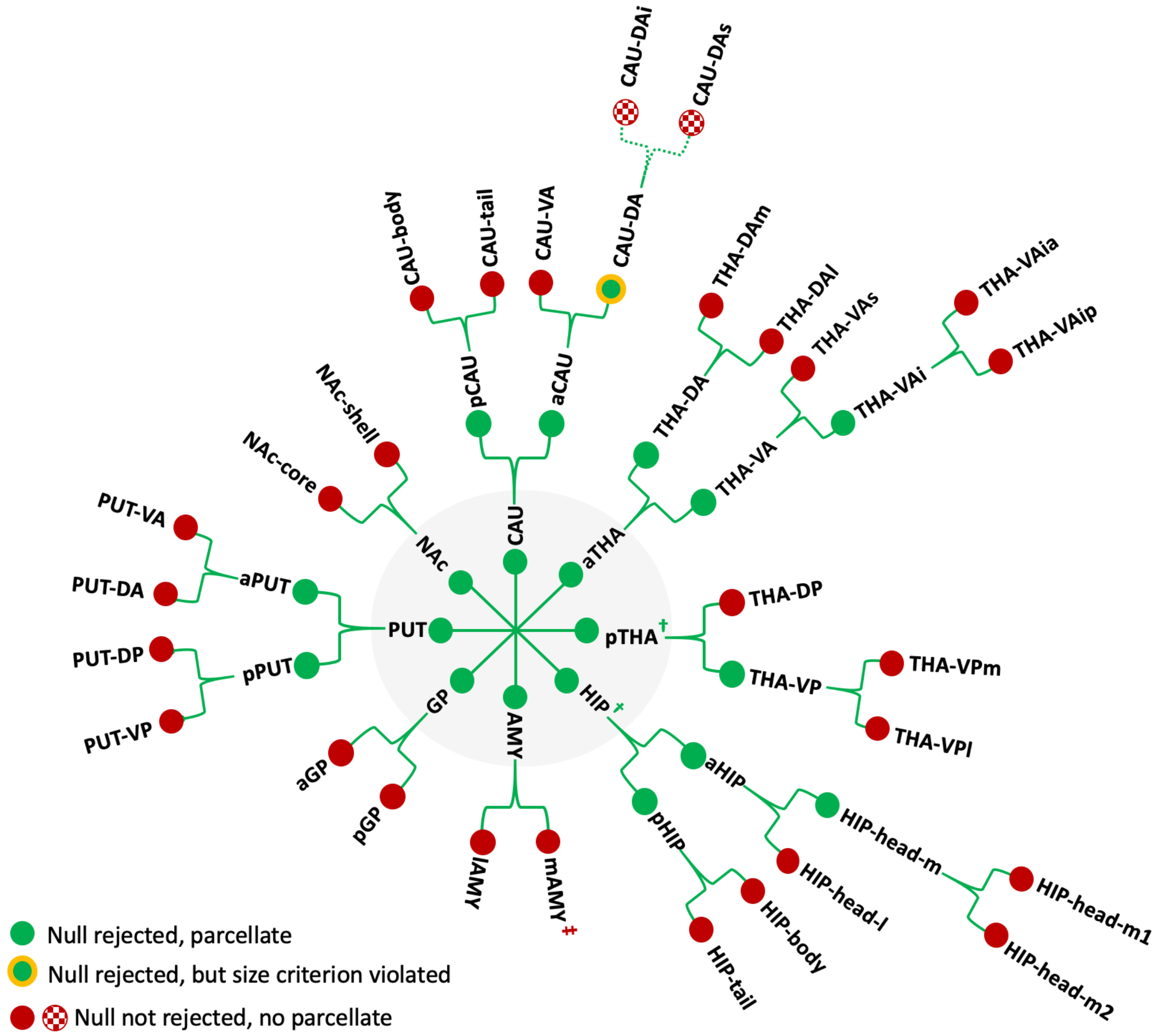

**Fig S6. Criteria used to justify boundary delineation.** Model selection and null hypothesis testing was used to determine whether gradient magnitude peaks were sufficiently large to warrant boundary delineation. This process was repeated recursively for each new region delineated, unveiling a multiscale parcellation architecture. The parcellation hierarchy is visualized in the form of a circular tree. The central node represents the entire subcortex and other nodes represent distinct regions. Regions are arranged within four concentric circles, where the innermost circle (gray) is the first level (Scale I). Nodes are colored according to criterion used to justify further boundary delineation within the region or to terminate the branch of the hierarchy. For the vast majority of regions, null hypothesis testing determined whether or not a boundary was delineated. However, additional criteria were used for three regions (see Methods). Green: null rejected, parcellate; green with yellow circle: null rejected, but size criterion violated and resulting parcels considered too small to delineate (size criterion); red: null not rejected, no parcellate. †: prior knowledge criterion; ‡: inter-hemispheric homologue criterion. If the size criterion were to be relaxed, the dorsoanterior caudate (CAU-DA) would be subdivided into inferior and superior component.

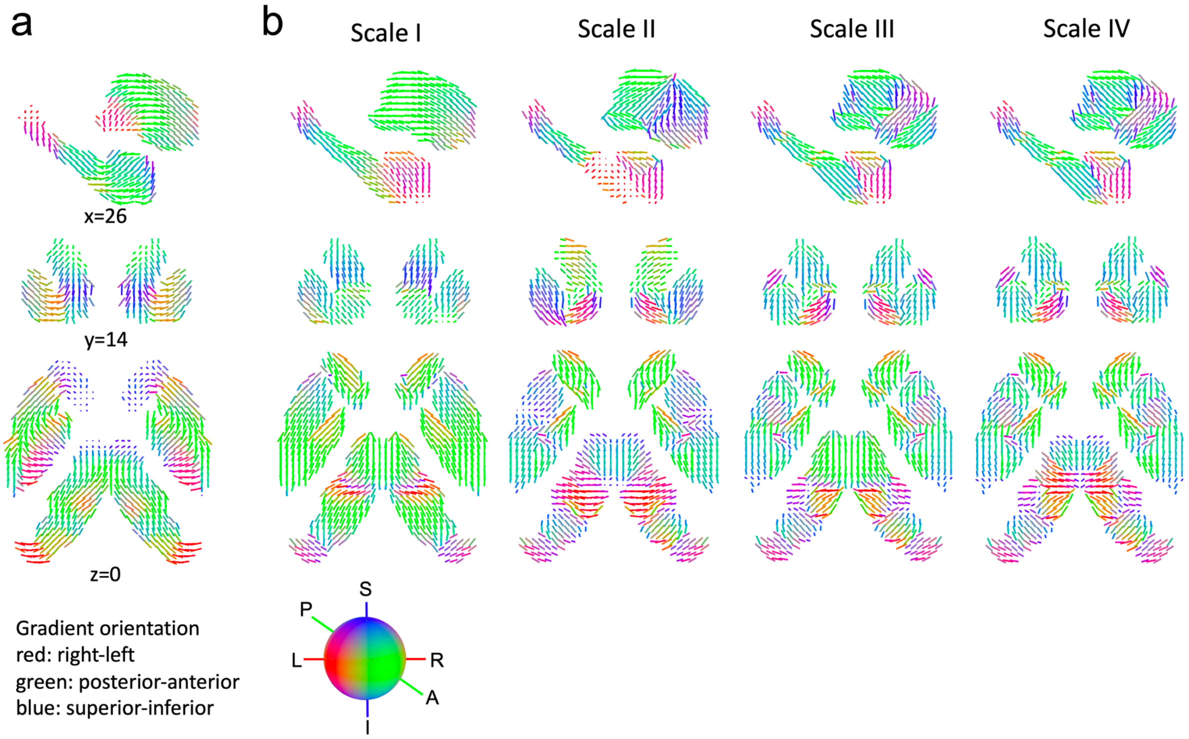

**Fig S7. Organization of connectivity gradients in the human subcortex. a,** The quiver lines represent gradient directions for each voxel. Each quiver is colored according to gradient direction (red: left-right, green: posterior-anterior, blue: superior-inferior) and scaled to unit length. **b,** Gradients are hierarchically organized along the parcellation hierarchy. Local gradient direction in each subcortical voxel was estimated within each region at each atlas scale (Scale I–IV).

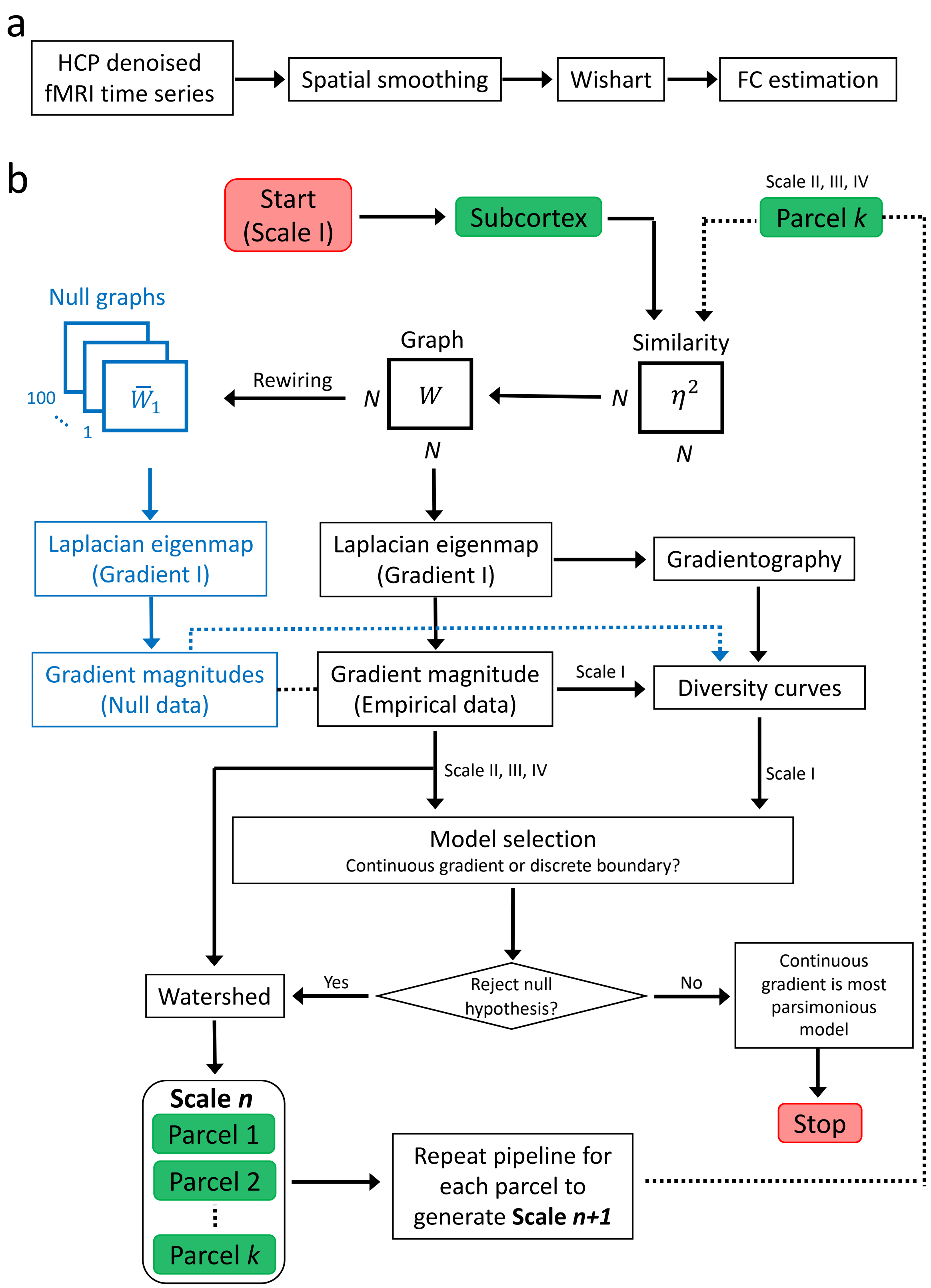

**Fig S8. Schematic of fMRI data preprocessing and boundary delineation. a,** The minimally preprocessed and denoised (ICA+FIX) fMRI data were sourced from the HCP. The preprocessed data were spatially smoothed and then adjusted using a recently developed Wishart filter[8, 9] to further improve the signal-to-noise ratio. The 3T images were smoothed with a Gaussian smoothing kernel of 6mm FWHM (full width at half maximum), whereas

4mm FWHM was used for the 7T images. Using the preprocessed fMRI data, functional connectivity (FC) was then measured between each subcortical voxel and all other cortical and subcortical voxels, as shown in Fig 1a. A similarity matrix was computed for each individual to quantify the extent of similarity in the functional connectivity profiles of all pairs of subcortical voxels. **b,** Boundary delineation pipeline. The subcortex was recursively parcellated into increasingly fine functional subdivisions, until the null hypothesis of a single, continuous region with no discrete boundaries could no longer be rejected, yielding a multiscale characterization of subcortical architecture. Rejection of the null hypothesis indicated that further subdivision of a region was warranted, leading to delineation of a finer scale. Model selection refers to the decision of whether to delineate a discrete boundary or represent spatial variation in functional connectivity as a gradual continuum. The arrow connecting the end of the pipeline to the beginning indicates the recursive nature of the pipeline, where each new recursion designates a finer parcellation scale. The flowchart shows that the entire pipeline (i.e. gradient mapping, null data and model selection) is performed separately for each newly delineated region, until the null hypothesis cannot be rejected. Model selection was performed differently for Scale I compared to the finer scales. For Scale I, where multiple candidate boundaries were observed, local peaks in the gradient magnitude in the diversity curves generated from the empirical data were benchmarked against the diversity curves generated with the null model (blue blocks). Model selection was performed to test whether the peak magnitude was sufficiently large (i.e. exceeded the null distribution) to warrant boundary demarcation. Next, the watershed transform algorithm was used to segment subcortical voxels into contiguous parcels. For finer scales (Scale >1), where regions were small and usually comprised a single candidate boundary, the Kolmogorov-Smirnov (KS) test was used to assess the null hypothesis of equality in the distribution across voxels between the observed gradient magnitudes and the null gradient magnitudes. The null hypothesis was rejected if the tail of the distribution of the gradient magnitude was longer in the empirical data. Other criteria that informed the decision of boundary delineation include: i) minimum size criterion to avoid excessively small regions; ii) prior anatomical knowledge; and, iii) presence of inter-hemispheric homologues. Similarly, the watershed transform algorithm was used to generate subcortical parcels. See Fig 1 for detailed gradient mapping and gradientography pipelines.

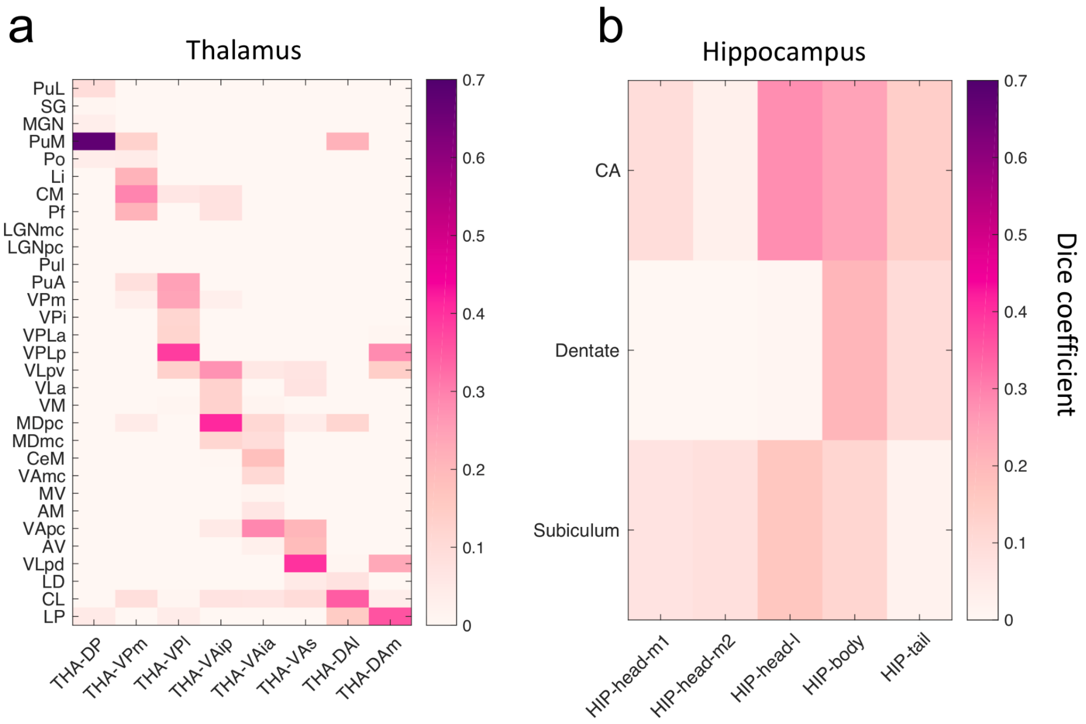

**Fig S9. Spatial correspondence between the new atlas and histological parcellations of thalamus and hippocampus. a,** Dice coefficient is shown between each thalamic anatomical nucleus comprising an existing histological atlas[10] (vertical axis) and each functional thalamic region delineated at Scale IV (horizontal axis). Despite the mismatch in parcellation resolution, several thalamic regions comprising the new atlas can be uniquely mapped to one, or a cluster, of histologically delineated nuclei. For example, THA-DP uniquely maps to PuM, whereas THA-VPm maps to Pf, CM and Li. The spatial correspondence is moderate to good (Dice coefficient: 0.3-0.7), although several histological nuclei cannot be uniquely assigned to a region in the new atlas (e.g. SG). Abbreviations of anatomical nuclei: PuL, inferior pulvinar; SG, suprageniculate nucleus; MGN, medial geniculate nucleus; PuM, medial pulvinar; Po, posterior pulvinar; Li, limitans nucleus; CM, centre median nucleus; Pf, parafascicular nucleus; LGNmc, lateral geniculate nucleus (manocellular part); LGNpc, lateral geniculate nucleus (parvocellular part); PuL, lateral pulvinar; PuA, anterior pulvinar; VPm, ventral posterior medial nucleus; VPi, ventral posterior inferior nucleus; VPLa, ventral posterior lateral nucleus (anterior part); VPLp, ventral posterior lateral nucleus (anterior part); VLpv, ventral lateral posterior nucleus (ventral part); VLa, ventral lateral anterior nucleus; VM, ventral medial nucleus; MDpc, mediodorsal nucleus (parvocellular part); MDmc, mediodorsal nucleus (manocellular part); CeM, central medial nucleus; VAmc, ventral anterior nucleus (magnocellular part); MV, medioventral nucleus; AM, anterior medial nucleus; VApc, ventral anterior nucleus (parvocellular part); AV, anterior ventral nucleus; VLpd, ventral lateral posterior nucleus (anterior part); LD, lateral dorsal nucleus; CL, central lateral nucleus; LP, lateral posterior nucleus. **b,** Dice coefficient is shown between each hippocampal subfield[11, 12] (vertical axis) and each functional hippocampal region delineated at Scale IV (horizontal axis). Unlike the thalamus, spatial correspondence between the histologically defined hippocampal subfields and the new atlas is relatively poor. This is because the subfields are delineated approximately parallel to the longitudinal hippocampal axis, whereas hippocampus is parcellated perpendicular to this axis in the new atlas. CA, cornu ammonis subfield. Dentate, dentate gyrus.

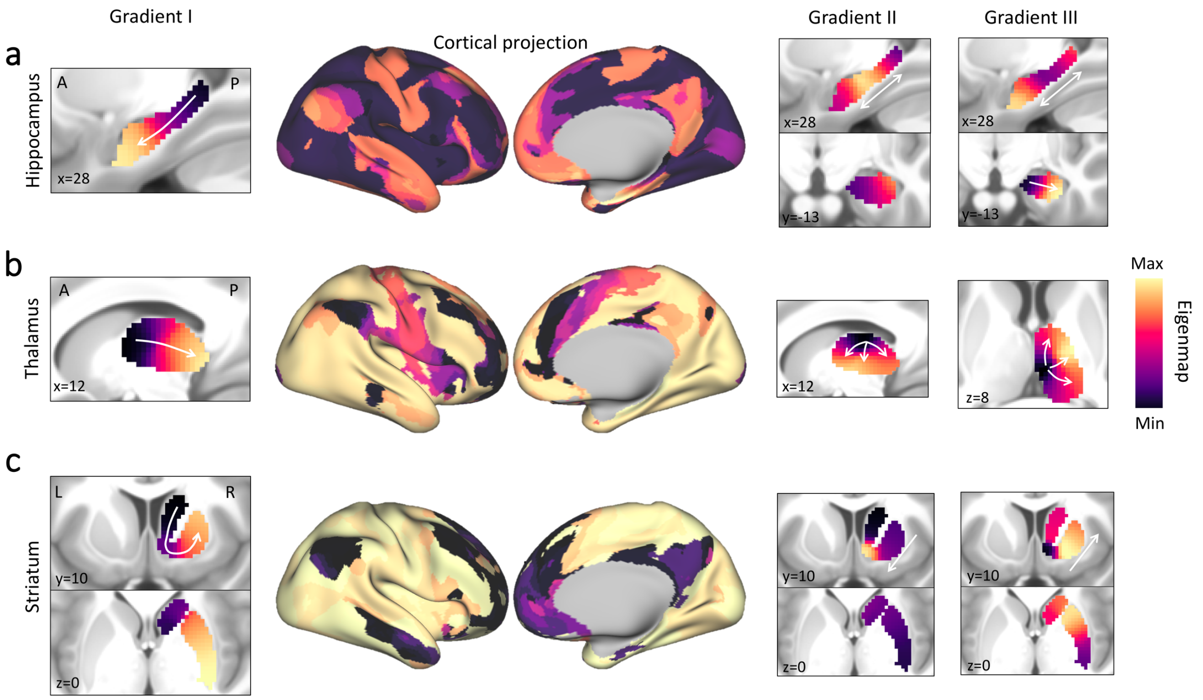

**Fig S10. Projection of gradients within specific subcortical nuclei onto the cortical surface.** Connectivity gradients within hippocampus (**a**) thalamus (**b**), and striatum (**c**) are shown as projections on the cortical surface for Gradient I. Eigenmaps for Gradients II and III are shown in anatomical space. **a,** Cortical surface projections show that progressively more anterior areas of hippocampus project to progressively more ventromedial parts of the frontal and cingulate cortices. **b,** The anterior thalamus is primarily connected to dorsoanterior cingulate, dorsolateral prefrontal, medial superior frontal and anterior insular cortices, which together form the cingulo-opercular network[13, 14]. In contrast, the middle-to-posterior thalamus is widely connected to unimodal and multimodal areas, including somatosensory, motor, premotor, visual and auditory cortices, as well as orbital frontal cortex. **c,** The dominant striatal gradient separates putamen, NAc and caudate, with NAc and caudate differentially projecting to association networks (i.e. default mode[15] and frontoparietal network[16]) along a ventromedial-to-dorsolateral gradient, and putamen widely projects to other parts of the brain. In contrast, the second and third striatal gradients, are organized along a dorsal-to-ventral axis[17-19]. Gradient I (eigenmap) was projected onto the cortical surface by coloring cortical vertices according to the subcortical voxel with which they were most strongly connected. White arrows indicate principal gradient directions. Functional connectivity between each subcortical voxel and cortical vertex was computed using the preprocessed dense connectome matrix (S1200 release) from the HCP, where the functional connectivity between all pairs of grayordinates (subcortical voxels and cortical vertices) was averaged across 812 individuals and across four runs.

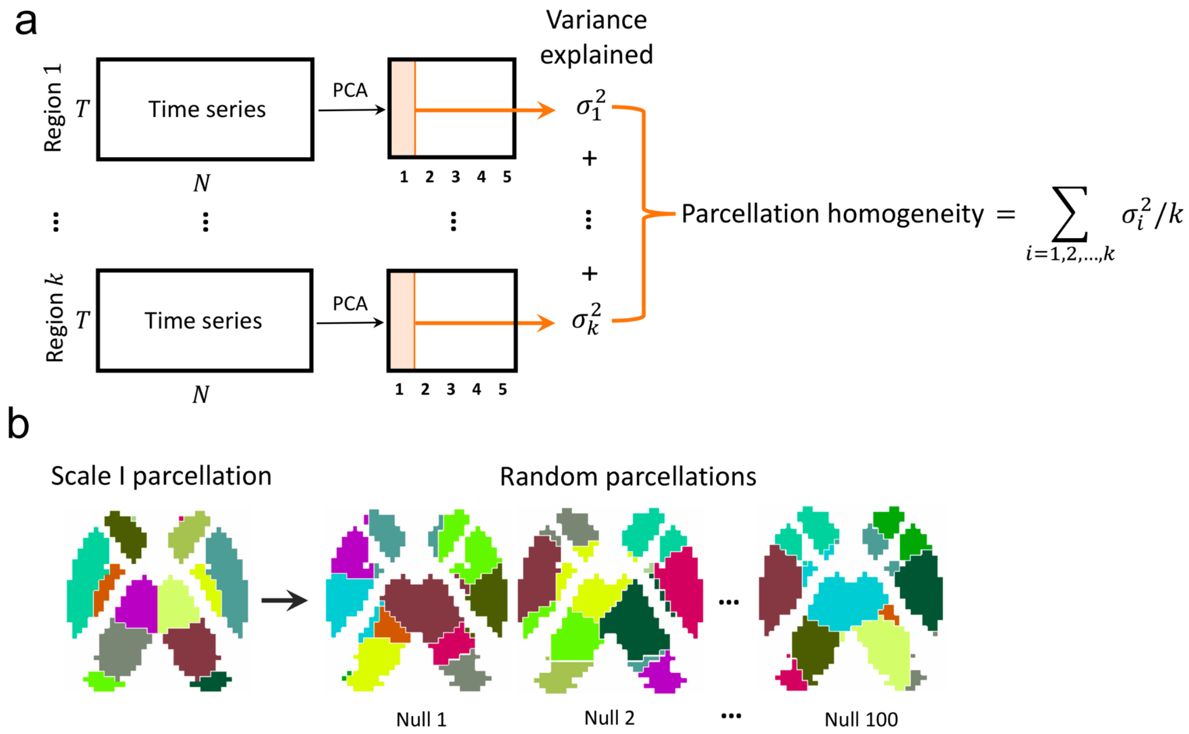

**Fig S11. Schematic of parcellation homogeneity estimation. a,** Schematic and formula representing the computation of parcellation homogeneity. Functional MRI time series for each voxel in a given region were concatenated, yielding a matrix of dimension $N \times T$, where $N$ and $T$ denote the number of voxels comprising the region and the number of time frames, respectively. Following previous work[20], principal component analysis (PCA) was applied to this matrix and the variance explained by the first principal component was estimated and referred to as regional homogeneity. The regional homogeneity was computed for each region and for each individual separately. For each individual, regional homogeneity was then averaged across all regions, yielding an overall estimate of parcellation homogeneity. **b,** Images show anatomical visualizations of the Scale I atlas (leftmost) and three instantiations of randomized versions of the atlas (right images). The random parcellations comprise an identical number of regions and comparable distribution of region sizes.

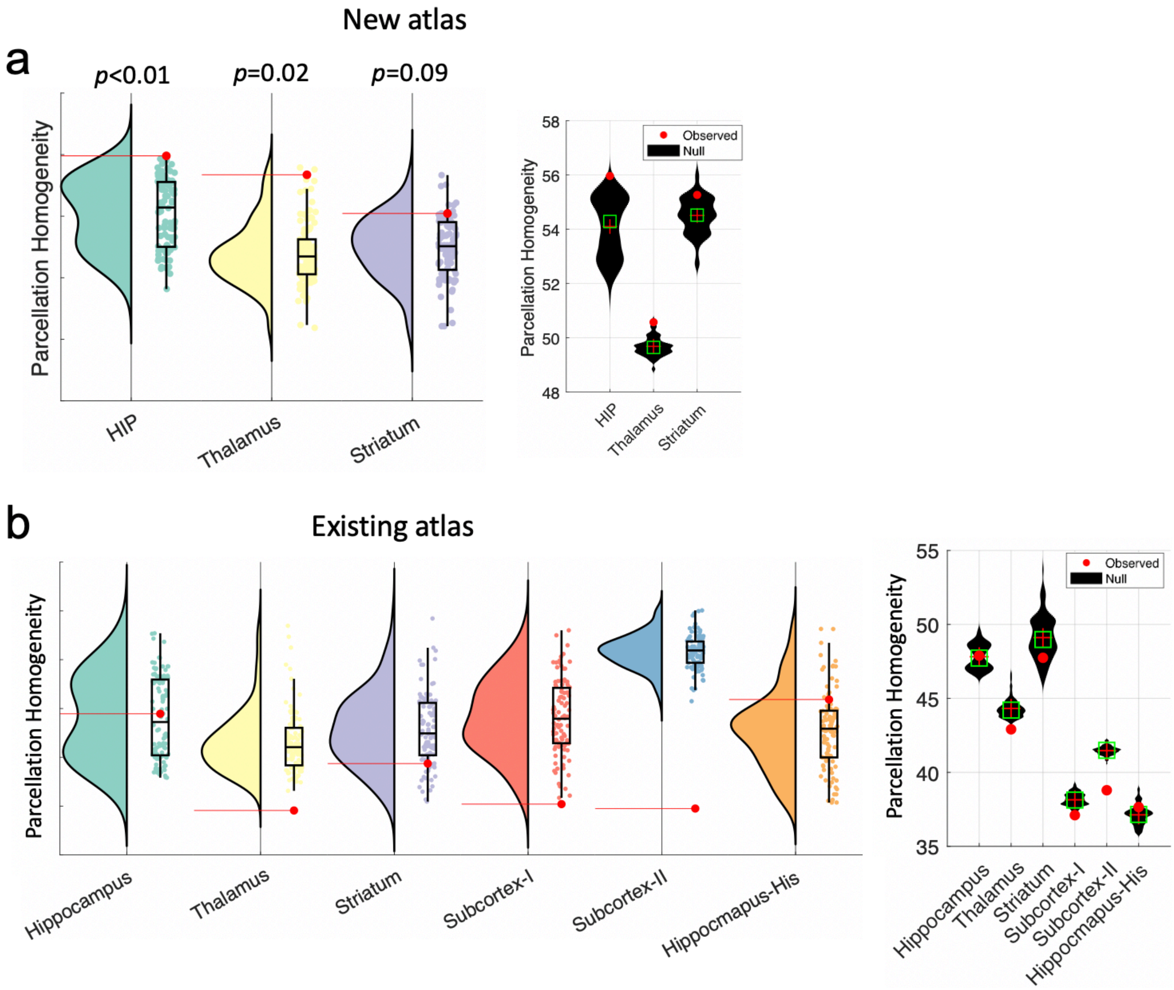

**Fig S12. Parcellation homogeneity of the new and existing subcortical parcellation atlases. a,** Parcellation homogeneity of the new atlas within the hippocampus (HIP), thalamus and striatum were benchmarked to random parcellations comprising an identical number of regions and a comparable distribution of region sizes. **b,** Parcellation homogeneity of five existing atlases was benchmarked to random parcellations comprising an identical number of regions and a comparable distribution of region sizes. Violin plots show the distribution of parcellation homogeneity measured within ensembles of 100 such random parcellations. Red dots and horizontal red lines indicate the observed parcellation homogeneity values. Bottom and top edges of the boxes indicate 25th and 75th percentiles of the distribution, respectively. The vertical axis is omitted to aid visualization. See Supplementary Table S3 for details pertaining to each atlas.

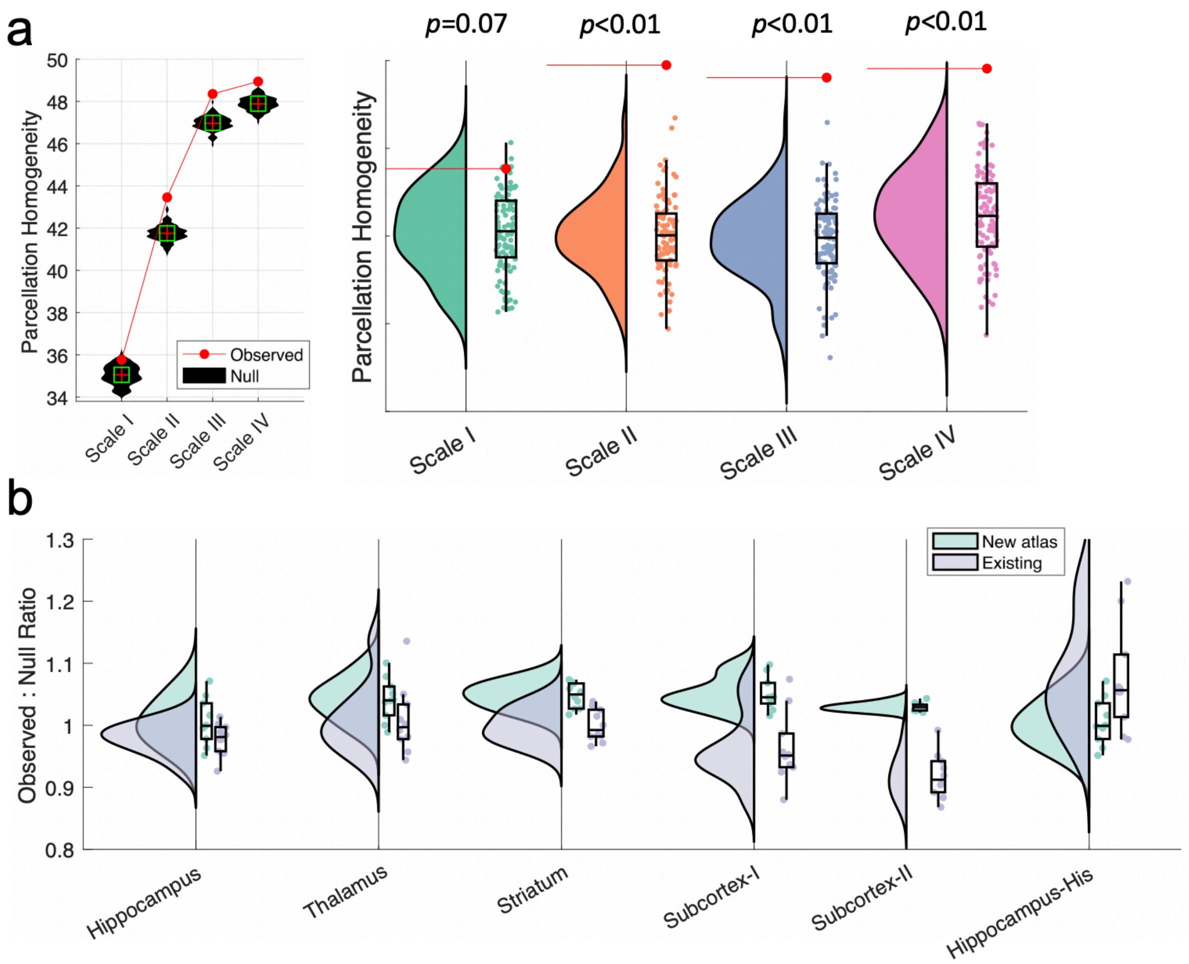

**Fig S13. Parcellation homogeneity estimated in an independent validation dataset.** Parcellation homogeneity was investigated in an independent dataset comprising 10 healthy individuals (Table S1) to exclude potential circularity confounds. **a,** Parcellation homogeneity of the new subcortex atlas was estimated and benchmarked to random parcellations of the subcortex comprising an identical number of regions and a comparable distribution of region size. The same computations performed on the primary dataset were repeated on the validation dataset (see Methods and Fig 5). Violin plots show the distribution of parcellation homogeneity measured within ensemble of 100 random parcellations. P-value shown for each scale is the proportion of random parcellations comprising the ensemble that were more homogenous than the observe parcellation. Horizontal red lines indicate the observed parcellation homogeneity values. Bottom and top edges of the boxes indicate 25th and 75th percentiles of the distribution, respectively. The vertical axis is omitted to aid visualization. The null hypothesis of parcellation homogeneity that is no better than chance could be rejected for all scales, except Scale I. **b,** Parcellation homogeneity comparison between the new atlas and existing parcellation atlases of the entire subcortex and specific subcortical nuclei (Supplementary Table S3). Parcellation homogeneity was computed separately for each individual (n=10) and normalized by the random parcellation homogeneity, yielding and observed-to-null ratio. The box plots show the distribution of this ratio across individuals for the new (turquoise) and existing (violet) parcellation atlases. Homogenous parcellations have a ratio that exceeds one.

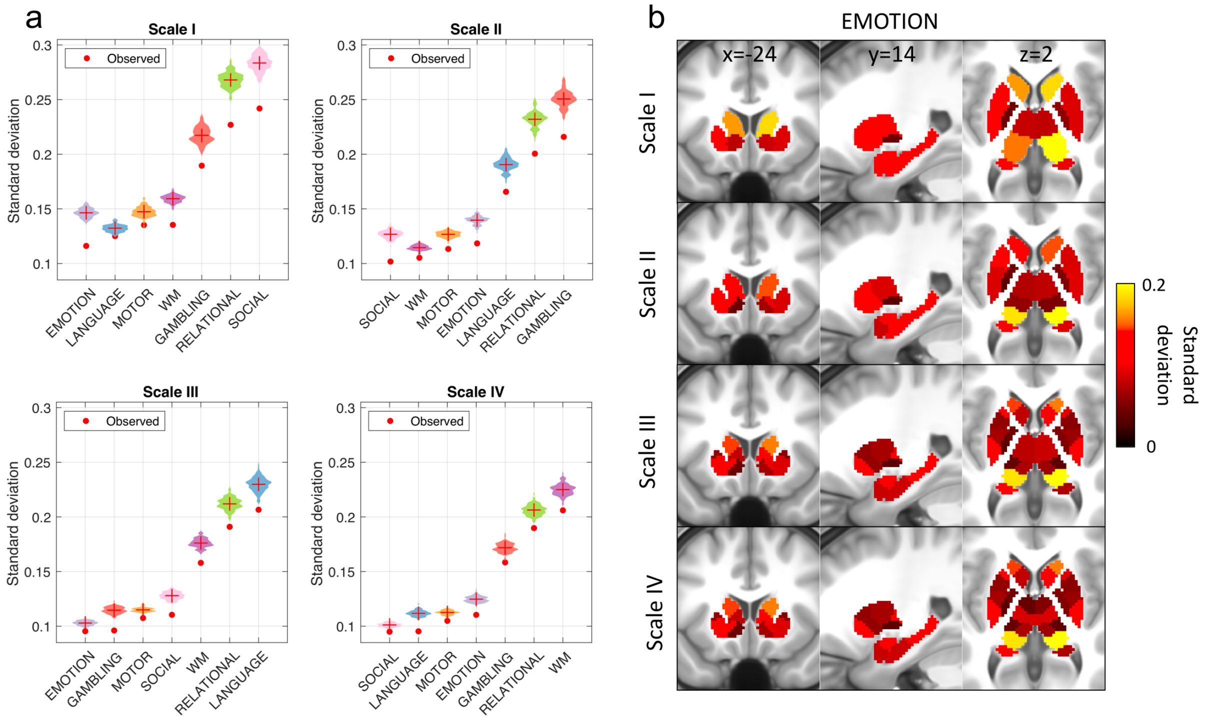

**Fig S14. Quantifying the extent to which task-evoked activity is circumscribed to regions comprising the atlas for Scale I-IV.** Task-evoked activity for seven distinct task conditions was computed by averaging over all blocked contrasts[22] and across 997 individuals (https://balsa.wustl.edu/), yielding a group- and contrast-averaged effect size map (Cohen's $d$) for each task condition. **a,** The standard deviation in task-evoked activity (i.e. effect size maps) was computed across voxels comprising each region and then averaged over all regions. This was repeated for Scale I-IV. The lower the standard deviation, the more circumscribed task-evoked activity was to particular atlas regions. The standard deviation was computed in the same way for ensembles of random subcortical parcellations (n=100). This enabled testing of the null hypothesis that the extent to which task-evoked activity was circumscribed to specific atlas regions was no greater than expected for a random parcellation. Violin plots show the distribution of standard deviation computed within ensembles of 100 random parcellations and red dots indicate the observed standard deviation values. **b,** Examples of activation homogeneity in each region at Scale I-IV in the emotional processing task.

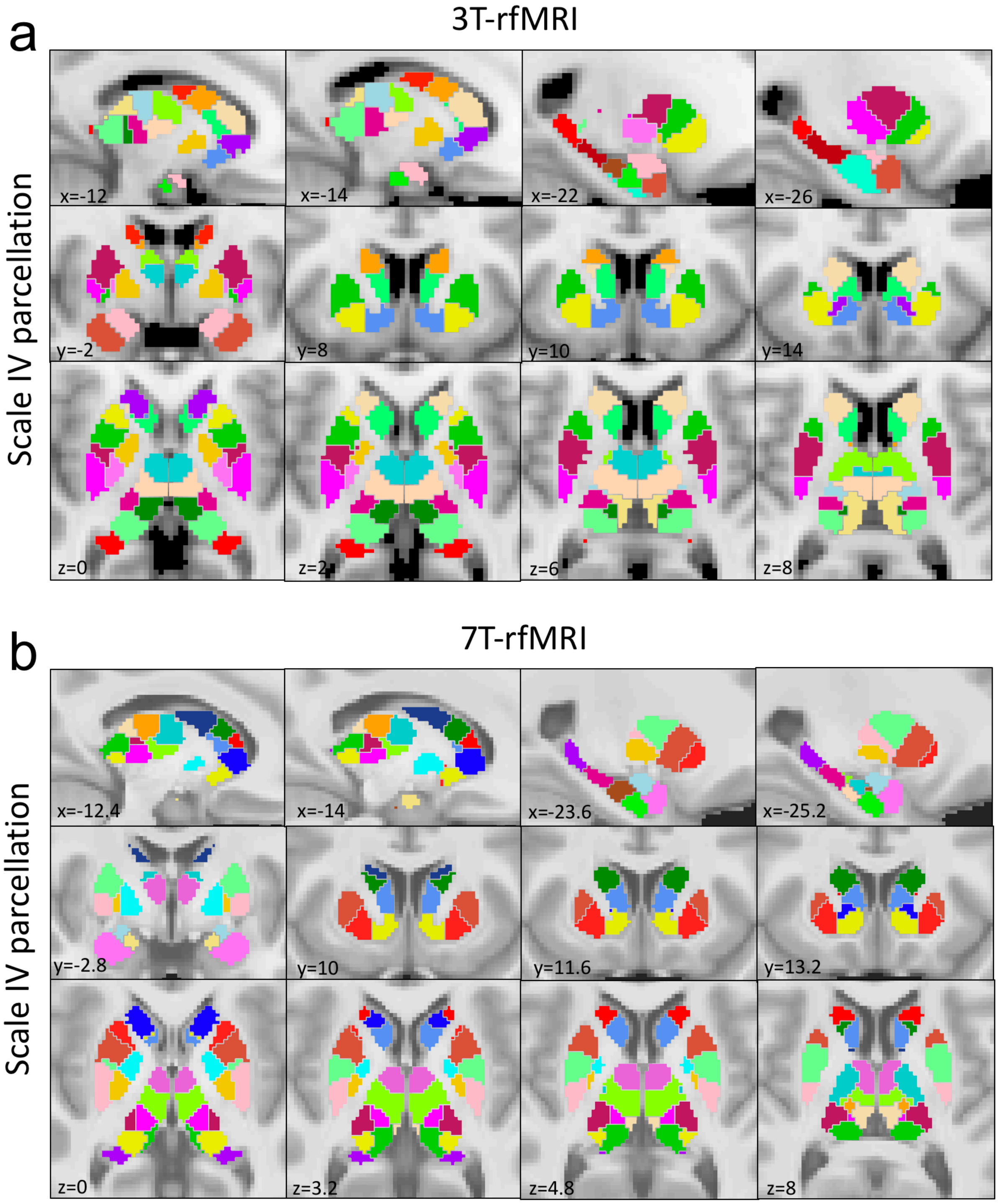

**Fig S15. 3 Tesla and 7 Tesla subcortex parcellation at Scale IV.** A total of four hierarchical scales were delineated using both 3T and 7T resting-state functional MRI (rfMRI) to parcellate the subcortex. Images show sagittal (x), coronal (y) and axial (z) slices from the group-consensus atlas at Scale IV delineated using 3T (**a**) and 7T (**b**) functional MRI. Scale IV at 3T comprises 27 bilateral regions, whereas Scale IV comprises 31 bilateral regions at 7T. The additional regions delineated at 7T include central and superior subregions of the medial amygdala, lateral and medial subregions of the hippocampal body, superior and inferior subregions of the lateral hippocampal head, and one more subregion in thalamus. All slice coordinates are indicated relative to MNI space (mm). A reference structural MRI image is used as the background. Homologous regions are colored identically.

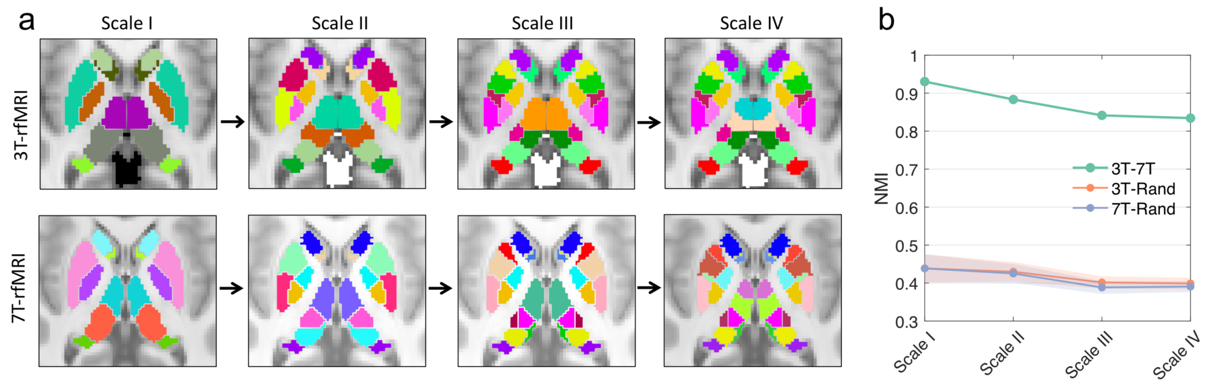

**Fig S16. Quantitative comparison between 3 Tesla and 7 Tesla atlas across parcellation scales. a,** Images show axial slices from the 3T and 7T group-consensus atlases across the four hierarchical parcellation scales (Scale I-IV, left to right). **b,** Spatial correspondence between the 3T and 7T atlas was estimated at each parcellation scale using normalized mutual information (NMI). NMI between 3T and 7T atlas (NMI, Scale I: 0.93; II: 0.88, III: 0.84, IV: 0.83, green) was benchmarked against ensembles of 100 random parcellations (3T-Rand, orange; 7T-Rand, blue) at each parcellation scale. NMI ranges from zero to one with higher value indicating greater spatial correspondence. The 3T atlas (2mm isotropic) was up-sampled to the same resolution as 7T (1.6mm isotropic) before computing NMI.

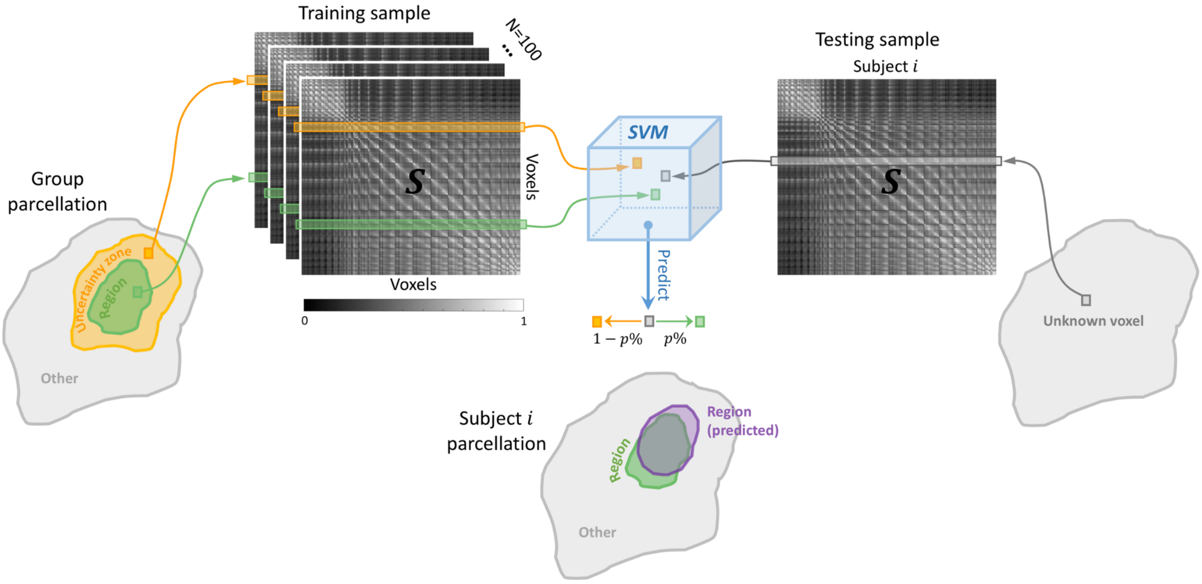

**Fig S17. Schematic of methodology for parcellation personalization.** A binary support vector machine (SVM) classifier (blue box) was trained to classify whether a voxel resided in the region proper (green) or its uncertainty zone (orange). The SVM feature space was defined over the similarity matrix ($S$), where each cell stored the similarity in functional connectivity between a pair of subcortical voxels. The feature for a particular voxel was selected as the relevant row in this matrix, highlighted by green and orange colored strips. One hundred randomly selected individuals were used to train a separate SVM classifier for each region. The uncertainty zone for each region was delineated by dilating the region's mask (see Methods). For a new individual (Subject $i$), the trained classifier was used to predict the posterior probability ($p\%$) of an unknown voxel (gray) belonging to the region as opposed to belonging to the region's uncertainty zone ($1 - p\%$). This was repeated for each region comprising the Scale IV atlas. The uncertainty zones of two distinct regions could potentially overlap, meaning that voxels could be assigned a probability of belonging to multiple regions. Such voxels were assigned to the region associated with the highest posterior probability, yielding a personalized parcellation atlas for Subject $i$ (purple).

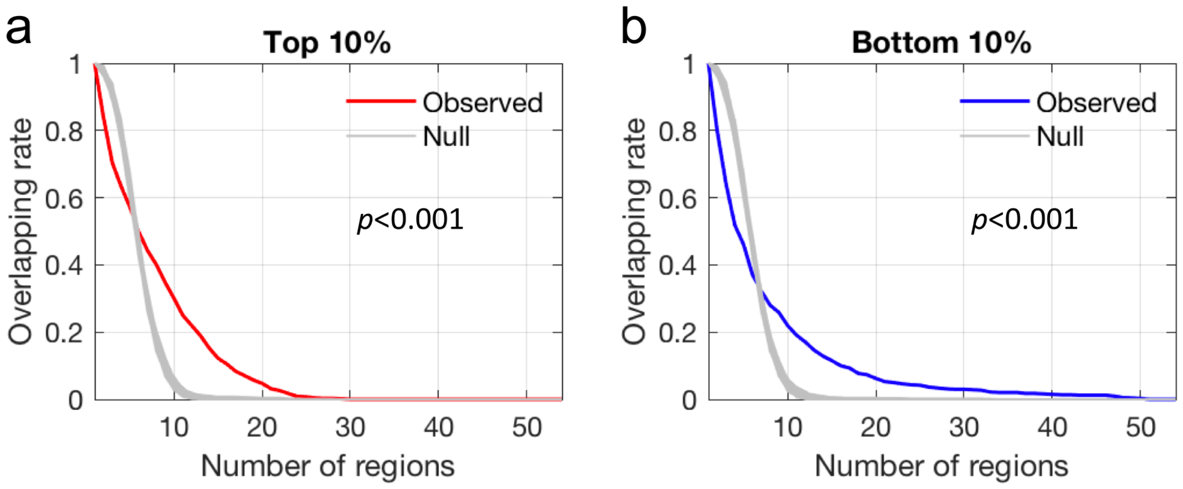

**Fig S18. Within-subject variation in Dice coefficient across subcortical regions.** The Dice coefficient for each Scale IV subcortical region in each individual was encoded in a Dice-matrix of dimension 921 × 54, where 921 is the total number individuals and 54 is the number of subcortical regions. Rows in this matrix were then re-ordered from the highest to lowest Dice coefficient. This re-ordering was performed separately for each column. Individuals ranked among the top 10% (high Dice) or bottom 10% (low Dice) for at least one column (region) were selected, yielding 696 and 739 unique individuals in each group, respectively. Some individuals only presented once, whereas some individuals repeatedly presented in more than one region. The proportions of individuals (overlapping rate) that repeatedly presented in at least one or more subcortical regions were computed. The curves show the overlapping rate as a functional of the number of regions in the high Dice (**a,** red curve) and low Dice (**b,** blue curve) group separately. The area under the curve of the overlapping rate was computed. Permutation testing (n=1000) was used to test whether the overlapping rate was larger than expected due to chance. Note that each row in the Dice-matrix was shuffled as a whole in each permutation to control the relationship across regions. The overlapping rate was recomputed for each permuted sample and plotted as a functional of number of regions (Null, gray curves). The p-values shown are given by the proportion of null curves that had larger area under the curve than the observed curve.

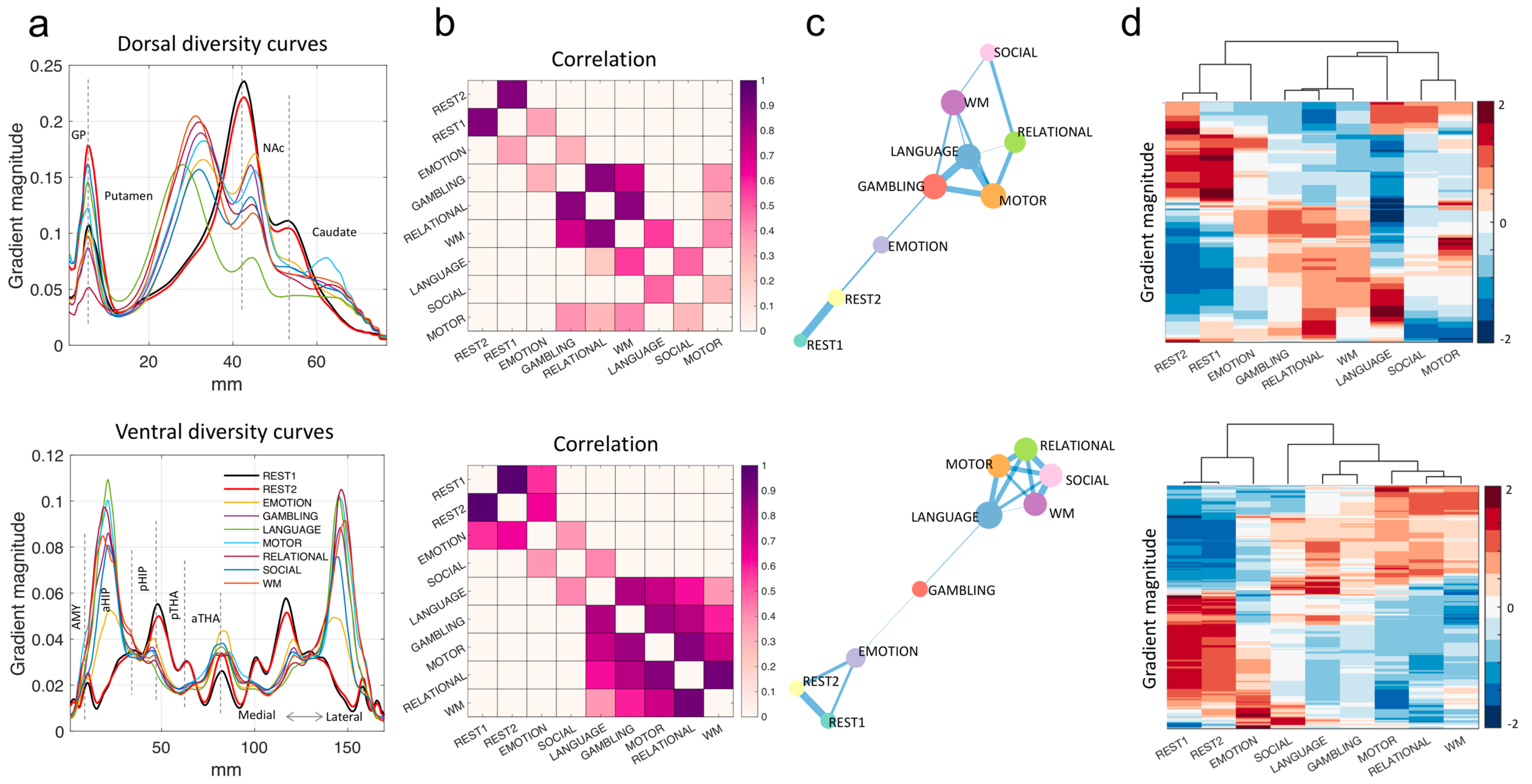

**Fig S19. Variation in subcortical connectivity gradients between rest and task-evoked conditions. a,** Dorsal and ventral diversity curves are shown for each rest and task condition separately. **b,** Similarity in the diversity curves between each pair of conditions was estimated using the Pearson correlation coefficient. Permutation testing (n=10,000) was used to determine the significance (p-value) of each correlation coefficient using a corrected significance threshold of $p < 0.05/36=0.0014$ (Bonferroni correction). Correlation coefficients surviving Bonferroni correction are colored in the matrix shown and also visualized as a graph. **c,** In the graph, each node represents one of the nine conditions and the edges between them indicate correlations. Node size is scaled by nodal degree, whereas edge thickness varies according to the correlation coefficient magnitude. **d,** Matrix representation of diversity curves for rest and task conditions. Matrix rows correspond to specific points along the diversity curves. Matrix columns correspond to rest and task conditions. Rows and columns are reordered to accentuate similarity between diversity curves. Average-linkage clustering was used to determine the reordering and the cluster tree shown.

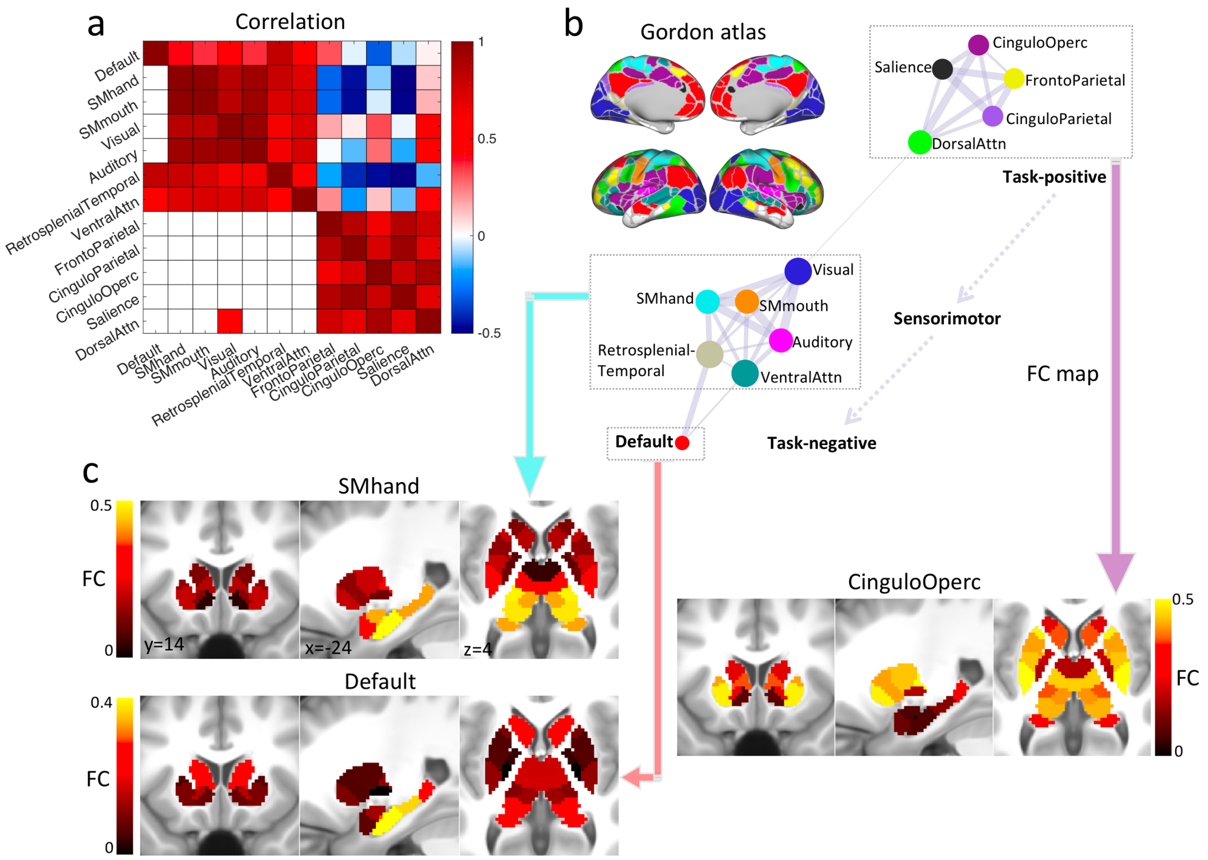

**Fig S20. Linking canonical cortical networks based on their functional connectivity with subcortical regions.** Functional connectivity (FC) was computed between each Scale IV subcortical region and each cortical parcel delineated in an existing cortical atlas[20], yielding a connectivity matrix of dimension 54 x 333 for each individual. The connectivity matrices were then averaged across all individuals (REST1, n=1080) to enable a group-consensus representation of cortical-subcortical connectivity. Each column in this matrix thus represents a subcortical connectivity map for a given cortical parcel. Subcortical connectivity maps were averaged across cortical parcels that belong to the same canonical network[20], yielding 12 FC maps that represent how each cortical network connects to the subcortex. The 12 networks are as follows: visual, dorsal somatomotor (SMmouth), ventral somatomotor (SMhand), auditory, default, frontoparietal, dorsal attention, ventral attention, cingulo-opercular (cinguloOperc), salience, cingulo-parietal and retrosplenial-temporal network. **a,** Similarity in subcortical FC maps between each pair of cortical networks was estimated using the Pearson correlation coefficient, yielding a 12×12 correlation matrix. The rows and columns of this matrix were reordered to accentuate modular structure, which was determined by the Louvain community detection algorithm[21]. The 12 × 12 correlation matrix was thresholded using a statistical approach based on null hypothesis testing. A p-value was computed for each correlation coefficient as follows. Permutation testing (n=1000) was used to determine the significance (p-value) of each correlation coefficient using a corrected significance threshold of $p<0.05/66=0.008$ (Bonferroni correction). Correlation coefficients that survived correction are shown in the lower triangle of the correlation matrix and visualized as a graph. **b,** In this graph, each node represents a canonical cortical network. Edges are drawn between pairs of networks that share common patterns of functional connectivity with subcortical regions. Node size is scaled by nodal degree, whereas edge thickness varies according to the correlation coefficient. Nodes were parsed into three groups (gray blocks) and positioned on a task-positive to task-negative organizational axis. **c,** Representative subcortical FC maps for each group of canonical networks. It can be seen that the cingulo-opercular network (task-positive) is preferentially connected to thalamus and striatum, especially the dorsal striatum. In contrast, sensorimotor networks are predominantly connected to the hippocampus, amygdala and posterior thalamus. The default mode network (task-negative) is preferentially connected to the hippocampus but also show moderate connectivity with posterior thalamus and caudate.

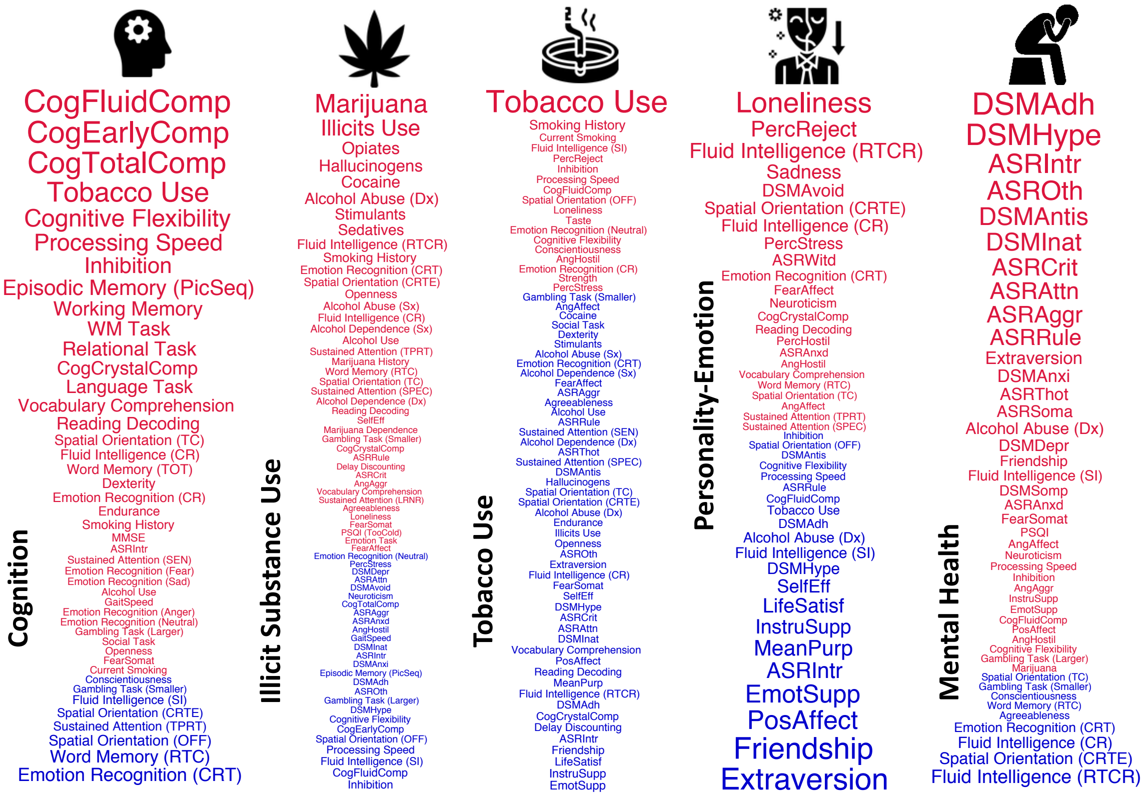

**Fig S21. Behavioral dimensions.** Five orthogonal behavioral dimensions characterizing: i) cognition, ii) illicit substance use, iii) tobacco use, iv) personality and emotion traits, as well as v) mental health were derived from a total of 109 behavioral items using independent component analysis (see Methods). The top weighted (30-40%) behavioral items in each dimension are visualized as word clouds. The font size of each item is scaled according to its absolute weight in the ICA de-mixing matrix, and the font color denotes weight polarity (red-positive, blue-negative). Behavioral icons sourced from: https://www.flaticon.com/

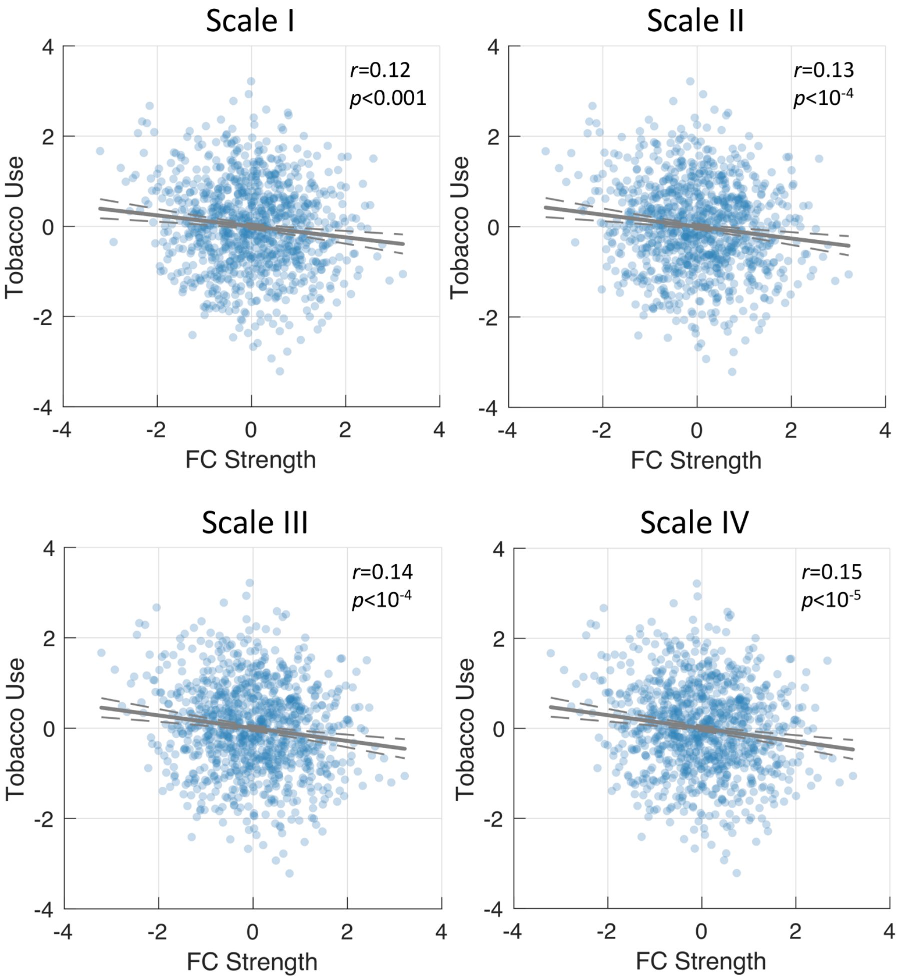

**Fig S22. Association between subcortical functional connectivity and the behavioral dimension characterizing tobacco use reproduced using functional MRI from a second session.** Scatter plots and lines of best fit show the association between individual variation in tobacco use dimension and functional connectivity (FC) strength within the thalamo-striato-hippocampal network measured using functional MRI acquired in the REST1 session. The significant association was discovered using functional MRI acquired in REST2 (see Fig 8). Each blue dot in the scatter plot represents one individual. Solid gray lines indicate lines of best fit. Dashed lines indicate 95% confidence intervals. The association between subcortical functional connectivity and tobacco use was reproducible across independent functional MRI sessions (REST1 and REST2) and across parcellation scales (Scale I-IV).

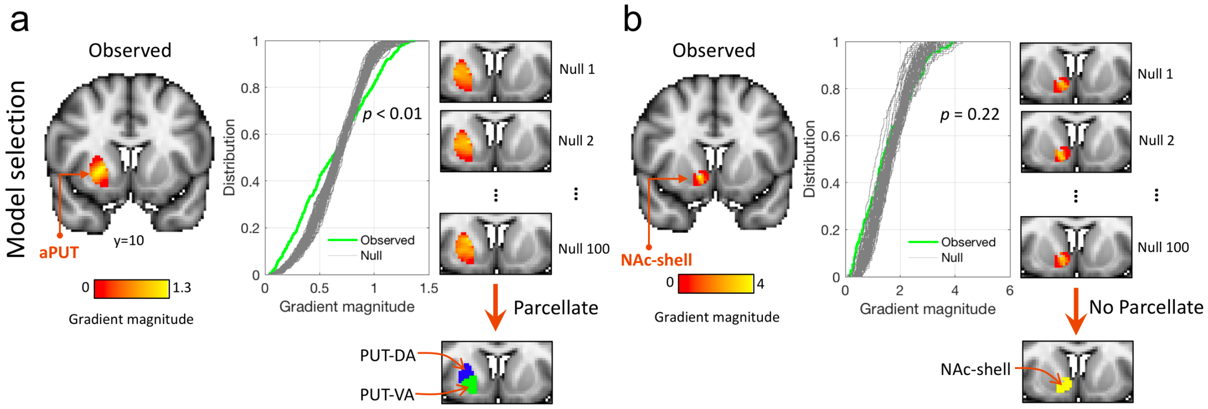

**Fig S23. Model selection with Kolmogorov-Smirnov (KS) test. a,** Gradient magnitude image for anterior putamen (aPUT) is shown as a coronal slice (left). The yellow colored strip within the aPUT indicates a gradient magnitude peak, suggesting the location of a putative boundary. The KS test suggests that the gradient magnitude distribution across the voxels in this region (green curve) is significantly longer tailed than the null data (gray curves, $p<0.01$). Therefore, a boundary was delineated that resulted in parcellation of the anterior putamen into one dorsal (PUT-DA, blue) and one ventral (PUT-VA, green) component. **b,** KS test for the shell of the nucleus accumbens (NAc-shell). Although a strong gradient is evident, the null hypothesis could not be rejected with the KS test ($p=0.22$), and thus subdivision of NAc-shell was not warranted.

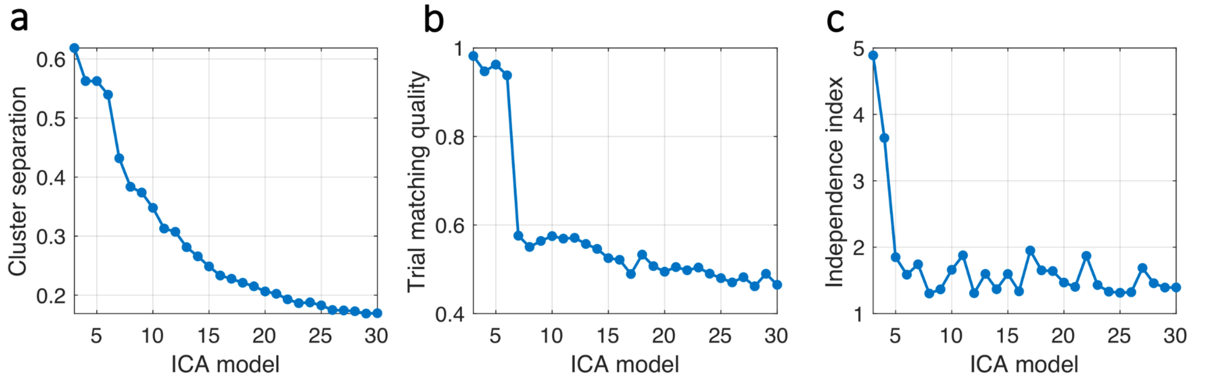

**Fig S24. Determining the optimal number of independent components.** Independent component analysis (ICA) was used to decompose a total of 109 behavioral measures into a set of latent behavioral dimensions. ICA performance for candidate ICA models (i.e. total number of components) ranging from 3 to 30 was evaluated based on cluster separation (**a**), trial matching quality (**b**) and independence index (**c**). Each measure was computed as a function of the total number of components and selection of the optimal model was guided by identifying abrupt changes in these measures as a function of the ICA model.

### Supplementary Tables

**Table S1. Basic demographic characteristics and acquisition details**

| Dataset | Modality | Magnetic field | Sample size | Age (mean $\pm$ std yrs) | Sex (M/F) |
|---|---|---|---|---|---|
| 3T-rfMRI (REST1) | Resting-state | 3 Tesla | 1080 | $28.8 \pm 3.7$ | 495/585 |
| 3T-rfMRI (REST2) | Resting-state | 3 Tesla | 1021 | $28.7 \pm 3.7$ | 471/550 |
| 7T-rfMRI | Resting-state | 7 Tesla | 183 | $29.4 \pm 3.3$ | 72/111 |
| tfMRI | Task-evoked | 3 Tesla | 725 | $28.7 \pm 3.7$ | 360/365 |
| Independent validation dataset* | Resting-state | 3 Tesla | 10 | $26.0 \pm 2.1$ | 6/4 |

*Image acquisition and data pre-processing are described in **S3**. **Independent validation dataset** section.

**Table S2. Nomenclature for 3T parcellation hierarchy**

| Name | Abbreviation | Name | Abbreviation |
|---|---|---|---|
| **Hippocampus** | HIP | **Amygdala** | AMY |
| Anterior hippocampus | aHIP | Lateral amygdala | lAMY |
| Hippocampus head, medial division | HIP-head-m | Medial amygdala | mAMY |
| Subdivision1 | HIP-head-m1 | **Caudate nucleus** | CAU |
| Subdivision2 | HIP-head-m2 | Anterior caudate | aCAU |
| Hippocampus head, lateral division | HIP-head-l | Dorsoanterior caudate | CAU-DA |
| Posterior hippocampus | pHIP | Ventroanterior caudate | CAU-VA |
| Hippocampus body | HIP-body | Posterior caudate | pCAU |
| Hippocampus tail | HIP-tail | Caudate tail | CAU-tail |
| **Thalamus** | THA | Caudate body | CAU-body |
| Anterior thalamus | aTHA | **Nucleus accumbens** | NAc |
| Dorsoanterior thalamus | THA-DA | Nucleus accumbens, shell | NAc-shell |
| Lateral dorsoanterior thalamus | THA-DAl | Nucleus accumbens, core | NAc-core |
| Medial dorsoanterior thalamus | THA-DAm | **Putamen** | PUT |
| Ventroanterior thalamus | THA-VA | Anterior putamen | aPUT |
| Superior ventroanterior thalamus | THA-Vas | Dorsoanterior putamen | PUT-DA |
| Inferior ventroanterior thalamus | THA-VAi | Ventroanterior putamen | PUT-VA |
| Anterior division | THA-VAia | Posterior putamen | pPUT |
| Posterior division | THA-VAip | Dorsoposterior putamen | PUT-DP |
| Posterior thalamus | pTHA | Ventroposterior putamen | PUT-VP |
| Dorsoposterior thalamus | THA-DP | **Globus pallidus** | GP |
| Ventroposterior thalamus | THA-VP | Anterior globus pallidus | aGP |
| Medial ventroposterior thalamus | THA-VPm | Posterior globus pallidus | pGP |
| Lateral ventroposterior thalamus | THA-VPl | | |

**Table S3. Comparison parcellation atlases of the human subcortex and specific subcortical nuclei**

| Atlas | Study | Modality | Metric | Number of regions |
|---|---|---|---|---|
| Hippocampus | Plachti et al. 2019[23] | Resting-state fMRI | Functional connectivity | 5 |
| Thalamus | Behrens et al. 2003[24] | Diffusion MRI | Tractography | 7 |
| Striatum | Janssen et al. 2015[25] | Resting-state fMRI | Functional connectivity | 6 |
| Subcortex-I | Ji et al. 2019[26] | Resting-state fMRI | Functional connectivity | 12 |
| Subcortex-II | Fan et al. 2016[27] | Diffusion MRI | Tractography | 36 |
| Hippocampus - His | Amunts et al 2005[11]; Eickhoff et al 2005[12] | Histology | Cytoarchitecture | 3 |

**Table S4. List of 109 selected behavioral items**

| Category | Formal Name | Intuitive Name | Psychological Test |
|---|---|---|---|
| | MMSE_Score | MMSE | Mini Mental Status Exam |
| | PSQI_Score | PSQI | Pittsburgh Sleep Questionnaire |
| | PSQI_TooCold | PSQI (TooCold) | Pittsburgh Sleep Questionnaire |
| **Alertness** | PSQI_TooHot | PSQI (TooHot) | Pittsburgh Sleep Questionnaire |
| | PSQI_BadDream | PSQI (BadDream) | Pittsburgh Sleep Questionnaire |
| | PSQI_Pain | PSQI (Pain) | Pittsburgh Sleep Questionnaire |
| | PicSeq_Unadj | Episodic Memory (PicSeq) | NIH Toolbox Picture Sequence Memory Test |
| | CardSort_Unadj | Cognitive Flexibility | NIH Toolbox Dimensional Change Card Sort |
| | Flanker_Unadj | Inhibition | NIH Toolbox Flanker Inhibitory Control and Attention |
| | PMAT24_A_CR | Fluid Intelligence (CR) | Penn Progressive Matrices |
| | PMAT24_A_SI | Fluid Intelligence (SI) | Penn Progressive Matrices |
| | PMAT24_A_RTCR | Fluid Intelligence (RTCR) | Penn Progressive Matrices |
| | ReadEng_Unadj | Reading Decoding | NIH Toolbox Oral Recognition Test |
| | PicVocab_Unadj | Vocabulary Comprehension | NIH Toolbox Picture Vocabulary Test |
| | ProcSpeed_Unadj | Processing Speed | NIH Toolbox Pattern Comparison Processing Speed Test |
| | DDisc_AUC_200 | Delay Discounting | Delay Discounting |
| | VSPLOT_TC | Spatial Orientation (TC) | Variable Short Penn Line Orientation |
| | VSPLOT_CRTE | Spatial Orientation (CRTE) | Variable Short Penn Line Orientation |
| | VSPLOT_OFF | Spatial Orientation (OFF) | Variable Short Penn Line Orientation |
| | SCPT_TPRT | Sustained Attention (TPRT) | Short Penn Continuous Performance Test |
| | SCPT_SEN | Sustained Attention (SEN) | Short Penn Continuous Performance Test |
| **Cognition** | SCPT_SPEC | Sustained Attention (SPEC) | Short Penn Continuous Performance Test |
| | SCPT_LRNR | Sustained Attention (LRNR) | Short Penn Continuous Performance Test |
| | IWRD_TOT | Word Memory (TOT) | Penn Word Memory Test |
| | IWRD_RTC | Word Memory (RTC) | Penn Word Memory Test |
| | ListSort_Unadj | Working Memory | NIH Toolbox List Sorting Working Memory Test |
| | CogFluidComp_Unadj | CogFluidComp | NIH Toolbox Cognition Fluid Composite |
| | CogEarlyComp_Unadj | CogEarlyComp | NIH Toolbox Cognition Early Childhood Composite |
| | CogTotalComp_Unadj | CogTotalComp | NIH Toolbox Cognition Total Composite Score |
| | CogCrystalComp_Unadj | CogCrystalComp | NIH Toolbox Cognition Crystallized Composite |
| | ER40_CR | Emotion Recognition (CR) | Penn Emotion Recognition Test |
| | ER40_CRT | Emotion Recognition (CRT) | Penn Emotion Recognition Test |
| | ER40ANG | Emotion Recognition (Anger) | Penn Emotion Recognition Test |
| | ER40FEAR | Emotion Recognition (Fear) | Penn Emotion Recognition Test |
| | ER40HAP | Emotion Recognition (Happy) | Penn Emotion Recognition Test |
| | ER40NOE | Emotion Recognition (Neutral) | Penn Emotion Recognition Test |
| | ER40SAD | Emotion Recognition (Sad) | Penn Emotion Recognition Test |
| | AngAffect_Unadj | AngAffect | NIH Toolbox Anger-Affect Survey |
| | AngHostil_Unadj | AngHostil | NIH Toolbox Anger-Hostility Survey |

| | | | |
|---|---|---|---|
| **Emotion** | AngAggr_Unadj | AngAggr | NIH Toolbox Anger-Physical Aggression Survey |
| | FearAffect_Unadj | FearAffect | NIH Toolbox Fear-Affect Survey |
| | FearSomat_Unadj | FearSomat | NIH Toolbox Fear-Somatic Arousal Survey |
| | Sadness_Unadj | Sadness | NIH Toolbox Sadness Survey |
| | LifeSatisf_Unadj | LifeSatisf | NIH Toolbox General Life Satisfaction Survey |
| | MeanPurp_Unadj | MeanPurp | NIH Toolbox Meaning and Purpose Survey |
| | PosAffect_Unadj | PosAffect | NIH Toolbox Positive Affect Survey |
| | Friendship_Unadj | Friendship | NIH Toolbox Friendship Survey |
| | Loneliness_Unadj | Loneliness | NIH Toolbox Loneliness Survey |
| | PercHostil_Unadj | PercHostil | NIH Toolbox Perceived Hostility Survey |
| | PercReject_Unadj | PercReject | NIH Toolbox Perceived Rejection Survey |
| | EmotSupp_Unadj | EmotSupp | NIH Toolbox Emotional Support Survey |
| | InstruSupp_Unadj | InstruSupp | NIH Toolbox Instrumental Support Survey |
| | PercStress_Unadj | PercStress | NIH Toolbox Perceived Stress Survey |
| | SelfEff_Unadj | SelfEff | NIH Toolbox Self-Efficacy Survey |
| **Motor** | Endurance_Unadj | Endurance | NIH Toolbox 2-minute Walk Endurance Test |
| | GaitSpeed_Comp | GaitSpeed | NIH Toolbox 4-Meter Walk Gait Speed Test |
| | Dexterity_Unadj | Dexterity | NIH Toolbox 9-hole Pegboard Dexterity Test |
| | Strength_Unadj | Strength | NIH Toolbox Grip Strength Test |
| **Personality** | NEOFAC_A | Agreeableness | NEO-FFI |
| | NEOFAC_O | Openness | NEO-FFI |
| | NEOFAC_C | Conscientiousness | NEO-FFI |
| | NEOFAC_N | Neuroticism | NEO-FFI |
| | NEOFAC_E | Extraversion | NEO-FFI |
| **Sensory** | Odor_Unadj | Odor | NIH Toolbox Odor Identification Scale |
| | PainInterf_Tscore | PainInterf | NIH Toolbox Pain Interference Survey |
| | Taste_Unadj | Taste | NIH Toolbox Regional Taste Intensity |
| | Mars_Final | Visual Contrast Sensitivity | Mars Contrast Sensitivity Score |
| **Psychiatric and Life Function** | DSM_Depr_Raw | DSMDepr | ASR-DSM |
| | DSM_Anxi_Raw | DSMAnxi | ASR-DSM |
| | DSM_Somp_Raw | DSMSomp | ASR-DSM |
| | DSM_Avoid_Raw | DSMAvoid | ASR-DSM |
| | DSM_Adh_Raw | DSMAdh | ASR-DSM |
| | DSM_Inat_Raw | DSMInat | ASR-DSM |
| | DSM_Hype_Raw | DSMHype | ASR-DSM |
| | DSM_Antis_Raw | DSMAntis | ASR-DSM |
| | ASR_Anxd_Raw | ASRAnxd | ASR-DSM |
| | ASR_Witd_Raw | ASRWitd | ASR-DSM |
| | ASR_Soma_Raw | ASRSoma | ASR-DSM |
| | ASR_Thot_Raw | ASRThot | ASR-DSM |
| | ASR_Attn_Raw | ASRAttn | ASR-DSM |
| | ASR_Aggr_Raw | ASRAggr | ASR-DSM |

| | | | |
|---|---|---|---|
| | ASR_Rule_Raw | ASRRule | ASR-DSM |
| | ASR_Intr_Raw | ASRIntr | ASR-DSM |
| | ASR_Oth_Raw | ASROth | ASR-DSM |
| | ASR_Crit_Raw | ASRCrit | ASR-DSM |
| **Substance Use** | Num_Days_Drank_7days | Alcohol Use | Alcohol Use 7-Day Retrospective |
| | SSAGA_Alc_D4_Dp_Sx | Alcohol Dependence (Sx) | Alcohol Use and Dependence |
| | SSAGA_Alc_D4_Ab_Dx | Alcohol Abuse (Dx) | Alcohol Use and Dependence |
| | SSAGA_Alc_D4_Ab_Sx | Alcohol Abuse (Sx) | Alcohol Use and Dependence |
| | SSAGA_Alc_D4_Dp_Dx | Alcohol Dependence (Dx) | Alcohol Use and Dependence |
| | Num_Days_Used_Any_Tobacco_7days | Tobacco Use | Tobacco Use 7-Day Retrospective |
| | SSAGA_TB_Smoking_History | Smoking History | Tobacco Use and Dependence |
| | SSAGA_TB_Still_Smoking | Current Smoking | Tobacco Use and Dependence |
| | SSAGA_Times_Used_Illicits | Illicits Use | Illicit Drug Use |
| | SSAGA_Times_Used_Cocaine | Cocaine | Illicit Drug Use |
| | SSAGA_Times_Used_Hallucinogens | Hallucinogens | Illicit Drug Use |
| | SSAGA_Times_Used_Opiates | Opiates | Illicit Drug Use |
| | SSAGA_Times_Used_Sedatives | Sedatives | Illicit Drug Use |
| | SSAGA_Times_Used_Stimulants | Stimulants | Illicit Drug Use |
| | SSAGA_Mj_Use | Marijuana History | Marijuana Use and Dependence |
| | SSAGA_Mj_Ab_Dep | Marijuana Dependence | Marijuana Use and Dependence |
| | SSAGA_Mj_Times_Used | Marijuana | Marijuana Use and Dependence |
| **In-Scanner Task** | Emotion_Task_Acc | Emotion Task | In-Scanner Task Performance |
| | Gambling_Task_Perc_Larger | Gambling Task (Larger) | In-Scanner Task Performance |
| | Gambling_Task_Perc_Smaller | Gambling Task (Smaller) | In-Scanner Task Performance |
| | Language_Task_Acc | Language Task | In-Scanner Task Performance |
| | Relational_Task_Acc | Relational Task | In-Scanner Task Performance |
| | Social_Task_Perc_TOM | Social Task | In-Scanner Task Performance |
| | WM_Task_Acc | WM Task | In-Scanner Task Performance |

**Table S5. NIFTI and CIFTI file names for 3T and 7T atlas**

| Magnetic strength | Scale | Number of regions | Spatial resolution (mm)[a] | File name[b] |
|---|---|---|---|---|
| 3 Tesla | I | 16 | 2×2×2 | Tian_Subcortex_S1_3T.nii<br>Tian_Subcortex_S1_3T_dscalar.nii |
| | II | 32 | | Tian_Subcortex_S2_3T.nii<br>Tian_Subcortex_S2_3T_dscalar.nii |
| | III | 50 | | Tian_Subcortex_S3_3T.nii<br>Tian_Subcortex_S3_3T_dscalar.nii |
| | IV | 54 | | Tian_Subcortex_S4_3T.nii<br>Tian_Subcortex_S4_3T_dscalar.nii |
| 7 Tesla | I | 16 | 1.6×1.6×1.6 | Tian_Subcortex_S1_7T.nii<br>Tian_Subcortex_S1_7T_dscalar.nii |
| | II | 34 | | Tian_Subcortex_S2_7T.nii<br>Tian_Subcortex_S2_7T_dscalar.nii |
| | III | 54 | | Tian_Subcortex_S3_7T.nii<br>Tian_Subcortex_S3_7T_dscalar.nii |
| | IV | 62 | | Tian_Subcortex_S4_7T.nii<br>Tian_Subcortex_S4_7T_dscalar.nii |

a, Atlas is in MNI standard space (MNI ICBM 152 nonlinear 6$^{th}$ generation)
b, NIFTI: *.nii; CIFTI: *dscalar.nii. Atlas is openly available at: https://github.com/yetianmed/subcortex

#### S1. Connectivity gradients and subcortical geometry

A clear peak in the diversity curve separates the hippocampus and thalamus (Fig 2d). However, the transition between the hippocampus and thalamus is quite narrow and represents a local constriction in the geometry of the subcortex. Because our resampling preserves the geometry of the empirical data, a relatively abrupt transition is also present between the thalamus and hippocampus in the null data. In other words, the functional separation between the hippocampus and thalamus can be largely explained by the intrinsic geometry of the subcortex. This does not imply that the hippocampus and thalamus are not functionally separated but simply suggests that the functional separation is consistent with the underlying geometry of the two structures and does not require an additional disruption in the functional connectivity gradients. Conversely, there are occasional peaks in the null diversity curves that are not present in the real data (e.g. the broad peak within the caudate, Fig 2c). This presumably arises when the local functional connectivity gradient is smoother (more homogenous) in the real data across a geometric feature than predicted by the null data.

#### S2. Multiscale organization and model selection

Multiscale (hierarchical) organization is not necessarily a *fait accompli* of the recursive application of model selection. Indeed, we explicitly tested for the possibility of an absence of multiscale structure for each region comprising Scale I, but found that the null hypothesis could be rejected for all regions, suggesting multiscale organizational structure. If we were unable to reject the null hypothesis, this would have instead suggested an absence of any multiscale structure. Furthermore, the number of scales differed substantially between subcortical nuclei. For example, the thalamus, hippocampus and caudate each comprised four levels, suggesting a deeper and more complex multiscale structure relative to the amygdala and globus pallidus, which only comprised two levels. This is consistent with the greater organizational complexity of the thalamic nuclei, from molecular[29] and cellular[10], to connectivity[24, 30, 31] and function[32, 33]. In contrast, the globus pallidus is primarily involved in the regulation of movement control[34] and the amygdala plays an essential role in emotional process[35].

#### S3. Independent validation dataset

An independent fMRI dataset was acquired in Australia to test the reproducibility of the parcellation homogeneity results. Ten healthy adults were recruited and confirmed not to have any current or history of any neurological or psychiatric disorders at the time of MR scanning (mean age 26$\pm$2.1yrs, 6 males). Imaging was performed in all participants using a Siemens Prisma 3T MRI scanner. Structural images of brain anatomy were acquired using an optimized Magnetization-prepared Rapid Acquisition Gradient Echo (MPRAGE) T1-weighted sequence with 176 sagittal slices of 1mm thickness; field of view (FOV)=240$\times$256 mm$^2$; flip angle = 9°; repetition time (TR)=1900ms; echo time (TE)=2.98ms; voxel size=1.0$\times$1.0$\times$1.0 mm$^3$. Resting-state fMRI was acquired using a T2*-weighted multiband gradient-echo EPI sequence of approximately 12 mins, resulting in 880 volumes (TR=810ms, TE=30ms, voxel size=2.0$\times$2.0$\times$2.0 mm$^3$).

Resting-state fMRI data were preprocessed using fMRIPrep-1.5.9 pipeline[28]. In brief, each T1-weighted volume was corrected for intensity non-uniformity and skull-stripped. The extracted brain was spatially normalized to MNI standard space through nonlinear registration using ANTs. Cortical surfaces were reconstructed using FreeSurfer. Functional data was motion corrected using mcflirt (FSL), followed by co-registration to the corresponding T1-weighted image using boundary-based registration with six degrees of freedom. Motion correcting transformations, BOLD-to-T1w transformation and T1w-to-MNI warp were concatenated and applied in a single step using antsApplyTransforms (ANTs).

Confounds including the 24 head motion parameters (six basic motion parameters + six temporal derivatives + 12 quadratic terms and their six temporal derivatives), mean white matter and CSF signals were computed and regressed from the preprocessed fMRI data for each individual. The residuals of this regression were then subjected to spatial smoothing with a Gaussian smoothing kernel of 6mm FWHM and Wishart filtering (see Methods).
